# Zero-shot design of drug-binding proteins via neural selection-expansion

**DOI:** 10.1101/2025.04.22.649862

**Authors:** Benjamin Fry, Kaia Slaw, Nicholas F. Polizzi

**Affiliations:** Harvard Graduate Program in Biophysics, Harvard University, Boston, MA, 02215, USA; Department of Cancer Biology, Dana-Farber Cancer Institute, Boston, MA 02215, USA; Department of Biological Chemistry and Molecular Pharmacology, Harvard Medical School, Boston, MA 02215, USA

## Abstract

Computational design of molecular recognition remains challenging despite advances in deep learning^1–3^. The design of proteins that bind to small molecules has been particularly difficult because it requires simultaneous optimization of protein sequence, protein structure, and ligand conformation^1–7^. Despite their promise, current deep-learning algorithms have struggled to navigate this landscape, precluding the zero- or few-shot design of binders. Here we show that the combination of two neural networks in an iterative design algorithm can create small-molecule binding proteins from scratch with high accuracy. To optimize a design in the joint distribution of sequence, structure, and ligand conformation, we use a pair of neural networks that were trained on reciprocal tasks. We train and use a graph neural network, LASErMPNN, to design protein sequence given protein−ligand co-structure, and we use RoseTTAFold-All Atom^8^ (RFAA) to predict protein−ligand co-structure given protein sequence. We iteratively apply these two networks to design proteins that bind the drug, exatecan, a topoisomerase I inhibitor that is prone to inactivation by hydrolysis^9^. Each of four experimentally tested designs bound the drug, with the lowest dissociation constant (*K*_d_) near 100 nM. The hit rate and highest affinity design each surpassed the current state-of-the-art method by 5- and 70-fold, respectively. We further show that LASErMPNN can improve upon its own designs in a manner resembling chain-of-thought reasoning. Without experimental input, LASErMPNN suggested two mutations that increased affinity by over two orders of magnitude (*K*_d_ = 1.2 ± 0.2 nM). Designs were selective, structurally accurate, and achieved their intended purpose to protect the drug from hydrolysis. Our work describes a recipe for using neural networks to automate the design of high affinity small-molecule binding proteins, which should have wide application in the creation of novel drug-delivery vehicles, antidotes, sensors, and enzymes.

## Main text

The de novo design of small-molecule binding proteins remains a significant challenge^1–4,8,10–14^, despite rapid progress elsewhere in the field^15–24^. The high-dimensional search space—including protein sequence, protein structure, and ligand conformation—is difficult to navigate by computation alone. An algorithm that could efficiently explore this space would enable the rapid development of novel antidotes, sensors, or delivery vehicles. Deep neural networks are poised to fill this role but so far have only modestly contributed to the design of small-molecule binders^1,2,5,14,25^.

There have been sparks of success in the design of small-molecule binding proteins^1,2,4–6,14,26–28^, but these cases have often relied on high-throughput experimental selection and optimization. In the few examples where computational hit rates were high (33%)^5,6^, interactions with ligands were modeled by approximating their functional groups as parts of amino acids. To generalize binder design, neural networks can, in principle, learn directly from training data to predict protein sequence and protein–ligand co-structure.

Self-consistency has been a guiding principle for protein design since the compelling demonstration of a novel topology^29^. The idea is that the designed sequence—computed from a static input backbone—should give rise to a predicted structure that closely resembles the input backbone (Fig. 1a). This principle has been successfully implemented using modern neural networks, such as ProteinMPNN^30^ for sequence design and AlphaFold2^31^ (AF2) for structure prediction, to design protein sequences that fold to distinct shapes and that bind to peptides and protein targets^15,16,19,32^. However, self-consistency has been difficult to compute for small-molecule binding proteins because these networks cannot represent non-amino-acid-based chemistry. Now, neural networks like RFAA^8^, Boltz-1^33^, and AlphaFold3^34^ (AF3) can predict protein–ligand co-structure from a protein sequence and a ligand SMILES string. Using such models, we can now require that a self-consistent design also has a ligand predicted to bind in the same site as the input structure used to design the sequence (Fig. 1a).

**Fig. 1.**
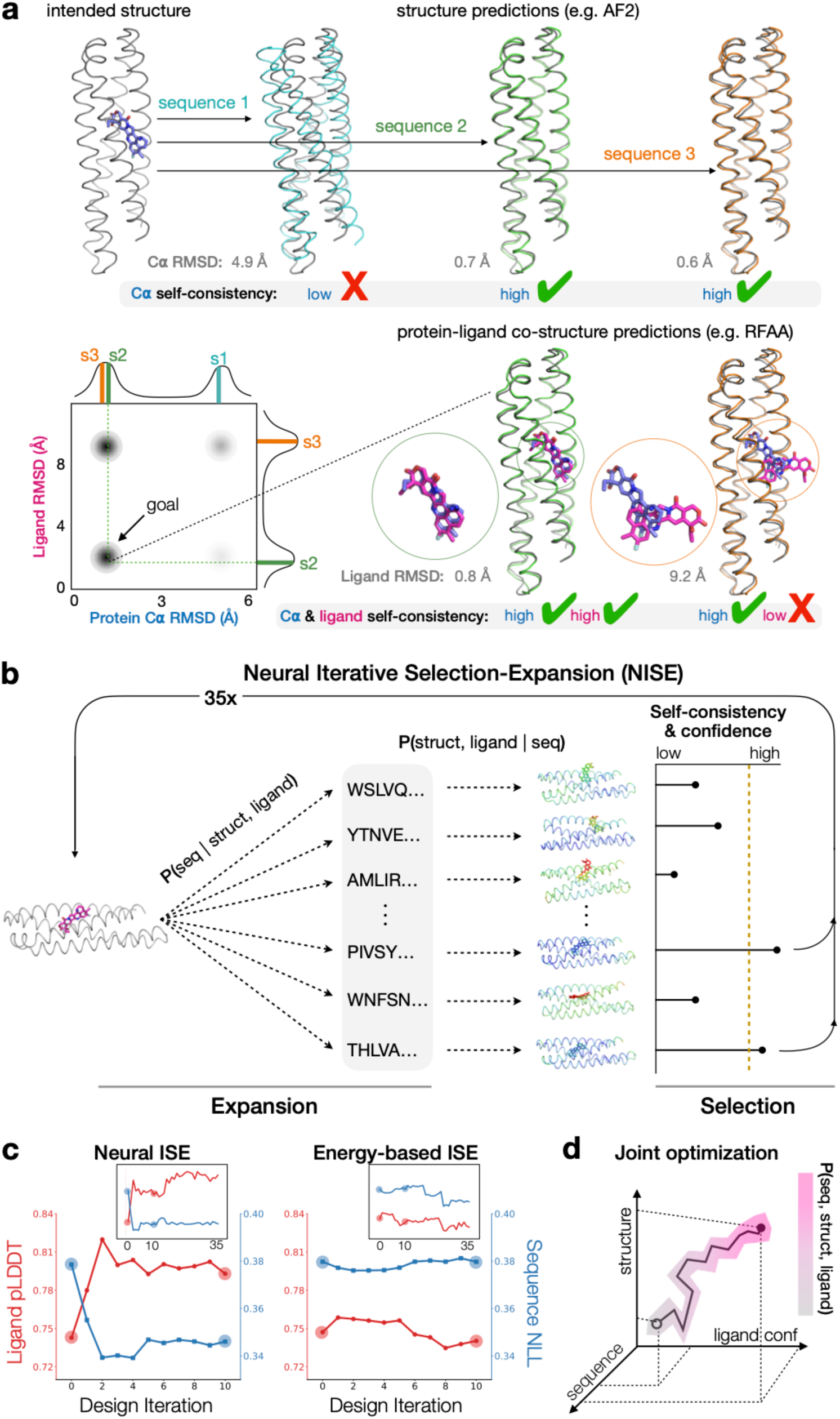
A self-consistency optimization algorithm for the design of small-molecule binding proteins. **a**, The classical principal of self-consistency in protein design seeks a sequence that gives rise to a predicted protein structure (for example, with AlphaFold2, AF2) that closely resembles the intended protein structure (low alpha-carbon root mean square deviation, C⍺ RMSD). Neural networks that predict protein–ligand co-structure, such as RoseTTAFold-All Atom (RFAA), can now extend this concept into the ligand dimension, by additionally seeking that the predicted ligand location (magenta) closely agrees with the intended ligand location (blue), as measured, e.g., by heavy-atom RMSD. Lower left: A two-dimensional illustrative plot of protein–ligand self-consistency, with hypothetical marginal distributions shown above the axes. The design objective is to compute sequences that have predicted co-structures that populate the lower left quadrant (low C⍺ RMSD and low ligand RMSD, green structure). Note that predicted structures with low C⍺ RMSD can have high ligand RMSD (orange structure), suggesting that the designed binding site might not sufficiently encode specific binding. **b**, Schematic of our neural iterative selection–expansion protocol (NISE), which uses an initial protein structure (backbone coordinates only) and ligand location (magenta) to conditionally design protein sequences (expansion), using protein–ligand co-structure prediction not only to filter by protein–ligand self-consistency and rank by model confidence (e.g. predicted local distance difference test, pLDDT, of ligand), but also to be used as direct input into the next round of sequence design. The depicted proteins and ligands are colored by model confidence (red-yellow-green-cyan-blue, low to high). The yellow dashed line denotes a cutoff for selecting protein–ligand co-structures for input into the sequence predictor (backbone and ligand only) during the next round of design. In this work, NISE proceeds for 35 cycles of expansion and selection. The probability distributions used to create sequences and predict co-structures can be approximated by trained neural networks such as LASErMPNN (vide infra) and RFAA. P(seq | struct, ligand) denotes probability of protein sequence (seq) given protein structure (struct) and ligand conformation (ligand), and similarly for P(struct, ligand | seq). **c**, Comparison of iterative selection–expansion (ISE) protocols for neural *vs* energy-based approaches to generating modified protein–ligand coordinates. Energy-based ISE is an ISE protocol that replaces P(struct, ligand | seq) with a traditional structure optimization algorithm, Rosetta energy minimization. In the energy-based protocol, we select designs based on low energy of the ligand after structural optimization under the Rosetta energy function. After proceeding through 35 ISE iterations, we predict the structures of all designed sequences using RFAA and plot the ligand pLDDT (red, third quartile) *vs* the design iteration (inset shows up to 35 iterations). In both cases, we use LASErMPNN to compute the protein sequences and plot the negative log likelihood (NLL) of the designed sequences (blue, first quartile) under the probability distribution of the model, P(seq | struct, ligand). A lower NLL is a more optimal sequence under the model. We see that NISE (but not energy-based ISE) simultaneously optimizes both ligand confidence (higher pLDDT) and protein sequence quality (lower NLL). These plots were created using the starting point to NISE shown in Fig. 3c, using exatecan as the ligand. See supplement for additional details. **d**, As shown in **c** by the increase in pLDDT and decrease in NLL, simultaneous optimization of samples from two reciprocal conditional probability distributions, P(seq | struct, ligand) and P(struct, ligand | seq), strongly suggests that NISE is optimizing on the joint probability distribution of sequence, structure, and ligand conformation, P(seq, struct, ligand), shown schematically here.

Here, we design binders by implementing a self-consistency optimization algorithm that explicitly considers small-molecule ligands. To do this, we use a pair of reciprocal ligand-aware neural networks. We train a graph neural network, LASErMPNN, to design protein sequence given a protein backbone and a docked ligand; in tandem, we use RFAA to predict protein−ligand co-structure given a protein sequence and ligand SMILES string. These networks are used in an iterative loop that refines both sequence, structure, and ligand conformation.

## NISE sampling algorithm

Because initial sequence−structure pairs are likely not optimal for both folding and binding, we need a way to perform local refinement in the joint space of sequence, structure, and ligand conformation. To maximize self-consistency, we iteratively perform sequence design and co-structure prediction, using coordinates from predicted protein–ligand co-structures as new inputs for sequence design (Fig. 1b). To avoid getting trapped in local minima, we encourage exploration by sampling many sequences for a given backbone, drawing from the probability distribution of LASErMPNN (see below) at elevated temperature. For each sequence, we compute a protein−ligand co-structure, filtering out those with low self-consistency with respect to both ligand and backbone coordinates. We use structures with the most confidently predicted ligands as new inputs (both backbone and ligand atoms) for sequence design. Throughout this process, which we call NISE (neural iterative selection–expansion), we avoid use of energy functions and instead rely on model confidence to guide the optimization (Fig. 1b,c).

This approach is similar to iterative coordinate ascent, where alternating argmax sampling from two conditional distributions, p(b|a) and p(b|a), locally climbs a high probability mode in the joint distribution, p(a,b)^35^. By using neural networks to sample from the conditional probabilities p(sequence | structure, ligand conformation) and p(structure, ligand conformation | sequence), we optimize toward a high probability mode in the distribution p(sequence, structure, ligand conformation) that is representative of the underlying training data, here consisting of protein– ligand co-structures in the protein databank (PDB, Fig. 1d).

We use NISE—implemented with LASErMPNN and RFAA as the joint reciprocal networks—to design proteins that bind the small-molecule drug, exatecan, a topoisomerase I inhibitor. This general algorithm harnesses the power of neural networks to unlock small-molecule binder design, with hit rates in this work of 100% (*K*_d_ of 120 nM - 17 µM). We further use LASErMPNN to drive affinity maturation of its own designed sequence, in a process we call neural proofreading, resulting in a 100-fold improvement in affinity (*K*_d_ = 1.2 ± 0.2 nM) after installing just two predicted mutations.

## LASErMPNN neural network

LASErMPNN (ligand-aware sequence engineering message-passing neural network) is a heterograph neural network trained on protein–ligand co-crystal structures in the PDB to predict protein sequence and sidechain dihedral angles given protein backbone coordinates and ligand atomic coordinates (Fig. 2a). During inference, LASErMPNN autoregressively decodes each residue of an input protein–ligand complex using a randomly chosen decoding order. We trained the model until convergence (accuracy of train and validation sets overlapped) and tuned its hyperparameters to maximize foldability of designed sequences without significantly sacrificing sequence recovery (Fig. E1). The model accurately recovers native protein sequence on a held-out test set, outperforms a similarly trained ligand-free version of itself, and is comparable to (slightly outperforms) a retrained version of a similar model, LigandMPNN^25^ (Fig. 2b).

**Fig. 2.**
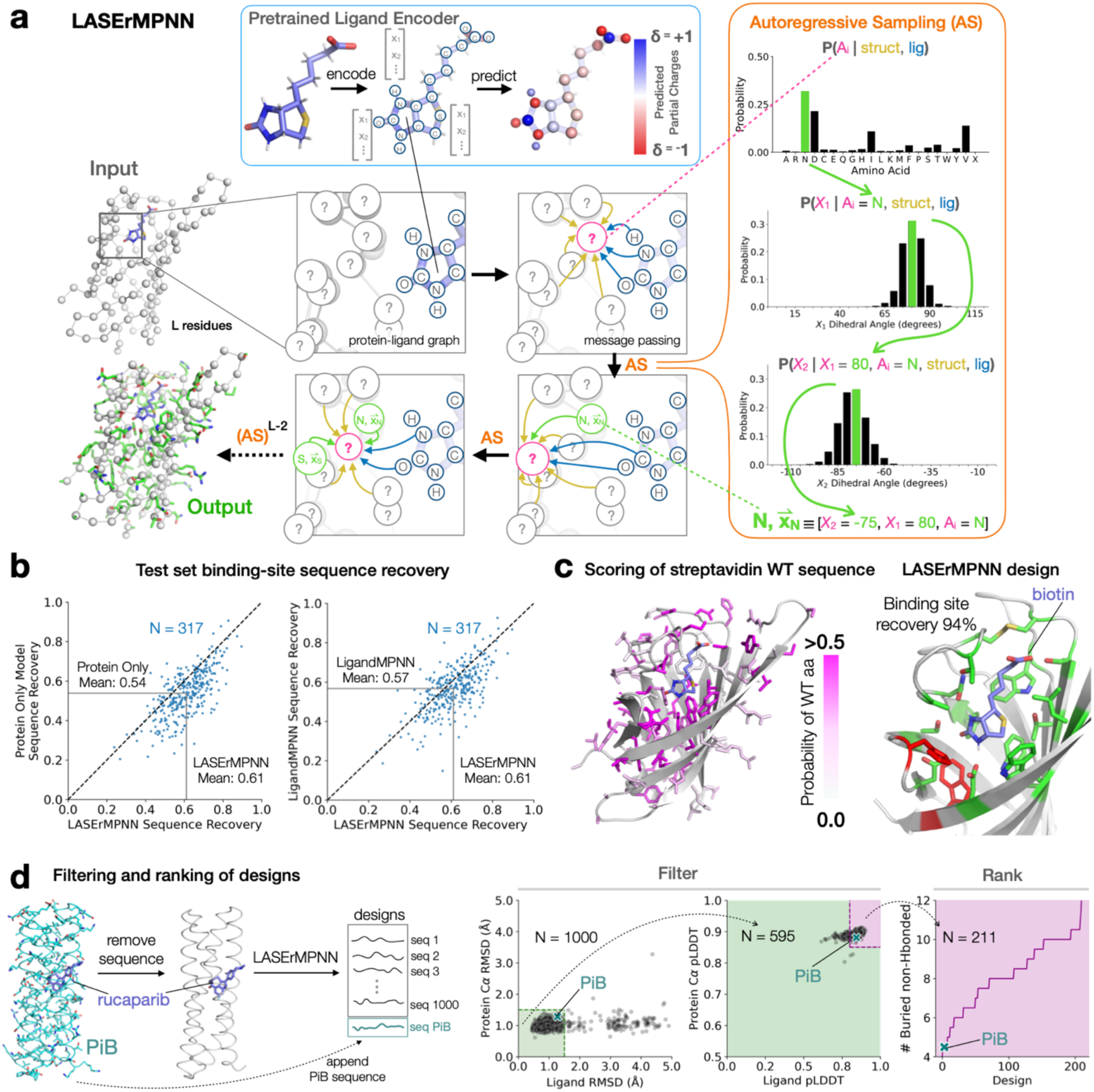
LASErMPNN designs protein sequence conditioned on protein–ligand co-structure. **a**, The LASErMPNN neural network is an encoder-decoder network that takes a protein–ligand co-structure as input (protein backbone atoms and ligand atoms) and is trained to autoregressively predict (orange box) the amino-acid identity and sidechain dihedral angles of each protein residue. LASErMPNN forms a heterograph from the protein–ligand co-structure that encodes the protein and ligand nodes separately. The ligand node embeddings are taken from a pretrained ligand encoder (blue box) and are frozen during training. The ligand encoder is trained to predict atomic properties of ligands (e.g. partial atomic charge) conditioned on atomic coordinates and elements. Autoregressive sampling is performed for each of *L* residues of the protein in turn, with each new residue prediction also conditioned on previously decoded residue identities and their associated dihedral angles (green). LASErMPNN outputs a 3-dimensional structure of the sampled sidechain coordinates built from the predicted residue identities (A_i_) and sidechain dihedral angles (X_1_, X_2_, X_3_, and X_4_). **b**, LASErMPNN performance on a held-out test set of 317 proteins *vs* a similarly trained protein-only model (left, all ligand information removed) and *vs* a retrained LigandMPNN model (right). These plots show the prediction accuracy over the binding-site residues when the model is tasked with predicting the entire protein sequence of each test protein (argmax sampling). Note that overall sequence recovery also shows similar trends (see supplement). **c**, LASErMPNN favorably scores the wildtype (WT) sequence of monomeric streptavidin, with high-probability amino acids (aa), shown in magenta, located in the biotin binding site and protein core. Using the protein backbone and biotin as input, we designed 10,000 sequences of monomeric streptavidin with LASErMPNN. The design with the highest binding-site sequence recovery (94%, right) is shown with accurately predicted residues in green and two designed mutations in red. This design appeared in the top 25 designs when ranked by buried, non-H-bonded polar atoms. **d**, Sequences designed by LASErMPNN can be ranked to prioritize high affinity binders. Here we use the X-ray crystal structure (pdb accession 8tn6, chain A) of a previously de novo designed rucaparib-binding protein (PiB, cyan) as input to LASErMPNN. We design 1000 sequences for the protein–ligand co-structure and append the previously designed PiB sequence to the set. We then predict the protein–ligand co-structures for each sequence using RFAA. We filter these designs by *i*) the ligand heavy-atom root mean square deviation (RMSD), *ii*) protein C⍺ RMSD, *iii*) ligand pLDDT, *iv*) protein C⍺ pLDDT, and *v*) sidechain clashes with the ligand after repacking in the absence of ligand (not shown, see supplement). The remaining designs are then ranked by number of buried, polar atoms that are not H-bonded (right). Note that the filtering and ranking protocol ranks the PiB sequence 4^th^ out of 1001, showing it enriches for sequences with high affinity (*K*_d_ of PiB and rucaparib is < 5 nM).

Key differences for LASErMPNN with respect to LigandMPNN include *i*) the presence of a distinct, pretrainable ligand encoder, *ii*) simultaneous decoding of sidechain dihedral angles with amino-acid identity, and *iii*) the inclusion of the ligand nodes in each round of encoding and decoding. Ablation studies showed that performance depended on simultaneous prediction of sidechain dihedral angles during sequence decoding and on pretraining a ligand encoder module on a large set of synthetic ligands^36,37^, with the training task to predict atom-level properties derived by quantum chemical computations, such as partial charge (Table E1).

As a strict in silico test case, we rigorously held out the streptavidin fold from the training set via sequence, structural, and evolutionary similarity. The streptavidin–biotin complex is one of the strongest known non-covalent complexes in nature; recovery of the sequence would be a convincing indication that LASErMPNN is performing as intended. LASErMPNN favorably scores the native sequence of monomeric streptavidin, displaying high probabilities of native core- and binding-site residues (Fig. 2c). Designed sequences for monomeric streptavidin with bound biotin showed high sequence recovery of the binding-site residues (average of 53% and 38% with and without biotin, respectively.) (Fig. E2). Although RFAA struggled to predict self-consistent structures for this β-barrel topology, many of the designed sequences were predicted by Boltz-1 to fold without use of multiple sequence alignments (65% success rate for Boltz-1 *vs* 0.4% for RFAA; protein C⍺ and ligand heavy-atom RMSD < 2.5 Å). We ranked self-consistent structures by a formula that penalizes buried, non-hydrogen-bonded (non-H-bonded), polar atoms and saw an inverse correlation between this metric and binding-site sequence recovery (Fig. E3). The top 25 designs showed up to 94% binding-site sequence recovery (out of 10,000 total designs, Fig. 2c). Thus, LASErMPNN can rapidly sample binding-site sequences that took millions of years to evolve.

We tested the same filtering and ranking approach on a previously de novo designed drug-binding protein^5^, PiB, a four-helix bundle that tightly binds the drug rucaparib, an inhibitor of poly-ADP ribose polymerase (Fig. 2d and Fig. E4). The structure of PiB was held out from the training sets of both LASErMPNN and RFAA, so it represents a benchmark closely aligned with our design objective. We used LASErMPNN to design 1000 sequences for the PiB backbone bound to rucaparib. Most designed sequences had self-consistent co-structures predicted by RFAA. We filtered RFAA-folded designs by self-consistency (protein C⍺ and ligand RMSD) and ligand confidence (pLDDT) from RFAA, then ranked the remaining designs by the same formula that penalizes buried, non-H-bonded, polar atoms. The top-ranked design had a binding-site sequence recovery of 74% (differing mainly at surface residues) and a recovery of 80% over all core residues, giving us confidence that the design method and ranking scheme could discover and prioritize high quality de novo binders. The PiB sequence, when assessed using the same metrics, was ranked near the top of the designed sequences (4th out of 1001).

## Design objective

Exatecan is a member of the camptothecin class of clinically approved small-molecule anticancer therapeutics, binding to and inhibiting the DNA–protein complex of topoisomerase I^9^ (Fig. 3a). It is one of the most soluble and potent members of the camptothecins (IC_50_ ∼ 1 nM) and is therefore popular as a payload in antibody drug conjugates (ADCs)^38,39^. The lactone ring of exatecan, characteristic of this class of drug, is prone to hydrolysis at physiological pH, and the free drug rapidly hydrolyzes (half-life about 2 hours) in plasma^9^ (Fig. 3a and Fig. 6a). The hydrolyzed, ring-open, carboxylate form of the drug is significantly less bioactive, due in part to its lower cell permeability and that this form is bound predominantly by serum albumin^9,39^. We sought to design a small protein that could encapsulate the labile lactone ring of the drug, thereby protecting it from hydrolysis (see Fig. 3b,c and Fig. 6c). A designed binder could be used ultimately as a delivery vehicle for the bioactive form of the drug or as a sponge for free drug that has been prematurely liberated from ADCs (although this is outside the scope of the present work). There are no structures of exatecan in the PDB or the Cambridge structural database (CSD), and few structures of related compounds, making this a challenging target for binder design with neural networks.

**Fig. 3.**
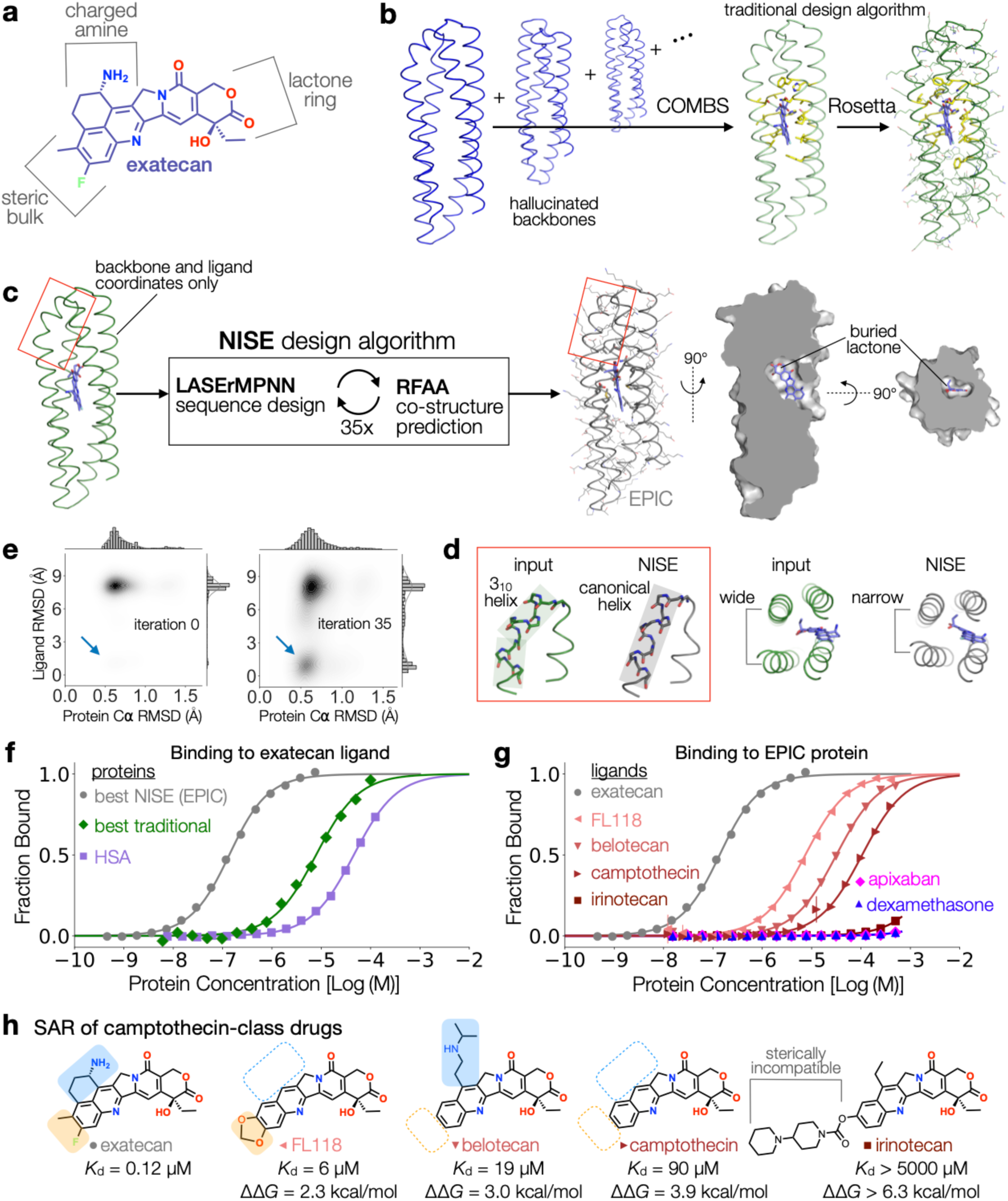
Design of an exatecan-binding protein using neural iterative selection–expansion (NISE). **a**, Chemical structure of exatecan. **b**, An example of a traditional design algorithm using COMBS and Rosetta to design an exatecan-binding protein. Starting from an ensemble of computationally generated four-helix bundles (colored by AF2 C⍺ pLDDT; blue > 90; note that all residues have high pLDDT), COMBS uses van der Mers (yellow) to dock exatecan (blue) within the bundle. A flexible-backbone design protocol using Rosetta then designs the remainder of the sequence. **c**, The NISE design algorithm takes backbone and ligand coordinates as input. Here, we strip the sequence of the traditional design in **c** and use this as input to NISE. We chose four designs from NISE. Shown explicitly (gray) is the design model of the protein, EPIC (exatecan–protein interaction construct). Like the traditional design, the NISE designs bury the lactone ring of the exatecan. **d**, Structural refinements of the design model as a result of the NISE protocol (gray), with respect to the input structure (green). A 3_10_ helix (green) near the ligand (red box in **d**) is converted to canonical helix (gray). A wide helix–helix interface (green) is converted to a narrow interface that allows for better helix–helix packing and more protein–ligand contacts. Note also that exatecan (blue) gets drawn deeper into the bundle as a result of NISE. **e**, A two-dimensional density plot of protein-*vs* ligand RMSD for two iterations of the NISE protocol, using the input structure in **c.** The plot shows an increase in protein–ligand self-consistency in iteration 35 *vs* iteration 0 (blue arrow). Marginal distributions are shown above the axes (*N* = 4000). The design objective—to compute sequences that have predicted co-structures that populate the lower left quadrant (low C⍺ RMSD and low ligand heavy-atom RMSD)—is achieved by NISE. **f**, Fluorescence anisotropy experiments, here converted to fraction of the bound exatecan (fits shown as solid lines), for several different proteins: the highest affinity NISE design (EPIC, gray, *K*_d_ = 0.12 ± 0.03 µM), the highest affinity traditional design (green, *K*_d_ = 8 ± 0.7 µM), and human serum albumin (HSA, light purple, *K*_d_ = 43 ± 3 µM). The traditional design that served as input to NISE after removing the sequence formed a higher order oligomer (see supplement). The concentration of exatecan was 50 nM for each experiment. **g**, Fluorescence anisotropy experiments, here converted to fraction of the bound ligand (fits shown as solid lines), for several different ligands with the EPIC protein. The concentration of each ligand was 50 nM for each experiment. Irinotecan showed only weak binding at the highest protein concentrations, precluding an accurate fit. Dexamethasone and apixaban (conjugated with fluorescein isothiocyanate (FITC) fluorophore) showed no appreciable binding. **h**, Structure–activity relationship (SAR) for the series of camptothecin-class molecules shown in **g**. Change in binding free energy (ΔΔG) is computed relative to the affinity of exatecan–EPIC complex (leftmost).

As initial scaffolds for docking the ligand, we performed family-wide hallucination of single-chain four-helix bundles (see supplement). Briefly, we created four-helix coiled coils using parametric equations^6,40^, generated loops via RFdiffusion^15^ (with loop lengths determined by structural bioinformatics^41^), designed sequences for each using ProteinMPNN, and predicted their structures with AF2. The resulting focused library consisted of forty AF2-predicted four-helix bundles with very high confidence (C⍺ pLDDT). Each structure had at least eight different sequences that passed confidence thresholds, making each sufficiently designable as a scaffold for small-molecule binder design.

We built a few exatecan conformers using an X-ray crystal structure of a camptothecin derivative from CSD as a starting point, then further optimized the rebuilt coordinates using a molecular mechanics forcefield. We docked these conformers into each of our forty backbone templates using a previously described method, COMBS^5,6^, and filtered docked ligands by burial within the protein. These docked ligands underwent a traditional design process using COMBS and van der Mers^5,6^ to design the binding site and then using Rosetta^42^ to compute the remainder of the sequence (with a flexible backbone algorithm) (Fig. 3b). Final designs passed metrics for ordering, such as self-consistency via co-structure prediction with RFAA (low backbone, binding-site residue, and ligand RMSDs). One of these design models, chosen based on a combination of self-consistency and confidence metrics (see below and supplement), served as initial input for our NISE design protocol (Fig. 3c).

## Design of exatecan binders using NISE

We used LASErMPNN and RFAA as the reciprocal networks within the NISE algorithm to design exatecan binders, starting from a docked computational model from COMBS (Fig. 3c). For the initial model given to NISE, we chose from 16 diverse designs produced by a protocol using COMBS and Rosetta (see supplemental methods). We ranked the models by structural self-consistency using a linear combination of ligand RMSD, binding-site residue C⍺ RMSD, and overall backbone C⍺ RMSD. The sequence of the top-ranked design was discarded, and only the backbone and ligand coordinates were used as input to NISE. No experimental characterization of the designs was performed prior to this choice.

In the selection–expansion protocol, we apply a self-consistency filter by retaining only those RFAA-predicted structures with low RMSD (C⍺ and ligand atoms) to the LASErMPNN input from the previous round (Fig. 1b). We avoid ligand drift by requiring that the ligand remains buried in the bundle. We optimize the designs by selecting only the top three self-consistent structures by ligand confidence (pLDDT) each round, then expanding these in the next round by sampling a thousand sequences each (designing the entire sequence with LASErMPNN each time).

During the NISE protocol, we noticed that, while most designed sequences were predicted by RFAA to be self-consistent with respect to C⍺ coordinates, far fewer were predicted to bind the ligand in the correct orientation, though most had an accurately predicted center of mass (Fig. E5). More ligands were predicted to bind in the correct orientation at later design cycles, but these remained the minority compared to alternative binding modes that tended to expose most of the polar atoms of exatecan to solvent (Fig. 3e). These results strongly suggest that self-consistency with respect to the ligand is a much more stringent computational screen for binding than self-consistency with respect to backbone alone, although backbone-only self-consistency has been used with some success recently^14^.

While we only explicitly select for self-consistency and ligand confidence, we observe that NISE also optimizes the confidence of designed sequences under the LASErMPNN model (i.e. the negative log-likelihood, NLL, decreases) (Fig. 1c). Thus, by iteratively optimizing the reciprocal conditionals, NISE climbs the nearest mode in the joint distribution of p(sequence, structure, ligand conformation) (Fig. 1d). This contrasts with a similar iterative optimization scheme using Rosetta energy minimization to sample new backbone- and ligand coordinates, with top structures selected by ligand energy (Fig. 1c and Fig. E6). Iteratively selecting Rosetta-minimized structures and expanding via sequence design by LASErMPNN does not reduce the NLL of designed sequences nor does it increase the pLDDT of ligands in predicted co-structures, implying that these structures are not sufficiently altered to more strongly encode optimal sequences.

After pooling and filtering the designs by self-consistency (Figs. E7 and E8), we selected four NISE designs with the least number of buried, non-H-bonded, polar atoms. The NISE algorithm altered the exatecan binders from the original input in several ways (Fig. 3d). First, as expected, the sequences were overwhelmingly changed relative to the input design from COMBS and Rosetta, including most binding-site residues (Fig. E9). Second, a segment of 3_10_ helix near the binding site was remodeled into more ideal coiled coil. Third, the gap in a wide helix–helix interface was narrowed to allow for better interhelical sidechain packing and increased supercoiling of the bundle. This global structural rearrangement, which reduced the superhelical radius from 7.5 Å to ∼7.2 Å, allowed for more designed contacts to the ligand. Fourth, the ligand was translated and rotated by several Å into the bundle interior, increasing its buried hydrophobic surface area. The intended binding mode was maintained, with the labile lactone ring buried in the core of the protein, although some ligand torsional angles became distorted by RFAA. The four designs, while maintaining the same binding mode, sampled various degrees of ligand burial, polarity of the binding pocket, and protein supercoiling, differing from each other by an average C⍺ RMSD of 1.2 Å and by an average C⍺ RMSD of 1.3 Å to the input backbone.

## Characterization of designs

We ordered synthetic DNA for each NISE design and expressed the proteins in *E. coli*. Each design expressed in good yield and was monomeric by size-exclusion chromatography (SEC) (Fig. E10). Exatecan is intrinsically fluorescent, so its binding can be tracked via fluorescence polarization (FP) experiments using unlabeled ligand (Fig. 3f,g and Fig. E11). FP experiments showed that four out of four designs bound exatecan (Fig. 3f), with dissociation constants (*K*_d_) ranging from 0.12 ± 0.03 to 17 ± 1 µM (3 of 4 with *K*_d_ < 10 µM). For comparison, a non-specific binder, human serum album (HSA), has an affinity with exatecan of *K*_d_ = 43 ± 3 µM. The highest affinity design, which we call EPIC (exatecan–protein interaction construct), binds approximately 360x more tightly.

We also expressed 16 COMBS designs, representing 8 distinct ligand positions and backbones (i.e. poses). Only three designs bound exatecan, with *K*_d_ values of 8 ± 0.7, 12 ± 2, and 44 ± 7 µM (Figs. 3f and E11). The design used as the input pose to NISE caused an increase in FP when mixed with exatecan, but the data could not be fit to a single-site model, and SEC showed the protein aggregated. Of the set of 16, the three monomeric binders were the top-ranked designs by both self-consistency (ligand RMSD and backbone C⍺ RMSD) and ligand pLDDT, metrics that have previously been shown to correlate with peptide binding using AF2^43^ (Fig. E12). Interestingly, these three models were also top-ranked after applying the same metrics to the mean of five designed sequences from LASErMPNN, suggesting that the backbone- and ligand coordinates were particularly designable (Fig. E13).

Because the strongest binder was similar to the input pose for NISE, we sought to more thoroughly explore this pose in the context of our more traditional methods. We therefore ordered two additional designs using the COMBS and Rosetta protocol for this backbone–ligand pair (see supplement). The additional designs, which shared most of the binding-site residues with the parent binder, also bound exatecan, albeit more weakly (*K*_d_ = 15 ± 2 and 43 ± 10 µM, Fig. E11). These results suggest that a designable pose, to some extent, can give rise to a binder using a variety of sequence design methods^1,2^. However, affinities of the COMBS designs were not dramatically different from that of a non-specific binder (HSA).

The substantially higher affinities of designs from NISE *vs* COMBS cannot be attributed to the use of RFAA as a filter for quality, since designs from both methods had self-consistent co-structures using RFAA. Instead, we attribute the tighter binding achieved by NISE to its ability to resculpt the backbone and binding site to make the overall sequence more compatible with both folding and binding (Fig. 3d). Indeed, EPIC’s binding site was almost completely changed compared to the input COMBS model, which contained several residues forming H-bonds with polar atoms of the ligand (Fig. E9). EPIC’s binding site takes a more subtle balance between shape-complementary apolar packing and engagement with ligand polar groups; only one (Asp132) of the original H-bonding residues was preserved.

The high affinity EPIC contrasted with the lower affinity NISE designs in several ways. EPIC had low sequence identity to other designs (ranging from 34-57%; 36-81% for core residues), and it buried more ligand apolar surface area than the weakest affinity NISE binder (*K*_d_ = 17 ± 1 µM). EPIC’s binding site contained the least number of polar residues, although it shared with other designs a buried glutamine at position 51 (near the lactone of exatecan) and an aspartate at position 132, which interacts with exatecan’s positively charged amine.

EPIC is highly thermostable (Fig. E14) and specific for the camptothecin class of drugs, with affinity proportional to the steric and chemical similarity to exatecan (Fig. 3g,h and Fig. E11). FP experiments showed binding to FL118, belotecan, and camptothecin, in order of increasing dissociation constants (*K*_d_ of 6 ± 1, 19 ± 5, and 90 ± 21 µM, respectively). These molecules show a clear structure–activity relationship for the fluorophenyl ring substituents and the amine of exatecan. While elimination of either functional group is detrimental to binding, desolvation and packing of the fluoro and methyl groups (ΔΔG = 3.0 kcal/mol) appear to drive affinity more so than the interaction with the amine (ΔΔG = 2.3 kcal/mol). Elimination of both the amine and the ring substituents in camptothecin is partially additive with respect to binding-energy destabilization (ΔΔG = 3.9 kcal/mol). EPIC showed no appreciable binding to irinotecan, which contains a bulky prodrug substituent that is sterically incompatible with the designed binding pocket. As expected for a well-sculpted pocket, EPIC does not bind drugs of other classes, such as anticoagulants (apixaban) and steroids (dexamethasone).

## Neural proofreading of EPIC sequence

We wondered if we could improve the affinity of EPIC for exatecan via computation alone. Some strategies might employ an additional round of backbone diversification around the computational model followed by ligand docking and several rounds of sequence design to create and test thousands of new sequences^1,2^. Instead, we reasoned that LASErMPNN could refine the EPIC sequence in a more focused approach, understanding that even a single amino-acid change can contribute significantly to affinity. We therefore used LASErMPNN to “proofread” the EPIC sequence by suggesting single amino-acid mutations in the binding site that decrease its NLL (Fig. 4).

**Fig. 4.**
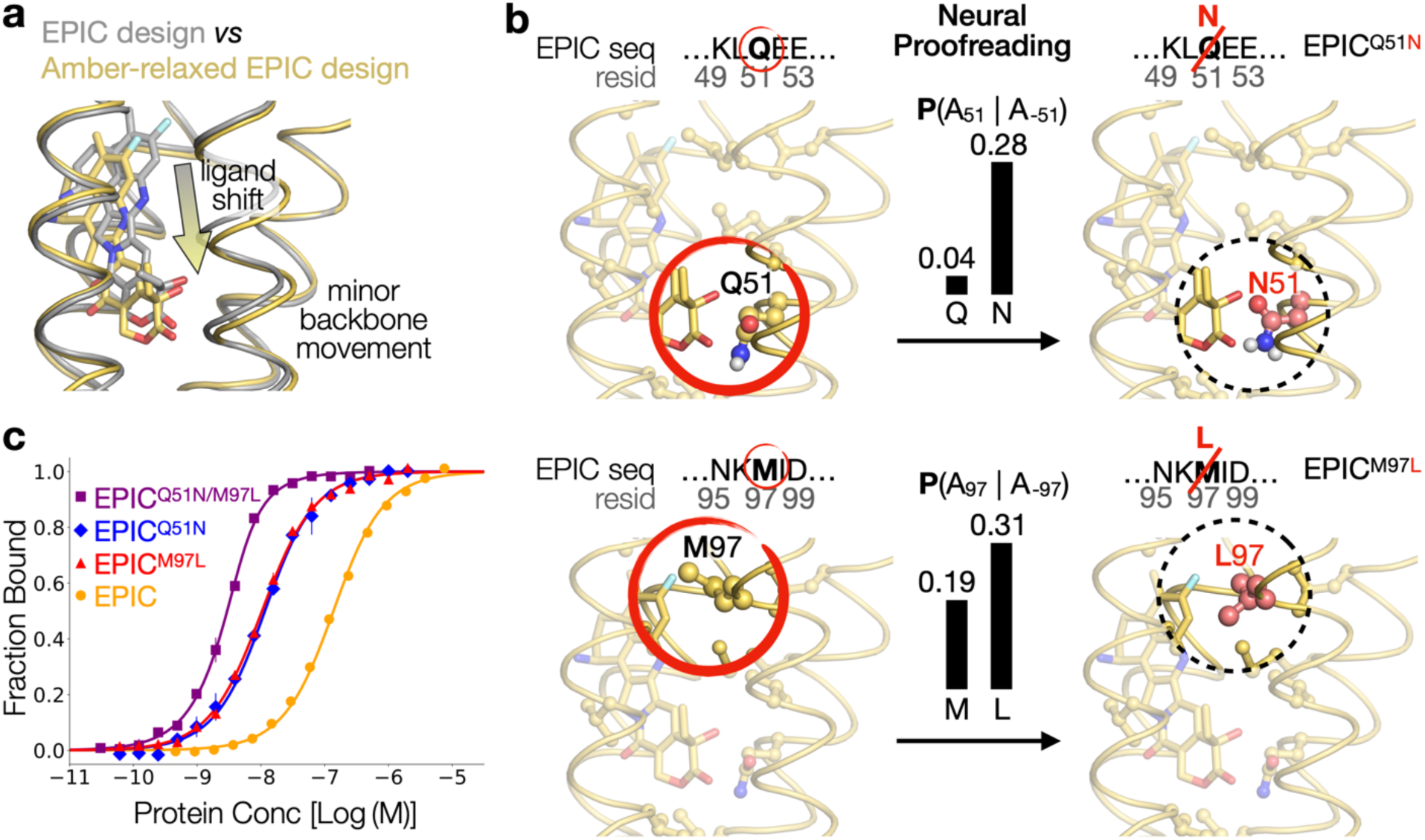
Neural proofreading of EPIC by LASErMPNN increases affinity to exatecan. **a**, Computational model of EPIC (gray, RFAA prediction) superimposed onto the same model after coordinate relaxation using the Amber molecular mechanics energy function (yellow). As a result of energy minimization, exatecan shifts deeper into the binding pocket, and the backbone around the ligand moved slightly. These new backbone and ligand coordinates were input into LASErMPNN for neural proofreading. **b**, Neural proofreading of EPIC using LASErMPNN computes the probability of an amino acid (A_i_) given the entire context of the remainder of the designed sequence (A_-i_). Explicitly shown are examples of neural proofreading for residue 51 (top) and residue 97. For each, LASErMPNN is given the ligand coordinates, backbone coordinates, and amino-acid identities of the other 147 residues of the EPIC sequence (148 residues total), using the Amber-relaxed design. With this information, LASErMPNN computes the probability profile for residue 51, P(A_51_ | A_-51_), and, separately, for residue 97, P(A_97_|A_-97_), while also computing the dihedral angles of each residue (i.e. repacking the protein). In each case, the original residue in the EPIC sequence (Q and M, respectively) is lower probability than another residue (N and L, respectively). **c**, The single amino-acid mutations show higher binding affinity than EPIC as measured by fluorescence anisotropy experiments (Here, converted to fraction of ligand bound after performing the fitting; fits shown as solid lines.). The double mutant, EPIC^Q51N/M97L^, shows two orders of magnitude higher affinity. (Note that the concentration of exatecan in these experiments, 50 nM, was comparable to the *K*_d_ for the mutant proteins, so the quadratic form of the binding equation was used for fitting. Consequently, the *K*_d_, especially for the double mutant, does not appear exactly at the protein concentration corresponding to the half-maximum of fraction bound. See supplement for global fits at various ligand concentrations.)

There are a few reasons to expect that the same model that designed the EPIC sequence can also offer meaningful suggestions for mutations. First, because LASErMPNN designs sequence conditioned on the structure and amino-acid identity/rotamer of previously decoded residues, only the last designed residue has all the context of the full sequence. Like chain-of-thought prompting of large language models^44,45^, the additional context of performing all-but-one conditional design could therefore shift the probability distributions (Fig. E14). Second, we perform sequence design at a high temperature during NISE to encourage diversity. Revisiting each choice in turn allows for error correction due to high-temperature sampling. Third, and perhaps most important, we use different backbone and ligand coordinates as input for the ligand-bound structure (Fig. 4a). These coordinates come from an RFAA-predicted structure of the EPIC sequence, which can also be relaxed in several ways to add subtle structural diversity that can nevertheless influence the model probabilities.

Neural proofreading by LASErMPNN offered several suggestions for mutations to the EPIC binding site, using an Amber-relaxed^46^ model of the RFAA prediction (Fig. 4a,b and Fig. E15). We ordered and individually characterized the single amino-acid mutations, EPIC^Q51N^ and EPIC^M97L^, each of which improved the NLL of EPIC. FP experiments showed each mutant increased affinity for exatecan by over 10-fold (Figs. 4c and E16, *K*_d_ of 8.0 ± 1.6 and 7.4 ± 0.7 nM, respectively; ΔΔG ≅ – 1.6 kcal/mol). We combined these sterically compatible mutations in a double mutant, EPIC^Q51N/M97L^, and found them to be energetically additive, resulting in a 100-fold higher affinity than EPIC for exatecan (*K*_d_ = 1.2 ± 0.2 nM; ΔΔG = – 2.7 kcal/mol). Another mutation, EPIC^G104M^, that was predicted to decrease affinity (increase NLL for ligand-bound structure) while stabilizing the unbound state (decrease NLL for ligand-free structure), was shown to dramatically weaken binding (Fig. E17, *K*_d_ = 57 ± 8 µM). SEC experiments showed that each mutant protein remained monomeric (Fig. E18). These results demonstrate the predictive power of chain-of-thought proofreading for in silico affinity maturation, without use of an experimentally determined structure.

## X-ray crystal structures of EPIC and EPIC^Q51N^

The structure of EPIC was determined to 2.0 Å resolution and displayed clear density for bound exatecan (Fig. 5a,b and Fig. E19). The binding mode of the ligand is as predicted, with some deviation near the δ-lactone, which was expected due to inferior modeling in this region by RFAA (Fig. 5a). Asp132 shows the intended H-bond to the ligand amine. Gln51 receives an H-bond from exatecan’s hydroxyl, but the large sidechain is too sterically constrained to donate an H-bond to the carbonyl of the lactone ring. The backbone agreed closely with the input backbone to LASErMPNN that was used immediately upstream in the NISE design cycle (C⍺ RMSD 0.8 Å), and rotamers of both core and binding-site residues were accurately predicted by LASErMPNN (Fig. 5a). After superposition on binding-site residues (C⍺ within 8 Å of ligand heavy atoms, RMSD 0.4 Å), the ligand coordinates differed from those in the input structure by an average atomic displacement of less than 1 Å and a rotation of 14° about a central axis (Fig. 5a). Thus, the designed structure of EPIC was largely achieved.

**Fig. 5.**
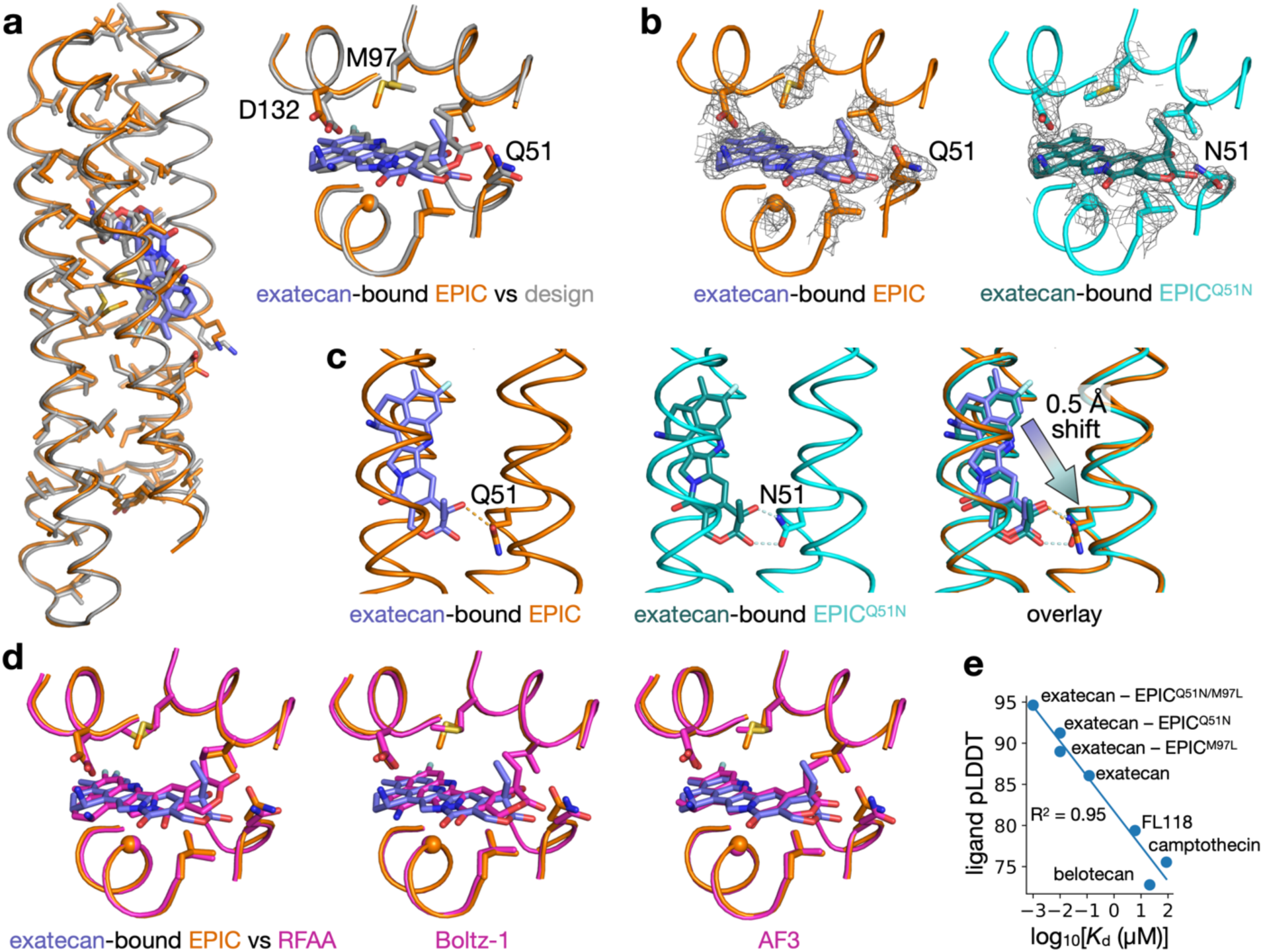
X-ray crystal structures of EPIC agree with the design. **a**, Overlay of EPIC design model (gray) with X-ray crystal structure of the exatecan-bound EPIC complex (orange protein, blue exatecan). The sidechains of core residues are shown, which agree overwhelmingly with the conformations designed by LASErMPNN (gray). The enlarged image of the binding site shows a few key residues, such as Asp132, Met97, and Gln51. **b**, Composite omit map of the 2.0 Å resolution structure of EPIC (orange, left), contoured at 1 σ, shows clear density (gray mesh) for the ligand (blue) and surrounding sidechains. Similarly, the composite omit map of the 2.2 Å resolution structure of EPIC^Q51N^ (cyan, right), contoured at 1 σ, shows clear density (gray mesh) for the ligand (green) and sidechains, including Asn51. **c**, Side view of EPIC (orange) and EPIC^Q51N^ (cyan) highlighting the H-bonding interactions (dashed lines) between exatecan and Gln51 *vs* Asn51. Exatecan moves slightly deeper into the pocket in EPIC^Q51N^ to interact with Asn51 (rightmost image, overlay of both structures). **d**, Comparison of X-ray structure of EPIC (orange protein, blue exatecan) with predicted co-structures (magenta) from RFAA, Boltz-1, and AF3. AF3 predicts the co-structure most accurately of the three. **e**, Plot of measured binding affinities (log_10_[*K*_d_], units of µM) *vs* average pLDDT of the ligand in AF3-predicted co-structures. The ligands were each predicted by AF3 to bind in the designed pocket in a similar orientation. Binding is between the labeled ligand and EPIC unless otherwise noted (for EPIC mutants).

The structure of exatecan-bound EPIC^Q51N^ (2.2 Å resolution) gives insight into the higher affinity of this complex (Fig. 5b,c and Fig. E19). The shorter sidechain of Asn51 draws exatecan slightly more into the pocket (by approximately 0.5 Å), and its carboxamide engages in a bidentate H-bonding interaction with exatecan’s hydroxyl and lactone carbonyl (Fig. 5c). The ligand is therefore more buried and properly desolvated.

The structures of EPIC and EPIC^Q51N^ present an opportunity to assess the performance of modern protein–ligand co-structure predictors. Each of three structure predictors accurately computes the backbone of EPIC; the main differences occur at the ligand (Fig. 5d). While each program generally predicts the correct binding mode of exatecan, they struggle to compute some aspects of the ligand conformation, often producing physically distorted molecules. Boltz-1 and RFAA tend to flatten functional groups into the plane of the extended ring system, e.g. the sp^3^ hybridized amine and the lactone. AF3 predicts the bound complex most accurately but still deviates slightly from the true ligand conformation. Interestingly, the ligand pLDDT of predicted co-structures by AF3 rank-orders the different camptothecin-class drugs by affinity (Fig. 5e). To a lesser extent, ligand pLDDT from Boltz-1 also correlates with binding affinity (R^2^ = 0.61, Fig. E20). While RFAA was not able to predict a consistent binding mode for the other camptothecin-class ligands, it is slightly more confident in the placement of exatecan for EPIC^Q51N/M97L^ *vs* EPIC (Fig. E20).

## EPIC protects exatecan from chemical degradation

EPIC dramatically repartitions the equilibrium distribution of hydrolyzed *vs* non-hydrolyzed exatecan (Fig. 6). At pH 7.4, the lactone ring in camptothecin-class drugs are predominantly found in the hydrolyzed, ring-open, carboxylate form, which is much less bioactive (Fig. 6a,b)^9,39^. Ring-closed exatecan has been shown to rapidly hydrolyze in human plasma, reaching a level of 15% within 10 hours^9^. While the high level of HSA in plasma is expected to bind exatecan, its binding does not prevent hydrolysis (Fig. 6c-e). This is consistent with a camptothecin-bound structure (PDB 4L9K) where the ring-open form of the drug was observed to be stabilized by nearby positively charged Arg residues (Fig. E21). On the contrary, we find that EPIC and its mutants dramatically reduce the amount of hydrolyzed exatecan by sequestering the labile lactone from solvent (Fig. 6d,e). Time-resolved absorption experiments showed that EPIC^Q51N/M97L^ maintains a 99% level of ring-closed exatecan in PBS buffer (pH 7.4) at room temperature for days (No major hydrolysis product was observed at the longest measured time points, up to 50 hr.). The ability of EPIC^Q51N/M97L^ to protect exatecan from hydrolysis is similarly observed in the presence of physiological concentrations (500 µM) of HSA (Figs. E22 and E23). Thus, EPIC can maintain a reservoir of the bioactive form of the drug, which has potential applications in delivery and targeting.

**Fig. 6.**
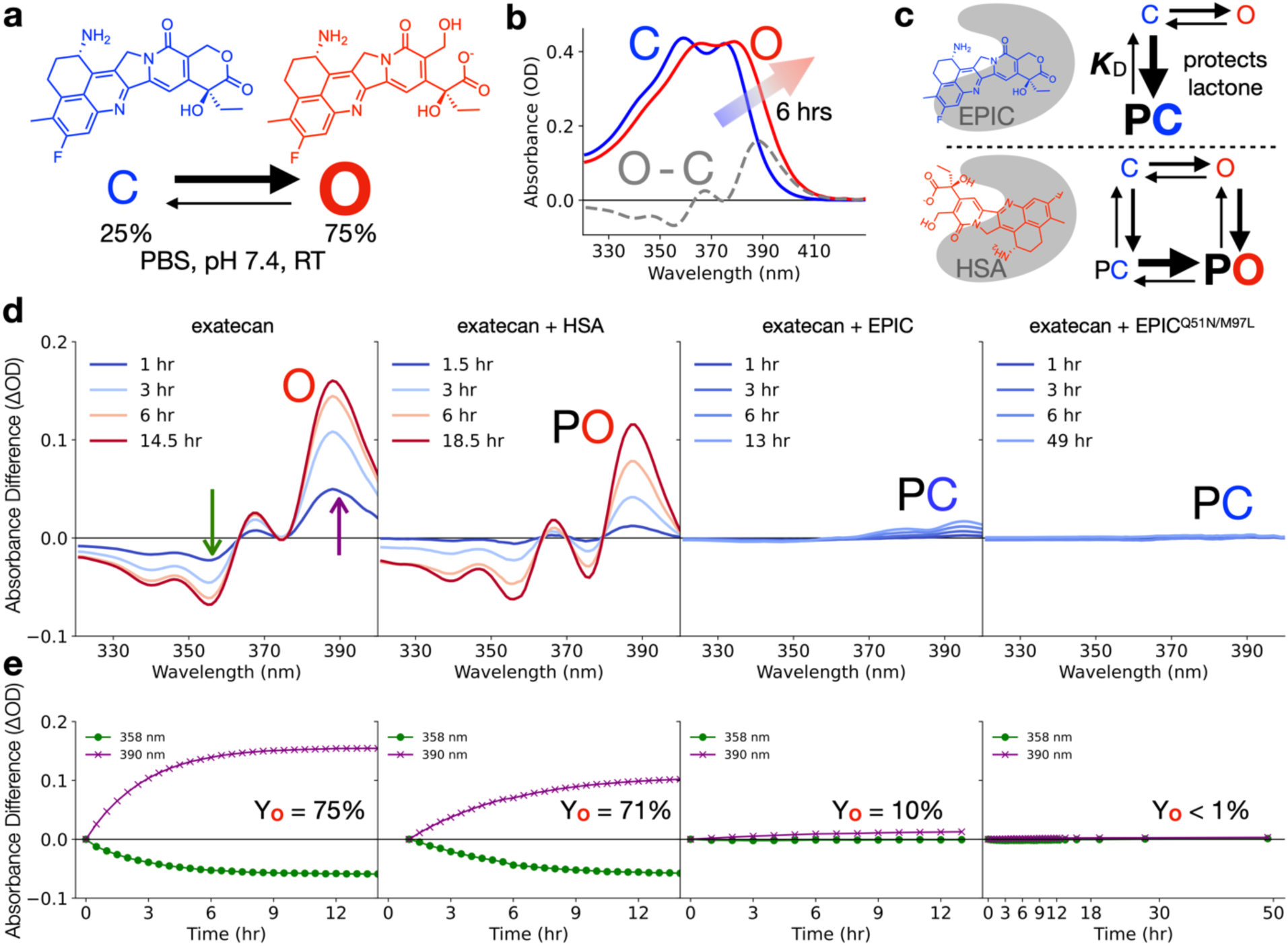
EPIC protects exatecan from hydrolysis. **a**, Exatecan exists in a ring-closed (C) lactone form and ring-open (O) carboxylate form. The O state is not considered to be bioactive. At physiological pH (7.4) and room temperature (RT), the O state is more favorable (equilibrium population 75%). In human plasma at 37 °C, the equilibrium is shifted further to 85% O. **b**, C *vs* O states can be monitored spectroscopically via absorbance. The O state (red) has a characteristic red-shifted absorbance and a narrowing of the two prominent peaks. The O - C absorbance difference spectrum is shown as a gray dashed line. At RT and pH 7.4, the conversion from C to O is nearly complete within 6 hr (3 half-lives). **c**, Thermodynamic schemes including EPIC (top) and human serum albumin (HSA, bottom). EPIC buries the lactone ring of exatecan, protecting it from hydrolysis. On the contrary, binding of exatecan to HSA does not protect it from hydrolysis, presumably because the binding mode exposes the lactone ring, as has been shown for camptothecin-bound structures with HSA. Dominant equilibrium species is shown in bold and enlarged. **d**, Time-resolved absorbance difference spectra of exatecan alone (left) and exatecan with HSA (500 µM), EPIC (20 µM), and EPIC^Q51N/M97L^ (20 µM). The concentration of exatecan was 20 µM for all experiments. Experiments were performed at RT (295 K) in PBS buffer (pH 7.4) and were initiated with 100% ring-closed exatecan from a DMSO stock. Legend denotes time delay. Difference spectra were computed by subtracting the absorbance spectrum at t = 0 hr (t = 1 hr for HSA, due to baseline shifts at early time points) from absorbance spectra recorded at the various time delays. The spectra are labeled by their predominant species at long time delay. Purple and green arrows are positioned at the corresponding wavelengths shown in **e**, pointing in the direction of spectral evolution. **e**, Kinetic traces at 358 nm and 390 nm of the absorbance-difference plots directly above. More data points are shown in the kinetic traces than spectra in **d**, for clarity. No significant spectral evolution was observed for EPIC^Q51N/M97L^ up to 50 hr. Equilibrium yield of O (Y_O_) is computed from a kinetic fit of a two-state mechanism. For HSA, O is considered as O + PO (HSA bound to ring-open exatecan).

## Discussion

The design of small-molecule binding proteins has been a major challenge due to the complexity of simultaneously navigating the spaces of protein structure, protein sequence, and ligand position. Here, we use two reciprocal neural networks in an iterative optimization algorithm, NISE, to efficiently sample these spaces, resulting in the accurate design of high-affinity exatecan-binding proteins entirely from scratch (Figs. 1 and 3). The principle of self-consistency, where a predicted structure closely resembles that intended by the designed sequence, can now be fully unified with small-molecule recognition.

We expect this protocol to be widely adopted for de novo design of small-molecule binding proteins. The described principles are general and are not specific to any particular fold or ligand. The only requirement of NISE is an initial structure of a backbone–ligand pair that can be modeled self-consistently by the two reciprocal neural networks. NISE will then optimize the initial structure within the joint distribution. Here, we used COMBS^6^ to create the initial pair, but other methods could substitute, such as random sampling of ligand positions within a backbone and filtering by structural self-consistency of designed sequences or directly sampling backbones around a target ligand using generative models like RFdiffusion-All Atom^8^ or BoltzDesign1^47^. Indeed, we showed that LASErMPNN and protein–ligand co-structure predictors can be used in combination to statistically enrich a set of backbone–ligand co-structures for those that lead to binders (Fig. E13).

The NISE algorithm is a departure from previous methods^1,2^, which use physics and bioinformatics-based energy functions to optimize the structure, given a sequence designed by a neural network. In these previous schemes, some (and perhaps a major) component of the optimization direction lies orthogonal to the gradient of p(structure, sequence, ligand conformation), which leads to inefficiencies in sampling (see Figs. 1c and E6). This could account for the limited success using similar models like LigandMPNN coupled to Rosetta energy minimization for coordinate ascent^1,2^. NISE bypasses the use of the Rosetta energy function and instead samples directly from the learned distributions of two reciprocal neural networks, iteratively climbing to a higher probability mode in the joint probability manifold, using a computationally derived starting point. Using the same backbone–ligand structure as input, NISE produced designs with nearly 70x higher affinity than those from more traditional methods (7,000x if considering affinity maturation by neural proofreading). NISE achieves this through its ability to overhaul inadequacies in the initial structure, through collective design of each variable: protein shape, ligand conformation, and the entire protein sequence.

LASErMPNN is inherently a probabilistic model, so it was unclear if zero-shot design of small-molecule binders was possible, despite the elevated hit rates from using ProteinMPNN to design protein binders^15–17,19^. We attribute success to the ability to computationally nominate the most designable backbone–ligand structures and then subsequently optimize, filter, and rank LASErMPNN sequences using NISE. It is likely that LigandMPNN could also work as a substitute for LASErMPNN in NISE because both models are trained on the same underlying dataset (the PDB) and show similar performance (sequence recovery) on a test set.

We found that accuracy of LASErMPNN improved through pretraining its ligand encoder on a large corpus of synthetic ligands^36,37^ not seen in the PDB, providing a useful workaround for the lack of chemical diversity in deposited structures. In addition, we showed that LASErMPNN could proofread its own sequences, offering suggestions for single amino-acid mutations that allowed for rapid and dramatic improvement in affinity (Fig. 4). Indeed, the proofread sequence of EPIC was able to protect exatecan from hydrolysis for several days (the longest measured time points) compared to the drug’s typical half-life of a few hours. We anticipate neural proofreading will find wide use in both protein design and natural protein engineering for ligand-aware affinity maturation.

Despite the high success of NISE, a few aspects of the approach could be improved upon in future work. RFAA struggled to model the appropriate bond angles and dihedrals of some ligand functional groups (e.g. the lactone ring), which makes sequence design around these regions challenging (Fig. 5d). Modeling the joint distribution of p(structure, sequence, ligand conformation) will improve as neural networks for co-structure prediction can more accurately capture the underlying distribution of experimentally determined small-molecule conformers. This might be achieved by adding custom constraints or by pretraining on a large dataset of synthetic ligands^36,37^ with coordinates optimized through quantum chemistry, as we showed for LASErMPNN. Additionally, RFAA struggled to predict the correct orientation of exatecan for many designs (Fig. E5), although the success rate iteratively improved during NISE. Failure to predict the correct orientation may be a shortcoming of the structure predictor, of the sequence design model, or both. Using new structure predictors such as Boltz-1 could help retain more designs, making NISE more efficient. Indeed, replacing RFAA with Boltz-1 in an identically performed NISE trajectory for exatecan-binder design showed improved prediction of ligand orientation, faster convergence to high-confidence ligand poses, and sequences with lower NLL from LASErMPNN (Figs. E24 and E25).

The design of small-molecule proteins is now approaching the hit rates of the design of PCR primers, yet the rules of protein–ligand molecular recognition are vastly more complex than Watson–Crick base pairing of nucleic acids. We have shown that modern deep neural networks, coupled with appropriate training data, can capture and distill this complexity, which can ultimately be extracted using design algorithms like NISE. We anticipate these capabilities will catalyze the use of bespoke proteins as ready-made reagents to manipulate biology at the single-molecule level.

## Methods

### LASErMPNN Model Details

#### LASErMPNN Overview

Ligand Aware Sequence Engineering Message Passing Neural Network (LASErMPNN) is an autoregressive SE(3)-equivariant graph neural network that takes as input a point cloud containing 3D coordinates encoding a protein backbone and any additional heteroatom coordinates, including metal, nucleic acid, or small-molecule ligand coordinates. The model outputs a choice of amino-acid sequence and dihedral angles encoding sidechain orientations, which are used to build all-atom protein structures. The input point cloud is parameterized as a heterograph consisting of two distinct node types corresponding to the protein backbone and non-protein context atoms respectively. LASErMPNN consists of three core modules: a pre-trainable ligand encoder, a heterograph encoder, and a heterograph decoder (Fig. S1).

#### Heterograph Construction

Protein backbone nodes are created for all backbone “frames” consisting of the N, C⍺, C, O, and Cβ atom coordinates for a single residue (Cβ atom coordinates are created from idealized bond lengths and angles if absent). Ligand nodes are constructed for all atoms in the ligand including hydrogens. A heterograph representation of the input structure with distinct edge types for each node pairing (protein-protein, protein-ligand, and ligand-ligand) is constructed by combining the protein-protein and protein-ligand subgraphs. During model training and inference, all protein backbone frames in the input structure are idealized through the alignment of an idealized alanine backbone frame before graph construction, which removes crystallographic artifacts in the backbone that could bias training while also preserving sidechain dihedral angles for the prediction of sidechain conformation.

Protein node representations are initialized to a vector of zeros, and ligand nodes are initialized to concatenated one-hot encodings of the group and period in the periodic table for each atom’s atomic number. The 5 nearest ligand atom neighbors of each ligand atom are computed (including self-edges) for the ligand-ligand graph and distances between them are computed and embedded with Radial Basis Functions (RBFs) as described in ProteinMPNN with 75 bins encoding distances between 0.0 Å and 15 Å^30^. Protein-ligand edges are constructed only for protein nodes with C⍺ carbons within 20 Å of a ligand atom. The 48 nearest ligand heavy atoms or polar hydrogens are connected to these protein nodes and the distances between all backbone frame atoms and the ligand atom are computed and embedded in RBFs with 75 bins encoding distances between 0.0 Å and 15 Å. Protein-protein edges are constructed similarly with all 25 pairwise distances between backbone frame atoms computed to the nearest 48 protein nodes (including self-edges) and embedded in RBFs with 22 bins encoding distances between 2.0 Å and 22 Å as described in ProteinMPNN. These 3 subgraphs are concatenated by taking the union of all node and edge sets forming the final heterograph representation of a protein-ligand complex.

#### Ligand Encoder Pretraining

Due to potential limitations in the diversity of small molecules in the PDB, we hypothesized that we could improve the model by pre-training the ligand encoder module that constructs ligand-atom embeddings on a larger, more diverse dataset of 3D ligand conformers. To do this, we use the SPICE-2 dataset which contains over 100,000 protonated systems of small molecules and ions ranging from 2 to 110 atoms^37^. Each training example has associated atomic partial charges and dipole vectors computed by density functional theory (DFT) calculations, which we use to train a neural network that predicts these features.

The inputs to the ligand encoder module are a nearest-neighbor graph constructed from the 3D ligand coordinates and the element of each atom as described above. We train a graph neural network to predict the partial charges (mbis_charges) and dipole vectors (mbis_dipoles) from the SPICE-2 dataset as well as molecular-connectivity derived features such as atomic hybridization, aromaticity labels, atomic formal charges, and bond orders computed using RDkit^48^. We freeze the weights of this module when training the full LASErMPNN model and use it to generate ligand atom embeddings, which are used downstream when performing ligand-to-protein message passing operations. Freezing these weights during training allows us to check the ligand encoder’s understanding of novel ligands by predicting partial and formal charges, hybridizations, etc, and evaluating these predictions against the ground truth.

The ligand encoder graph neural network employs three rounds of GATv2 message passing to build up 256-dimensional node embeddings for each ligand atom^49,50^. From these embeddings, separate 3-layer multilayer perceptron (MLP) adapter layers are used to predict the partial charge, atomic hybridization, formal charge, the number of covalently attached hydrogens, the number of covalently bonded neighbors, and whether the atom is aromatic^49^. These tasks are treated as a classification over discrete bins, and cross-entropy loss is applied and summed across all classification tasks. Dipole vector regression is performed with a 3-layer Geometric Vector Perceptron (GVP) adapter layer^51^. Initial vector representations are computed from a weighted average of neighbor atom displacement vectors and are updated after each round of GATv2 message passing using GVP layers prior to the final dipole regression layer. Only scalar embeddings from the ligand encoder are used for protein-ligand message passing.

#### LASErMPNN Inputs and Outputs

LASErMPNN discards any pre-existing protein sequence information contained within the input coordinates (unless instructed otherwise) and applies iterative rounds of message passing with its encoder module to build up representations for each protein node. The decoder module takes as input a randomized decoding order which is ultimately used to select a choice of amino acid (aa) and sidechain dihedral angles for each protein node and conditionally updates embeddings from the encoder module with information (predicted aa identities and dihedral angles) about previously decoded nodes.

The output of LASErMPNN is a protein sequence and associated dihedral angles that are used to create a 3D model of a protein structure constructed from the idealized input backbone and any available context atoms, including ligand heteroatoms. The model produces predicted orientations for all sidechain atoms including rotatable and non-rotatable hydrogens placements enabling the evaluation of designed hydrogen bonding interactions directly from LASErMPNN outputs.

#### LASErMPNN HeteroGATv2 Message Passing

LASErMPNN extends the ProteinMPNN architecture with SE(3)-equivariant vector representations using Geometric Vector Perceptron (GVP) layers and a Graph Attention Network inspired heterograph message passing formulation. The GVP layers are used to place equivariant vector representations on all protein nodes, which are used to encode residue orientation (Figure S9). We hypothesized that these vector representations would be useful for rotamer prediction as this task inherently depends on a robust encoding of frame orientation. To extend the

ProteinMPNN architecture to handle protein-ligand heterographs, we combine ProteinMPNN message passing layers with a heterograph generalization of the Graph Attention Network (GAT) architecture^49,50^. We apply this “Hetero-GATv2” architecture to jointly handle ligand-to-protein messages and protein-to-protein messages in both the encoder and decoder modules of LASErMPNN with message passing operations defined as follows:

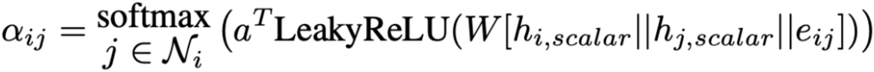

with separate learnable 𝑎 and 𝑊 matrices for protein-protein and protein-ligand node pairs. ℎ*_i,scalar_*_’_ and ℎ*_j,scalar_* refer to the scalar representations stored in node 𝑖 and node 𝑗 respectively while 𝑒_!(_ refers to the edge information stored for the edge connecting nodes 𝑖 and 𝑗. “||” denotes concatenation. A single softmax operation is applied over all pre-softmax attention coefficients so attention can be selectively paid to nodes of potentially different types in the neighborhood of a given node 𝑖. Node updates are computed as follows with 𝑊’ which is a separate set of learnable parameters from 𝑊 for a given node pair with 𝑙 denoting the index of the current message passing layer:

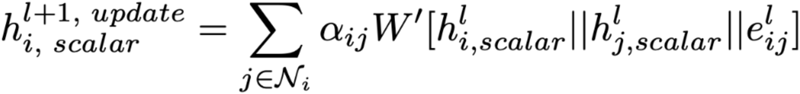

The final scalar representation update is then computed with residual connections as in the ProteinMPNN architecture:

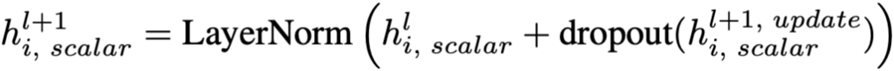

Vector representation updates are computed separately from the scalar updates using a GVP layer:

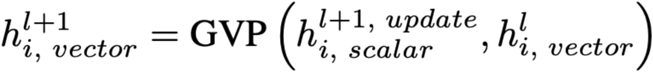

#### LASErMPNN Loss Terms

We hypothesized that we could improve the quality of incorrect predictions made by the model by introducing a loss based on statistics from multiple sequence alignments (MSAs), which we accessed from OpenProteinSet^52^. To do this, we add an additional cross-entropy term to the loss for residues whose highest probability is not the ground-truth. We compute a target distribution for these incorrectly predicted residues by computing per-column aa frequencies from the MSA and then compute the cross-entropy of the sequence logits produced by the model vs these frequencies. We dampen this additional loss term as a function of the depth of the MSA, linearly interpolating the weight between 0 for MSAs of depth 0 and 1 for MSAs of at least twice the median depth of 96 sequences. To prioritize sequence recovery, this loss term is further dampened by a factor of 0.1 relative to the main sequence classification loss.

To predict the dihedral angles of rotatable sidechains, we discretize the angles into 5-degree bins and encode these into a 72-bin circularly symmetric version of the RBF encoding used in the protein-protein edges to encode distance information (as described in ProteinMPNN). An offset between +/- 2.5 degrees within each bin is regressed with a mean squared error loss term and is summed with the angle represented by the predicted bin for a final dihedral angle prediction. The choice of dihedral angles is conditioned on the embeddings used to produce the choice of aa and the identity of the selected aa. Zero to four dihedral angles are decoded for each residue, and previously decoded dihedral angles are used to compute the next angle until the rotamer is fully decoded. Angles which do not exist for a given amino acid are predicted but masked out from the loss during training. Overall, the loss used in training the LASErMPNN model is:

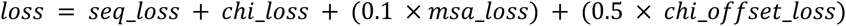

where seq_loss refers to the cross-entropy term over the protein sequence, chi_loss refers to a cross entropy loss over the predicted dihedral angle bins, msa_loss refers to a cross entropy computed from the sequence logits to the frequencies computed from MSA frequency statistics, and chi_offset_loss refers to a mean squared error loss term applied to the offset between +/- 2.5 degrees of the predicted chi angle degrees within each 5 degree dihedral angle bin.

#### Training & Inference Differences

During training, the sequence and rotamers are teacher-forced to condition on the ground truth for any previously decoded states. This means that when predicting dihedral angles, regardless of what the highest predicted aa probability would be, the model is told what the ground truth aa is before the first dihedral angle is predicted ensuring ground truth values are available for the loss computation. We use causal masks to selectively provide information as it would be available during autoregressive decoding while allowing fast computations of logits for all residues in parallel during training. During inference, the sequence and rotamers are sampled in a fully autoregressive manner following the input decoding order. Starting at the first residue in the decoding order, a choice of aa is predicted, then its first sidechain dihedral angle is predicted conditioned on the sequence choice and decoder embeddings. The process repeats, conditioning on previous samples until each of the sidechain dihedral angles (up to four) are predicted. The model then moves to the next protein node in the decoding order and repeats the sampling until all residues are designed.

#### Protein Residue Frame Noising

Small changes in the relative positioning of the backbone coordinates can result in high RMSD (> 1.0 Å) when rebuilding large rotamers from sidechain dihedral angles due to the lever-arm effect. To noise the protein complex to increase model robustness while also ensuring the model may still learn to predict sensible dihedral angles, we apply noise at the level of the backbone frame rather than on each backbone atom independently. This keeps the relative positioning of the N, C⍺, C, O, and Cβ coordinates constant and minimizes the lever-arm effect ensuring predicted dihedral angles remain close to the ground-truth before noising. We find that this strategy combined with backbone frame idealization removes any subtle information leakage from crystallographic artifacts contained in the backbone frame coordinates while improving the quality of the model’s predicted rotamers over noising each atom independently. We train the LASErMPNN model with 0.1 Å Gaussian noise applied to the protein backbone frames and 0.05 Å Gaussian noise applied to each ligand atom independently. The protein backbone noise level of 0.1 Å was determined by sweeping sequence foldability as a function of noise for a monomeric test set (see Training noise levels and foldability optimization).

#### Training LASErMPNN

To train the LASErMPNN model, we treat the task of sequence prediction as a classification over the 20 natural amino acids and therefore penalize incorrect sequence predictions with a sum of cross-entropy loss normalized by the number of “valid” residues. To define a valid residue, we select only canonical residues from the chains that were sampled from the dataset to construct the current batch. We mask out from the loss any residues in contact with a non-canonical amino acid as well as those which were loaded as part of the same biological assembly as the sampled chain.

To define an epoch of training, we sample once from each 30% sequence identity cluster (see Dataset Curation and Splitting), selecting a subcluster at random (70% sequence identity), then choosing one sequence from within each subcluster. If the sampled sequence exists in multiple biological assemblies, we sample one at random. We use a similar chain masking strategy to that used to train ProteinMPNN and selectively mask information about similar chains based on structural homology thresholds. Specifically, any chain loaded as part of the same biological assembly as the sampled chain with a TM-score less than 0.7 is provided as sequence and structure context for conditional generation, otherwise the sequence and rotamer identities are masked and decoded with the sampled chain with cross-entropy loss only applied to the sampled chain.

Deep learning experiments were performed using PyTorch and PyTorch Geometric primarily on 4x NVIDIA A6000 GPUs. Models were trained until train and test sequence accuracy curves intersected (typically around 60,000 optimizer steps) using the NoamOpt optimizer with learning rate annealing as was described in ProteinMPNN. Models were trained with a node dimensionality of 256 and an edge dimensionality of 128 on complexes of less than 2,500 residues or ligand atoms.

#### van Der Mer (vdM)-Inspired Pseudoligand Sampling

To increase the diversity of binding pockets the model is trained on, we use a residue atomization strategy similar to that reported in recent work^25,53^. Briefly, we use individual amino acids that are part of the polypeptide chain as “pseudoligands” (Fig. S1). We choose pseudoligands with at least 3 contacting residues (heavy atoms within 5 Å) that are distant in the linear polypeptide sequence (greater than 6 residues in primary sequence). We drop the sequence-local residues from the protein graph to form synthetic binding sites. We compute phi and psi backbone dihedral angles for each residue and use these to align methyl caps onto the sampled residue backbone frame, ensuring the residue is built with explicit hydrogens and is thus in-distribution with the training data of the ligand encoder.

### Dataset Curation and Splitting

#### Protein Data Bank Dataset Curation

To construct the LASErMPNN training dataset, we began with all experimentally determined structures in the RCSB PDB deposited before 06/01/2022. We downloaded mmCIF formatted data for all biological assemblies. We filtered out structures with over 5,000 amino acids or ligand residues and selected structures generated by X-ray crystallography or cryo-electron microscopy with at least 3.5 Å resolution. We then ran Reduce on the dataset to predict explicit hydrogen locations, adjust protonation states, and to perform N/Q/H residue rotamer flips^54^.

Covalently modified and non-canonical amino acids were recorded as “X” residues with masks stored which drop these and any contacting residues (heavy atoms within 5.0 Å) from the loss during training as “X” residues cannot be used for sequence design. We then idealize the backbone frames of all residues by aligning an ideal alanine onto each residue’s crystallographic N, C⍺, and C coordinates and rotate the idealized O atoms using the psi angle. Sidechain dihedral angles are then computed relative to these idealized frames, which are used as the ground-truth for rotamer decoding.

We tracked the full experimentally characterized sequences for each protein monomer within each biological assembly using information contained in the header of the mmCIF files and mapped these sequences to the OpenProteinSet MSA database which were used to compute an MSA-based loss term during training^52^. The dataset was then split into clusters of 30% sequence identity using MMSEQs easy-cluster and then sub-clustered further at 70% sequence identity within each cluster (see MMseqs2 sequence clustering code)^55^. The full processed dataset can be found on Zenodo^56^.

#### LigandMPNN Dataset Split Reconstruction

The LigandMPNN training code and dataset are not currently publicly available. An exact reconstruction of the model is therefore difficult as the sequence clusters used in the LigandMPNN dataset were not reported, and the test/validation splits are reported at the level of PDB codes rather than at the level of protein chains. Since biological assemblies of a given PDB code may contain multiple protein chains belonging to different sequence clusters, there may be train set chains in the test set biological assemblies making direct comparison between architectures difficult. Subtle differences in training cutoffs and resolution thresholds may also further complicate these comparisons. For the fairest comparisons possible, we reconstruct the LigandMPNN training dataset starting from our curated PDB data.

To do this, we dropped any 30% sequence identity cluster from our dataset containing any chain with a PDB code in the reported LigandMPNN validation or test sets. We then ran MMseqs easy-search to check the remaining data available for training for any sequences over 30% similarity to those in the validation or test sets ensuring there was no contamination. We then reimplement the LigandMPNN architecture and retrain it on our reconstructed LigandMPNN training dataset and evaluate the performance of these models on the “small-molecule” test set PDB codes (as opposed to the nucleotide/metal test set PDB codes which, while still held-out from training, we did not seek to optimize performance for).

#### Streptavidin Held-out Dataset Split Construction

We also sought to create a dataset split which rigorously holds out the streptavidin fold to evaluate model generalization to novel backbones. To do this, we hold out the 30% sequence identity cluster belonging to the streptavidin protein family and then further remove any sequences over 30% identity over at least 25 aligned residues to any sequence in the dropped streptavidin cluster using MMSEQS easy-search. To check for structural contamination and streptavidin fusion proteins, we TM-aligned all members of the streptavidin sequence cluster against all remaining data available for training. To do this we took the first chain of all streptavidin cluster biological assemblies to check for structural homology to the monomer as well as the full biological assembly. We then dropped any structures from the training dataset with a monomer or biological assembly TM score above 0.5 with greater than 25 aligned residues. We searched the PDB for all complexes containing biotin (residue code BTN) and visually inspected the set of structures containing BTN which were not yet dropped from the training dataset for the streptavidin fold, confirming the absence of any streptavidin-like folds bound to biotin. Finally, we checked the remaining dataset for any chains with CATH assignments of “Avidin-Like” as well as for circular permutants of the streptavidin sequence by permuting the PDB Code 4jnj sequence in 10 residue increments and checking the remaining dataset with MMSEQS easy-search. These analyses confirmed that all streptavidin-like sequences and structures were dropped from the training dataset for the streptavidin held-out split.

### Design of Exatecan-Binding Proteins

#### NISE Protocol Implementation

We devised an iterative “flexible-backbone” design loop (NISE) and applied it to an initial protein-exatecan binding pose generated by COMBS^6^. The NISE protocol uses LASErMPNN to design many proteins for the input pose (expansion) and uses protein-ligand co-structure prediction to generate ligand-bound structures for each sequence. Throughout the design loop, we keep a running list of up to 3 backbones sorted by highest backbone priority observed over all iterations of the design loop. In this work, we determined backbone priority as ligand pLDDT predicted by RoseTTAFold All-Atom (RFAA)^8^. By maximizing ligand confidence through ligand pLDDT, protein confidence (which can be tracked through protein C⍺ pLDDT) is implicitly also maximized as the two values are highly correlated (Fig. S2). We assigned the input backbone a priority of 1, ensuring it was used for sequence generation throughout the loop. As a consequence, two additional protein-ligand poses were considered for input to LASErMPNN for sequence generation at each iteration. (Using the input backbone within each sampling loop is likely unnecessary for NISE optimization and could feasibly be omitted in future implementations.) Each protein-ligand pose was used to generate 1000 sequences that were then predicted by RFAA using 15 recycles. Sequences with high protein C⍺ pLDDT (> 0.8) and low backbone C⍺ and ligand heavy atom RMSDs (< 1.5 Å) to the previous round’s input to LASErMPNN were ranked by ligand pLDDT. All ligand RMSDs were computed only over the heavy atoms in the ring system, excluding the rotatable carbon chain on the lactone ring of exatecan. New backbones (up to two) were chosen each round if ligand pLDDT surpassed that of a previously observed design. An early version of the LASErMPNN weights (prior to hyperparameter tuning) was used for sequence generation, the weights and inference code for which are available on GitHub under the name *epic_weights.pt*.

#### Design Filtering and Ranking

The NISE protocol was run for 35 iterations, at which point the ligand pLDDT had converged. We pooled the sequences across each iteration (100,000 sequences total) and filtered by self-consistency metrics (Fig. E7). Namely, we filtered out designs that did not satisfy any of the following criteria: i) protein C⍺ RMSD < 1.5 Å, ii) ligand heavy atom RMSD < 1.0 Å, iii) ligand pLDDT > 0.8, iv) protein C⍺ pLDDT > 0.8, and v) a core packing metric (packstat) computed with the program Rosetta > 0.55 after sidechain and backbone minimization using the Rosetta energy function. Rosetta energy minimization of the protein-ligand complex (backbone, sidechain, and ligand) was applied using the TaskAwareMinMover and Ref15 score function to the RFAA output structures to produce protonated structures which could be evaluated for designed hydrogen bonding interactions. Designs with protein and ligand RMSDs between the RFAA output and the energy minimized structure greater than 1.5 Å were also removed. To check for binding site preorganization, we also applied Rosetta energy minimization in the absence of the ligand to allow relaxation of binding-site residues. We reintroduced the ligand into the unbound structure by superimposition of the bound structure onto the unbound structure via binding-site residues. We then counted heavy atom clashes with the ligand (sidechain atoms within 2 Å of any ligand heavy atom). We removed designs for which the minimized unbound protein had any clashes with the ligand. Designs containing cysteine residues or buried charged residues were also removed. We then ranked designs for experimental characterization based on the number of buried, non-H-bonded, polar atoms of the ligand using the Rosetta energy-minimized structures. Two copies of each designed protein sequence were predicted using AlphaFold and were visually inspected for domain swaps or low interface pAE (pAE < 14; residues within 5.0 Å in predicted multimer), which we used as a proxy for aggregation potential. Four of the top ten exatecan designs remained after this process and were chosen for experimental characterization. Reranking these designs using a linear combination of the number of buried, non-H-bonded (“unsatisfied”), polar atoms (BUNs) of the protein and ligand (rather than only using the contribution from the ligand) ranks the EPIC sequence 1st out of the remaining top 4 designs (Table S1). We used BUNs Score = 2x ligand BUNs + 1x protein BUNs.

#### Model Benchmarking

Benchmarking LASErMPNN with Streptavidin–Biotin complex We used LASErMPNN to design sequences for monomeric streptavidin bound to biotin. We used each chain in the asymmetric unit of PDB accession 4jnj to design 10,000 sequences (2,500 per chain). For this task, we used the LASErMPNN model trained on 0.1 Å noise with the strict streptavidin held-out dataset. We designed sequences using sampling temperature of 1.0 for the binding-site residues and a temperature of 0.1 for all other residues. We sampled sidechain dihedral angles using argmax sampling (1e-6 chi angle sampling temperature). The designed sequences had self-consistent backbones when folded in single-sequence mode by RFAA, but RFAA struggled to predict the correct binding mode of biotin in most cases. When the same sequences were folded using the Boltz-1 model^33^ in single-sequence mode, the correct binding mode was observed in most designs, suggesting that alternative co-structure predictors may be more robust than RFAA (Fig. E2).

We filtered the Boltz-1 predicted structures for self-consistency (protein C⍺ RMSD < 2.0 Å and ligand heavy-atom RMSD < 2.0 Å). After ranking the remaining ∼1700 designs by BUNs Score, we found the design with the highest binding-site sequence recovery in the top 25 designs (binding-site sequence defined as residues that are within 5 Å heavy-atom distance of a biotin heavy atom). We also saw a general correlation between ranking by BUNs and binding-site sequence recovery (Fig. E3). On average, designs with lower BUNs Score had a higher binding-site sequence recovery.

#### Testing ranking protocol of LASErMPNN designs using rucaparib-PiB complex

We used a previously designed four-helix bundle, PiB (PARP-inhibitor binder), that tightly binds the drug rucaparib, as a relevant test case for ranking LASErMPNN designs (Fig. E4). We used chain A from the rucaparib-PiB crystal structure (PDB accession 8tn6) and the baseline LASErMPNN model (laser_weights_0p1A_noise_ligandmpnn_split.pt) trained with 0.1 Å frame noise. We generated 1000 sequences using a sequence sampling temperature of 0.2 with sidechain dihedral angles sampled at argmax temperature (1e-6) with Cys residues disabled. To filter the designed sequences, we applied self-consistency filters of 1.5 Å for both protein C⍺ RMSD and ligand heavy-atom RMSD, as well as a ligand pLDDT filter of > 0.8 and a protein C⍺ pLDDT filter of > 0.85. A final preorganization filter was applied by removing the ligand and running fixed-backbone Rosetta sidechain energy minimization in its absence, then subsequently realigning the ligand into the repacked binding site. Clashes were defined as sidechain heavy atoms within 2.0 Å of the ligand heavy atoms and only the resulting 211 designs with 0 clashes were kept for the final ranking.

The BUNs Score described above for streptavidin-biotin was used to rank the remaining 211 designed sequences. PiB was ranked as the 4th best sequence (using ligand pLDDT to break ties between designs with the same BUNs Score). 6 designs generated by LASErMPNN scored the same as the PiB sequence by BUNs Score. The highest binding site recovery design was 78.9% and ranked 32nd by BUNs Score, using pLDDT for tie breaking.

#### Details of ablation studies

We ablated several model features to quantify their contributions to the computational performance of LASErMPNN (Table E1). Ablations were performed on a baseline model trained with node- and edge-embedding dimensions of 256, an optimizer warmup rate of 5,000 steps, backbone frame noise of 0.1 Å, and ligand noise of 0.05 Å. Models which use a pretrained ligand encoder (SPICE2 training dataset) use the same model with node dimension of 256 and edge embedding dimension of 128. Performance was analyzed over the “small-molecule” subset of our reconstructed LigandMPNN test set. All LASErMPNN models were trained to convergence using idealized backbone frames and frame-noising (train and validation sequence recovery curves intersecting, typically around 60,000 optimizer steps). We retrained a LigandMPNN model, which we trained using per-atom noising with idealized backbone frames, 2% residue atomization (rather than our pseudoligand sampling protocol), node/edge dimensionalities of 128, and 300,000 optimizer steps (as reported by its authors^25^). All evaluations were performed on the same test set of 317 small-molecule binding proteins from the LigandMPNN test set with idealized backbone frames to avoid information leakage from artifacts in the crystallographic backbone coordinates emulating the de novo design task.

For all sequence recovery and BLOSUM62 substitution score calculations, 5 sequences were generated by each model using argmax sampling (1e-6 temperature) for both sequence and sidechain dihedral angles (where applicable). Results were averaged over the selected residues. The LigandMPNN (retrained) model refers to a reimplementation of the LigandMPNN architecture. While the LigandMPNN (author weights) model refers to the weights trained by the LigandMPNN authors (ligandmpnn_v_32_010_25.pt) on their unreleased processed dataset and clustering of PDB chains. Differences in the clustering of the PDB chains used during training could be responsible for the observed performance difference between the LigandMPNN public weights and our reimplementation. Our reimplementation allows for a direct architecture comparison as it was trained on the exact same dataset.

We computed BLOSUM62 substitution scores (A higher score is a better substitution.) to quantify the quality of the incorrect sequence predictions made by the models (Table E1). After computing sequence recovery, any incorrectly predicted residues were scored using the BLOSUM62 matrix as a proxy for physiochemical similarity. The final BLOSUM62 score averages the substitution scores of incorrectly predicted residues over binding sites (residues with heavy atoms within 5.0 Å of a ligand heavy atom) or the full sequence.

The “LASErMPNN (add edge angle features) model” performs the best by sequence recovery metrics and BLOSUM62 substitution scores. This model extends the baseline LASErMPNN model by adding angular information about ligand atom placement relative to the backbone, inspired by the “_make_angle_features” function used in LigandMPNN, but implements the encoding of angular information using dot-products of the learned equivariant protein node vector representations with normalized ligand atom displacement vectors rather than using Gram-Schmidt decomposition to build local frames. The performance reported for all models is relative to the baseline LASErMPNN model which does not incorporate these ligand displacement angular features.

Ablating either the MSA loss or sidechain-dihedral-angle loss results in model performance similar to a reimplementation of the LigandMPNN architecture, which is expected as these models were trained with the same objective functions. Ablating the pretrained ligand encoder results in worse performance compared to the LigandMPNN architecture, indicating that the ligand encoder of LASErMPNN may be more difficult to optimize than the ligand encoder of LigandMPNN using the PDB alone. This result may be rationalized by the ligand encoder of LigandMPNN containing more optimizable parameters (∼3,000,000 vs ∼400,000 in LASErMPNN). Ablating the equivariant protein node vector representations, by initializing them with zeros, drops performance slightly, suggesting these have a small contribution to overall performance.

#### Neural proofreading protocol

We sought to identify single amino-acid mutations that could improve binding affinity of EPIC for exatecan. We used LASErMPNN in a process we call neural proofreading to determine if any residues near the ligand could be mutated to a higher probability residue, given the identities of the remainder of the EPIC sequence. An energy-minimized RoseTTAFold-All Atom prediction of the EPIC sequence (and exatecan ligand) was used as input to the proofreading protocol. Energy minimization was performed using OpenMM^46^ and the Amber-14 forcefield with the SMIRNOFFTemplateGenerator to parameterize the ligand forcefield. The energy-minimized structure of exatecan-bound EPIC was fed into LASErMPNN, and the sequence identities of all residues were fixed.

The binding site residues were identified as those with heavy atoms within 5 Å of any ligand heavy atom. Each binding-site residue was independently unfrozen and sequence probabilities were computed conditioned on the identities of the remaining 147 residues of the EPIC sequence. LASErMPNN probabilities show a slight dependence on the decoding order (the order in which the sequence and dihedral angles are chosen), despite being trained using many random decoding orders. We therefore used multiple decoding orders to provide additional robustness to the computed leave-one-out probabilities. We also turned on dropout for additional diversity in the sampled probability distributions. During proofreading, the model is additionally tasked with repacking the sidechains of all residues, since changes in neighboring rotamers may also influence proofreading probabilities. Proofreading probabilities were created by averaging over 10 decoding orders and 10 model dropouts then renormalizing the resulting distribution. We selected mutations in two buried positions that showed higher probabilities than the amino acid in the EPIC sequence, precluding any mutations that would introduce buried unsatisfied polar atoms. We chose a Gln to Asn mutation at position 51 (EPIC^Q51N^) and a Met to Leu mutation at position 97 (EPIC^M97L^), both of which showed higher probability through proofreading (Figs. 4b and Fig E15). An early version of the LASErMPNN weights (prior to hyperparameter tuning) was used for sequence generation, the weights and inference code for which are available on GitHub under the name *epic_proofreading_weights.pt*.

#### Training noise levels and foldability optimization

To tune the hyperparameter that governs the amount of gaussian noise applied to the per-residue frames of the protein, we subsampled the LigandMPNN split validation set for 882 monomeric protein backbones with less than 300 amino acids. We swept the noise level applied to backbone frames and computed the C⍺-LDDT between the AlphaFold2-predicted structures of designed sequences and the input crystal structures to LASErMPNN (Fig. E1). We identified 0.1 Å frame noise to maximize sequence recovery and AlphaFold2 success rates.

#### AlphaFold3 Self-Consistency Test Set Evaluation

We wished to uncover which version of LASErMPNN or LigandMPNN could best perform for the small-molecule-binding protein design task. For this, we used a proxy of achieving self-consistent protein-ligand co-structures with AlphaFold3 (AF3). To test this, we compare the LASErMPNN model to our retrained LigandMPNN model on i) a small set of monomeric proteins from the LigandMPNN test set, ii) the initial helical bundle backbone used as input to the NISE optimization loop and iii) a computationally generated beta barrel structure (b11_L3B_11c.pdb)^4^. Using each network, we generated 250 sequences for each input protein-ligand complex, and predicted protein-ligand co-structures with AF3 using 10 recycles and 5 diffusion samples.

Overall, the LASErMPNN architecture and LigandMPNN architectures perform similarly well for these tasks (Figs. S3-S5). We saw a slightly lower binding-site and ligand RMSD from sequences generated by LASErMPNN for the de novo backbones (Figs. S3-S5). Where performance between the models differed, we hypothesized that some of this difference may come from the backbone-frame noising strategy used to train the baseline LASErMPNN model (LASErMPNN Frame Noise). In this noising strategy we only idealize the frames and add translational noise (no rotational noise) to preserve the dihedral angle predictions. We therefore trained and tested a different version of LASErMPNN with a per-atom noising strategy on top of the idealized frames (LASErMPNN Per-Atom Noise). Indeed, in a structure where the baseline LASErMPNN model underperforms relative to the LigandMPNN architecture with respect to self-consistency RMSDs (PDB accession 5eng), the LASErMPNN Per-Atom noise model appears to bridge the performance gap (Fig. S3).

The models perform similarly when comparing the confidence of predicted structures and native sequence recovery (Figs. S4 and S5). The baseline LASErMPNN model generates slightly higher ligand pLDDT (AF3) sequences than the other models on the de novo backbones when considering only self-consistent sequences.

#### Computation of buried, non-H-bonded atoms

To compute the number of buried, non-H-bonded, polar atoms in a design, we energy-minimized (sidechain and backbone) the predicted protein-ligand co-structures from RoseTTAFold-All Atom using RosettaScripts and the ‘-no_optH False’ command line flag (see Rosetta code for fixed backbone sidechain minimization)^57^. The resulting protonated and energy-minimized structures were analyzed for buried, non-H-bonded, polar atoms. To determine atom burial, we compute a protein surface from a convex hull of C⍺ and Cβ coordinates using an alpha-sphere size of 9 Å to define the interior and exterior of the protein^6^. We calculate the distance of polar atoms to the convex hull to determine burial and calculate H-bonding partners by interatomic distance (2.5 – 3.5 Å heavy-atom distance).

#### Rotamer analysis

We evaluated the sidechain packing performance of the baseline LASErMPNN model using the LigandMPNN test set (Fig. S6). We fixed each native protein sequence and generated 5 packing solutions with different decoding orders at argmax chi-angle sampling temperature for each protein complex. We group each residue into 3 burial categories based on the number of Cβ atoms within 10 Å of its Cβ atom. Sidechain heavy-atom RMSD was computed accounting for residue sidechain atom symmetries for ARG, ASP, GLU, PHE, and TYR.

#### Analysis of NISE design trajectory

Throughout the iterative selection and expansion algorithm, the confidence of the sequences generated at each iteration by the LASErMPNN model, P(sequence | structure, ligand), can be tracked by monitoring the negative log-likelihood (NLL). As an amino acid sequence is generated, the probability of each residue being the amino acid that was sampled during the decoding process can be tracked. This is computed as NLL = <-log (pi)>, where pi is the probability generated by LASErMPNN of residue i being the sampled amino acid with the mean taken over all residues in the protein. Lower values of the NLL correspond to more confident sequences generated by the model. The confidence of the ligand placement generated by structure-prediction methods like RFAA, which we loosely approximate as P(structure | sequence, ligand), can be approximated with the ligand pLDDT (higher values more confident). We track the optimization of pLDDT and sequence NLL throughout the optimization cycle initialized with the COMBS generated exatecan binder starting pose (Figs. 1c and E24). Plotting the mean pLDDT and NLL per iteration shows that the loop rapidly increases in both sequence and structure confidence for the generated datapoints and then plateaus. The optimization using RFAA appears noisier than an identical optimization using Boltz-1 as the structure-prediction network. The exact values to which the optimization converges are a function of the noisy optimization process and the starting pose as constraints on self-consistency restrain the space of structure that NISE can explore to structures that are similar to the input pose.

#### Rosetta iterative selection–expansion trajectory

An iterative selection–expansion trajectory was constructed which mimics that which generated EPIC while replacing the structure prediction network (RFAA) with Rosetta for backbone relaxation (Fig. E6). In place of pLDDT from RFAA, we used Rosetta ligand energy as a readout of pose quality. To do this, the LASErMPNN model was used to generate sequences and initial rotamer predictions which were then energy minimized in Rosetta. Energy minimization over the backbone and sidechains was performed using RosettaScripts with a fixed ligand conformer. The total ligand energy was extracted from the Rosetta output pdb file. For the next round of design, we selected the top-3 structures with lowest ligand energy, protein C⍺ RMSD < 1.5 Å, and ligand heavy atom RMSD < 1.5 Å, additionally requiring the ligand to remain buried in the convex hull of the protein (defined by the protein C⍺ coordinates). We repeated this iterative selection-expansion process for 70 iterations. Repeating this analysis with a flexible ligand conformer during energy minimization produces very similar results (Figure S10). Using Rosetta ligand energy to guide an iterative selection–expansion trajectory is a less efficient optimizer of LASErMPNN sequence likelihood compared to using ligand pLDDT from the RFAA or Boltz-1 structure-prediction networks (Fig. S11).

#### Correlation of ligand pLDDT and binding affinity using AF3, Boltz-1, and RFAA

We wondered if ligand confidence of predicted structures would correlate with binding affinity between EPIC, its mutants, and various ligands (Fig. E20). Protein-ligand co-structure prediction was performed using AF3 and Boltz-1 with default inference parameters (10 recycles, 5 diffusion samples vs 3 recycles, 1 diffusion sample). RFAA sampling was modified to use 100 recycles. For certain ligands and for EPIC^Q51N^ (red circles, Fig. E20), RFAA predicts a flipped orientation despite the large number of recycles.

#### Creation of exatecan conformers

We created two exatecan conformers starting from the coordinates of small-molecule crystal structure of a camptothecin derivative deposited in the Cambridge Structural Database (CSD, deposition number 1504071, 7-ethyl-10-hydroxycamptothecin monohydrate). We modified the structure using Avogadro software and minimized the newly built coordinates using the UFF forcefield in Avogadro, including all hydrogens. The two conformers differed by a 220-degree rotation about the torsional angle of the rotatable ethyl substituent.

#### Small molecules

Small molecules were purchased from commercial providers. Belotecan hydrochloride (Selleck Chemical, S6631) and irinotecan (SC-269253, Santa Cruz Technology) were dissolved at a concentration of 10 mM in DMSO. Exatecan mesylate (HY-13631A, MedChemExpress) was dissolved at a concentration of 3.76 mM in DMSO. Camptothecin (13637S, Cell Signaling Technology) was dissolved at a concentration of 5 mM in DMSO. FL118 (HY-12486, MedChemExpress) was dissolved at a concentration of 2.55 mM in DMSO. All small-molecule stock solutions were aliquoted and stored at -20 °C until use.

#### Family-wide hallucination of four-helix bundles

We generated forty single-chain four-helix bundle proteins using a combination of parametric design, structural bioinformatics, generative AI for loop generation, and a graph neural network for designability assessment. We first computed 750 four-chain coiled coils, with each helix consisting of 40 residues, using the CCCP server (https://grigoryanlab.org/cccp/index.gen.php) and sampling Crick parameter ranges from designable distributions seen in natural proteins. We focused on coiled coils with large supercoil radius (> 7.6 Å and < 8.0 Å) to fit small-molecule ligands in the interior. For each coiled coil, we then used the program MASTER^41^ to search for appropriate loop lengths to connect the helices, by searching a non-redundant database of the PDB with each protein separated into files containing individual, non-redundant chains (30% sequence identity). We used as a structural search query the terminal 7 residues of each helix (truncated to 28 residues each) to be connected by a loop and structural matches defined as a C⍺ RMSD < 0.8 Å. We varied the number of “gap residues” from 2 to 17 and chose the optimal loop lengths that summed to 36 residues, creating single chain proteins of 148 amino acids in length. The optimal number of 36 residues was chosen based on maximizing a score, averaged across each of the coiled coil structures, which favored many MASTER matches with low C⍺ RMSD (score = # of unique MASTER matches / <C⍺ RMSD>). We then ran RFdiffusion to generate loops with the chosen optimal lengths (which could be different lengths for different coiled coils, with the constraint that the total number of loop residues sums to 36). RFdiffusion generated 8 single-chain structures for each coiled coil. We then designed 16 sequences for each backbone using ProteinMPNN (v_48_020 model, T=0.15, no Cys allowed) and predicted their structures using AlphaFold2 (model 4, 1 recycle, single-sequence mode). We filtered the predicted structures using the following criteria: average C⍺ pLDDT > 0.9, average pAE < 5.0, C⍺ RMSD to the input structure (output from RFdiffusion) < 1.0 Å. Of these proteins, we selected designs whose input backbones to ProteinMPNN produced 8 or more (out of 16) sequences that passed these filters. Finally, we removed any designs with minimum C⍺ pLDDT of any non-terminal residue < 0.85. If 3 or more of the 16 sequences for a backbone remained after this final filtering, we added the protein to a final pool. We then clustered this final pool of designs by C⍺ RMSD (using a cutoff of 0.75 Å) after all-by-all pairwise superposition. 40 clusters were found, and we chose the centroids of each cluster as our hallucinated, highly designable ensemble to be used for design of small-molecule binders. Many of the final designs came from different parent parametric backbones (at most 5 from the same original parametric backbone with mostly singletons). For design, we directly used the AF2-predicted structures after removing the ProteinMPNN-designed sequence (by turning all residues to Gly or Ala using Rosetta). The ensemble of forty single-chain bundles, including the ProteinMPNN-designed sequences along with the Ala/Gly versions, are available on Zenodo^58^.

#### Fluorescence polarization experiments to determine binding affinities

We performed fluorescence anisotropy experiments in a 384-well plate-reader format to determine the binding affinities of proteins with various ligands (Fig. E11 and Table S2). We serially diluted an initial solution containing a high concentration of protein while keeping a constant concentration of ligand (50 nM) in each well. The buffer for these assays was composed of 1x PBS, pH 7.4, and 0.1% w/v PEG-8000. Campothecin-class molecules (exatecan, camptothecin, belotecan, FL118, irinotecan) were excited at a wavelength of 360 nm using a BMG LabTech PHERAstar FSX plate reader or a BMG LabTech CLARIOstar Plus plate reader. We detected the resulting parallel and perpendicularly polarized emission at 440 nm. Molecules containing a FITC fluorophore (apixaban-peg-FITC, dexamethasone-peg-FITC) were excited using 482 nm filter (16 nm bandwidth) using a BMG LabTech CLARIOstar Plus plate reader, and parallel and perpendicularly polarized emissions were detected through a 530 nm filter (40 nm bandwidth). Data was fit using the quadratic form of a single-site binding model. Gain settings for each molecule were calculated by auto-adjusting the focus on a well containing free ligand and buffer, setting the millipolarization (mP) of free ligand to mP = 35.

#### Fits to fluorescence anisotropy data

Data from fluorescence anisotropy (r) experiments, which varied total protein concentration, [P]T for a fixed total ligand concentration, [L]T, was fit to a single-site binding model (Equations 1 and 2, N = 1). Confidence intervals of the fit parameters were determined from bootstrapping the residuals of the optimum fit (random residual selection with replacement) to create 1000 synthetic datasets, which were each separately fit to Equations 1 and 2. Parameter means and 95% confidence intervals were determined from the distribution of the parameters computed from the bootstrapped fits.

We performed a global fit for mutant EPIC proteins, which had affinities with exatecan comparable to the ligand concentration (Fig. E16). We performed separate protein titrations across three fixed exatecan concentrations (25, 12.5, and 5 nM) and measured fluorescence anisotropy of exatecan as a function of protein concentration. For each EPIC mutant, the dissociation constant (*K*_d_) and anisotropy baseline (A_min_) were shared parameters across the three exatecan concentrations. Anisotropy maxima (A_max_) and ligand concentration ([L]_T_) were fit locally. We fit the ligand concentrations locally (to within 15% of the target concentration) to account for experimental discrepancies. Code for performing the fitting is provided in Zenodo^58^.

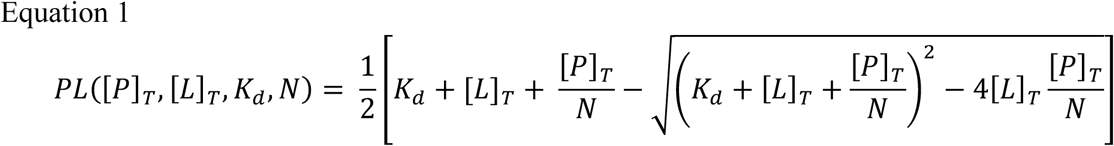

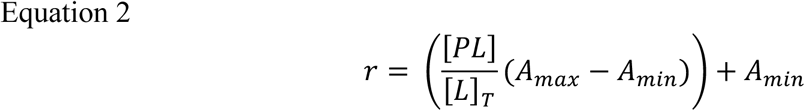

#### Cloning, protein expression, and protein purification

##### Cloning

Genes were ordered from Integrated DNA Technologies (IDT) and cloned into a peAIP32 plasmid using a Golden Gate reaction (cloning site BsaI). The plasmid also included a 6x His tag followed by a Cth Sumo protease cleavage sequence. We cloned the plasmid resulting from the Golden Gate reaction directly into competent BL21 *E. Coli* prepared with the Zymo Research Mix-n’-Go kit. Cells were then plated onto LB/kanamycin, and single colonies were selected for whole-plasmid nanopore sequencing (Quintara Biosciences). Once cells were verified to contain the correct plasmid, glycerol stocks were created and used for induction of subsequent growths.

##### Protein Expression

Growths (0.5 L each) were carried out in LB/kanamycin media containing 34.7 mM lactose for autoinduction. Cultures were allowed to grow overnight, for at least 16 hours, at 30 C shaking at 220 rpm.

##### Protein Purification

Following overnight growth, cells were centrifuged. Liquid media was discarded, and cell pellets were immediately resuspended in lysis buffer (PBS, 500 mM NaCl) then directly lysed on ice by sonication. Cell debris was removed by centrifugation at 16000 rpm (11600 x g) for 30 minutes. Protein was purified using magnetic Ni NTA beads (GenScript). A total of 1.5 mL of beads was incubated with each 0.5 L growth for at least an hour at 4 °C. The slurry was then added to a gravity column, and the flow-through was discarded. Washes performed included one 15 mL wash with wash buffer (PBS, 500 mM NaCl, 20 mM imidazole) and one 10 mL wash with PBS. The beads were then allowed to shake overnight at 4 C with Cth protease (1 µM) to cleave designed protein from the beads. Elution containing the purified cleaved protein was collected using a gravity column. The beads were washed with one column volume equivalent (1.5 mL) of PBS, which was collected and pooled with the elution. The resulting protein was stored in PBS buffer at 4 °C until use in experiments.

#### Size exclusion chromatography

Size exclusion chromatography (SEC) was carried out to determine the oligomerization state of each construct using a BioRad NGC system paired with a BioRad ENrich 650 chromatography column. Each protein eluted at column volumes that approximately matched their molecular weights (Figs. E10 and E18). Any constructs that were determined to have a minor oligomeric peak (only applicable to the COMBS/Rosetta designs) were purified via fractionation using the BioRad ENrich 650 column to isolate the major peak corresponding to the monomer.

#### Circular Dichroism to determine secondary structure and stability

We used circular dichroism (CD) spectroscopy to determine the secondary structure content and thermal stability of EPIC (Fig. E14). A sample of 0.1 mg/mL EPIC protein was purified by SEC into 1x PBS buffer. CD spectra were collected using a 0.1 cm path-length quartz cuvette (VWR) in a Jasco J-1500 CD Spectropolarimeter with temperature controlled at 25 °C using a Peltier temperature controller. Spectra were collected from 195 nm to 260 nm in continuous scanning mode, with a band width of 1 nm, scanning speed of 50 nm/min, data pitch of 0.1 nm, digital integration time of 1 second, and an average of 5 accumulations. Temperature-dependent spectra were similarly collected using a Peltier temperature controller to vary the sample temperature in the range of 25 °C to 95 °C with an interval of 5 °C, an increase rate of 10 °C/minute, and an average of 3 accumulations.

#### Hydrolysis experiments

Exatecan has a labile lactone ring that hydrolyzes (opens) to a carboxylate at physiological pH. We spectroscopically monitored lactone ring opening by collecting time-dependent absorption spectra using an Agilent 8453 G1103A spectrophotometer and a 1 cm path-length quartz cuvette. Samples in a PBS solution (pH 7.4) with a final volume of 2 mL included 20 µM protein and 20 µM ligand (final concentration of DMSO at < 0.54%). Spectra between 190 and 900 nm were collected every half hour until spectral changes were no longer observed. For exatecan with EPIC^Q51N/M97L^, we measured spectra until 49 hrs and observed no significant change in the absorbance spectrum of exatecan.

#### Binding to HSA is compatible with hydrolysis

To simulate conditions in human plasma, we tracked hydrolysis of exatecan via a time-resolved absorption experiment with a mixture of EPIC^Q51N/M97L^ and human serum albumin (HSA). We first mixed EPIC (40 µM) and exatecan (40 uM) in PBS for a total volume of 1 mL and allowed the solution to equilibrate at room temperature for an hour. We then combined the solution with 1 mL of HSA (1 mM) in PBS, yielding a final solution containing 20 uM EPIC^Q51N/M97L^, 20 uM exatecan, and 500 µM HSA. Absorbance spectra were collected every half hour using an Agilent 8453 G1103A spectrophotometer with a 3 mL 1 cm path-length quartz cuvette. A scattering component grew over time, presumably due to oxidation of the highly concentrated HSA. The scattering component was also observed in experiments with 500 µM HSA alone.

#### Analysis of exatecan hydrolysis data

Absorption spectra of exatecan and exatecan–protein mixtures were collected at various time delays after initial mixing from a DMSO stock of ring-closed exatecan (Fig. E22). Spectra were globally fit to a two-state kinetic model that solves for both the forward and backward rate constants (ring-closed to open states) as well as the corresponding species-associated spectra (Fig. E23). Samples that included HSA showed a time dependent increase in scattering, which we could mostly remove by fitting a polynomial to the absorption spectrum between 450 and 850 nm. For consistency, we applied this baseline procedure to each dataset before fitting the kinetic model. Fits to the time-resolved absorbance data were generally very good using a two-state model (small and normally distributed residuals). Because there was little time evolution observed for exatecan when mixed with EPIC, the two species-associated spectra derived from the fit were nearly identical. We therefore constrained the fits to obey a final equilibrium distribution of 90% ring-closed for EPIC, computed from using a thermodynamic model that incorporated the measured *K*_d_ of EPIC for exatecan. Using the equilibrium constraint, the resulting species-associated spectra more closely resembled those of ring-open and ring-closed observed from fits of exatecan only and exatecan with HSA. A single-state model with no time evolution (initialized in 100% ring-closed state of exatecan) was sufficient to model the data of exatecan with EPIC^Q51N/M97L^. A yield of < 1% for the ring-open form of exatecan was computed from a thermodynamic model that incorporated the measured *K*_d_ of EPIC^Q51N/M97L^ with exatecan. Code for performing the fitting analysis is provided in Zenodo^58^.

#### Design of exatecan binders via COMBS and Rosetta

We used an advanced version of COMBS (code provided), together with the two exatecan conformers and the 40 four-helix bundles, to simultaneously dock exatecan and design ligand-binding sites. COMBS uses van der Mers (vdMs) to first place fragments in favorable positions with respect to an input backbone. COMBS then docks the ligand to overlay these fragments with low heavy-atom RMSD. COMBS then uses a recursive H-bonding algorithm that treats polar, first-shell (i.e. contacting the ligand) sidechains as new “ligands” and finds second- and (sometimes) third-shell interactions that support the binding site in an H-bond network. The entire extended binding site is optimized using a statistical vdM energy function that considers steric clashing between sidechains and pairwise interactions modeled by vdMs. We also used a custom ligand constraint file in COMBS to only consider ligand positions that buried the lactone ring of exatecan and positioned the charged amine close to the protein surface. We additionally filtered any exatecan ligands that did not have at least 75% of apolar heavy atoms buried in the protein core (inside the convex hull created by the C⍺ atoms of the backbone). We also required that polar atoms of exatecan had an H-bonding interaction if the atoms were buried in the protein core. Lastly, we required that the exatecan A, B, or C rings (see chemical structure in Table S2 for ring labels) had at least one van der Waals contact from a designed residue. For each of the forty backbones, we used COMBS to output 75 designs with high variability in binding-site sequence and ligand location. We pooled these designs and selected 100 candidates based on COMBS statistical vdM energy and the number of buried polar atoms not participating in an H-bonding interaction. We then used a custom flexible backbone design algorithm implemented in Rosetta to design the remainder of the protein sequence. For this, the binding-site residue identities were frozen and designed H-bonding distances were constrained using a Rosetta constraint file. For each of the selected COMBS designs, we used Rosetta to fill in the remainder of the sequence, outputting 100 sequences each (which shared the same COMBS-derived residues). We then used AF2 and RFAA to predict the unbound and ligand-bound structures of each of these designs (approximately 10,000 sequences), selecting designs for experimental interrogation by the following criteria: ligand heavy-atom RMSD < 2.0 Å, ligand heavy-atom RMSD over the ring system < 1.5 Å, binding-site C⍺ RMSD < 0.9 Å (RFAA, aligning on binding-site residues only, defined as those with heavy atoms within 5 Å of the ligand), overall C⍺ RMSD < 1.0 Å (RFAA, aligning over all residues), average C⍺ pLDDT > 0.80 (RFAA), average C⍺ pLDDT > 0.90 (AF2), average protein pAE < 5.0 (RFAA), average protein pAE < 5.0 (AF2), binding-site C⍺ RMSD < 0.9 Å (AF2, aligning on binding-site residues only), overall C⍺ RMSD < 1.0 Å (AF2, aligning over all residues), number of buried unsatisfied atoms computed by Rosetta (BuriedUnsatHbonds filter) < 2, minimum pLDDT (AF2) of any non-terminal residue > 0.7, and average interfacial pAE > 15 (AF2 multimer, two copies of the sequence). After this filtering, 64 designs remained. We grouped them by those that shared the same input to Rosetta and ranked designs by a heuristic score that favors low C⍺ RMSD over the binding-site residues [score = 0.5 ligand heavy-atom RMSD + 1.0 binding-site C⍺ RMSD (RFAA) + 0.75 binding-site C⍺ RMSD (AF2) + 0.25 overall C⍺ RMSD (RFAA) + 0.25 overall C⍺ RMSD (AF2)]. We chose the top 3 (lowest scoring) designs from each group, leading to 24 designs overall. We ordered synthesized DNA for these sequences from IDT (e-block) and experimentally tested 16 of them. We subsequently ordered two additional designs for the input pose to NISE, which we selected by score from the 100 outputs of Rosetta that used the same COMBS input. The computational models of the designs are available in Zenodo^58^.

#### Initial COMBS model for NISE design

The initial model for the input to the NISE design algorithm, CR4, was chosen based on the heuristic score described above. The input to NISE was the COMBS/Rosetta design that had the lowest heuristic score. It was also one of the top-ranked designs when using a strategy involving LASErMPNN design to design 5 sequences and co-structure prediction (and ranking by ligand pLDDT or ligand heavy-atom RMSD) (Fig. E13). Size exclusion chromatography showed that CR4 formed a higher order oligomer when expressed and purified from *E. coli*. While CR4 showed an increase in fluorescence polarization when mixed with exatecan, the data could not be fit to a single-site binding model, presumably due to the oligomeric nature of the protein.

#### Comparison of input COMBS model vs EPIC design

The model used as input to NISE (CR4) was designed using van der Mers (vdMs) to both dock the ligand and establish many of the binding interactions, including a second-shell His-Glu interaction (Fig. E9). Residues derived from vdMs were frozen, and the remainder of the sequence was designed using a flexible-backbone design loop in Rosetta. The CR4 model was ranked the highest among the 16 models designed by COMBS and Rosetta when applying a heuristic self-consistency score. Although CR4 aggregated, it showed some signs of binding exatecan via fluorescence polarization experiments, but the fits were not consistent with a single-site model. A similar pose from traditional COMBS/Rosetta design, CR3, which used the same vdMs of the binding site, was monomeric and bound exatecan with an 8 µM dissociation constant. The NISE protocol not only resulted in sequences that were monomeric, but each of the four tested NISE designs bound exatecan, with EPIC binding the tightest (*K*_d_ = 0.12 µM). The EPIC binding site differed starkly from that of CR4 (Fig. E9). EPIC used fewer polar residues and more aliphatic residues, which still engaging with key polar groups of exatecan, such as the hydroxyl and amine.

#### Selection of additional pair of COMBS designs for experimental characterization

Because each of the four NISE designs bound exatecan, we sought to test more of the traditional designs that shared the same pose (backbone and ligand position) as the input to NISE. Two of the sixteen traditional designs share the same pose, one of which (CR4) aggregated and the other (CR3) bound exatecan with *K*_d_ = 8 µM. These two designs were the top 2 of 100 COMBS/Rosetta designs when ranked by a heuristic self-consistency score (see above). We selected the next two highest ranking designs from the remaining 98 using the same scoring function after applying the same filters described in section Design of exatecan binders via COMBS and Rosetta. These designs are CR5 and CR6 (Table S3).

#### Ranking of COMBS designs by metrics from protein–ligand co-structure prediction

We used RFAA, Boltz-1, and AF3 to predict structures of the 16 designs from COMBS and Rosetta (see Table S3 for sequences). Both ligand confidence (pLDDT) and ligand self-consistency (heavy-atom RMSD) could be used to rank binders over non-binders (Fig. E12). RFAA ligand pLDDT and AF3 ligand RMSD showed the best ranking performance.

#### Ranking of designable poses by metrics from protein–ligand co-structure prediction

We wondered if the most promising poses could be prioritized for input to NISE through a rapid computational screen using LASErMPNN and a structure-prediction method. To do this, we used the computational models of the 16 experimentally characterized COMBS/Rosetta sequences. For each COMBS/Rosetta design, we removed the sequence information and used only the backbone and ligand coordinates (pose) as input to LASErMPNN. If the COMBS/Rosetta design bound exatecan, we designated the pose as a binder. We designed 5 sequences using LASErMPNN and predicted their exatecan-bound structures using RFAA, Boltz-1, and AF3. AF3 showed the best performance, with ligand pLDDT and ligand heavy-atom RMSD both correlating with binding (Fig. E13). Both AF3 and Boltz-1 showed enrichment for binders when ranked by ligand RMSD after first filtering designs that had low self-consistency (Ca RMSD or ligand heavy-atom RMSD > 2 Å, Fig. E13b,c). In contrast to results using the designed COMBS/Rosetta sequences (Fig. E12), metrics from RFAA showed the worst performance, as neither variable correlated with binding (not shown). These results show that LASErMPNN and AF3/Boltz-1 can feasibly be used to rapidly screen for promising poses (ligands docked to a backbone by a variety of methods) as input into the more computationally intensive NISE algorithm for optimization.

#### X-ray crystal structures

EPIC and EPIC^Q51N^ were purified as described above. EPIC solutions were diluted so that exatecan from a DMSO stock (3.76 mM) could be added at a 1:1 molar ratio while keeping the total concentration of DMSO < 2.6% (to about 100 µM). After addition of exatecan, EPIC-exatecan solution was concentrated to 3.5 mM using 3k centrifugal filters (Millipore). We prepared the solution for co-crystallization at room temperature using the sitting drop vapor method. Crystallization conditions screened included the sparse matrix screens MCSG-1, MCSG-2, MCSG-3, MCSG-4, PEG/Ion, Index, and JCSG Plus. EPIC crystals obtained from 7 µL EPIC-exatecan solution combined with 7 µL of MCSG-3 well D6 solution (0.2 M Zinc Acetate, 0.1 M MES: NaOH, pH 6, 10 % (w/v) PEG 8000) diffracted to a resolution of 2.0 Å. EPIC^Q51N^ crystals obtained from 7 µL EPIC-exatecan solution combined with 7 µL of MCSG-3 well A2 solution (0.1 M Sodium Phosphate Dibasic: Citric Acid, pH 4.2, 2 M Ammonium Sulfate) diffracted to a resolution of 2.2 Å.

Data was collected on a DECTRIS EIGER X 9M detector at the 17-ID-1 (AMX) beamline at the National Synchrotron Light Source II of Brookhaven National Laboratory. Raw images were indexed and merged using autoPROC^59^, and the structures were phased via molecular replacement in Phenix^60^ using the design models of EPIC and EPIC^Q51N^. The models were then refined in Phenix and Coot^61^. The composite-omit maps shown in the main article were produced in Phenix (version 1.20.1-4487-000, default parameters of Omit map method: refine). Table 1 statistics (Tables S4 and S5) were produced with the same version of Phenix.

#### Runtime Quantification

Running in CPU inference mode with 32 CPU cores, LASErMPNN generates 100 designs for a 148-residue protein in around 50 sec. On a NVIDIA A4500 GPU, with the same inference settings, 100 designs can be generated in around 34 sec. One NVIDIA A4500 can run an iteration of the NISE algorithm over 1500 sampled sequences (500 sequences per backbone) using RoseTTAFold-All Atom with 15 recycles in around 6 hr for a 148-residue protein. Running an analogous NISE loop with Boltz-1 (default inference settings) on a single NVIDIA A4500 similarly takes around 6 hr. Using Boltz-1, we noticed that the number of sequences sampled per NISE iteration can be reduced to around 150 sequences per iteration (50 sequences per backbone) to observe similar quality sequence/structure optimization taking around 40 min per iteration on one NVIDIA A4500. Boltz-1 is more successful at predicting self-consistent structures (Figure E2, E5, E25) and is therefore more efficient for NISE optimization. The main time bottleneck in NISE is the structure prediction step, so we used between 4 and 8 GPUs to parallelize the structure-prediction computations while developing the algorithm. Further time savings can be made by reducing the number of recycles or diffusion steps (where applicable) used by the structure-prediction networks.

## Data Availability

Coordinates and data files for X-ray crystal structures of exatecan-bound EPIC and exatecan-bound EPIC^Q51N^ have been deposited in the PDB with accession codes 9NZE (EPIC) and 9NZG (EPIC^Q51N^).

## Code Availability

All model training code, processed datasets, and dataset split information can be found at https://github.com/polizzilab/LASErMPNN. An implementation of the NISE protocol using Boltz-1 can be found at https://github.com/polizzilab/NISE.

## Acknowledgements

N.F.P acknowledges funding from NIH (R00GM135519) and the Innovation Research Fund of Dana-Farber Cancer Institute to support this work. N.F.P. and B.F. thank SBGrid for support of on-site compute infrastructure and crystallography software. N.F.P. and K.S. thank John Alberta for help with equipment and resources for experiments. K.S. thanks Roksana Azad for help with setting up crystal trays for X-ray crystallography. N.F.P. and K.S. thank AI Proteins for sharing sumo-tag and sumo-protease expression plasmids for protein expression and purification, respectively. N.F.P. thanks Eric S. Fischer for helpful comments on the manuscript. This research used resource 17-ID-1 of the National Synchrotron Light Source II, a U.S. Department of Energy (DOE) Office of Science User Facility operated for the DOE Office of Science by Brookhaven National Laboratory under Contract No. DE-SC0012704. B.F. acknowledges funding from the National Science Foundation Graduate Research Fellowship Program.

## Author contributions

B.F. performed the computations and trained LASErMPNN. K.S. performed the experiments. B.F., K.S., and N.F.P. performed analysis. B.F., K.S., and N.F.P. wrote the paper.

## Materials & Correspondence

Correspondence and request for materials should be addressed to Nicholas Polizzi.

## Extended Data

See supplementary information.

---

**Fig. E1.**
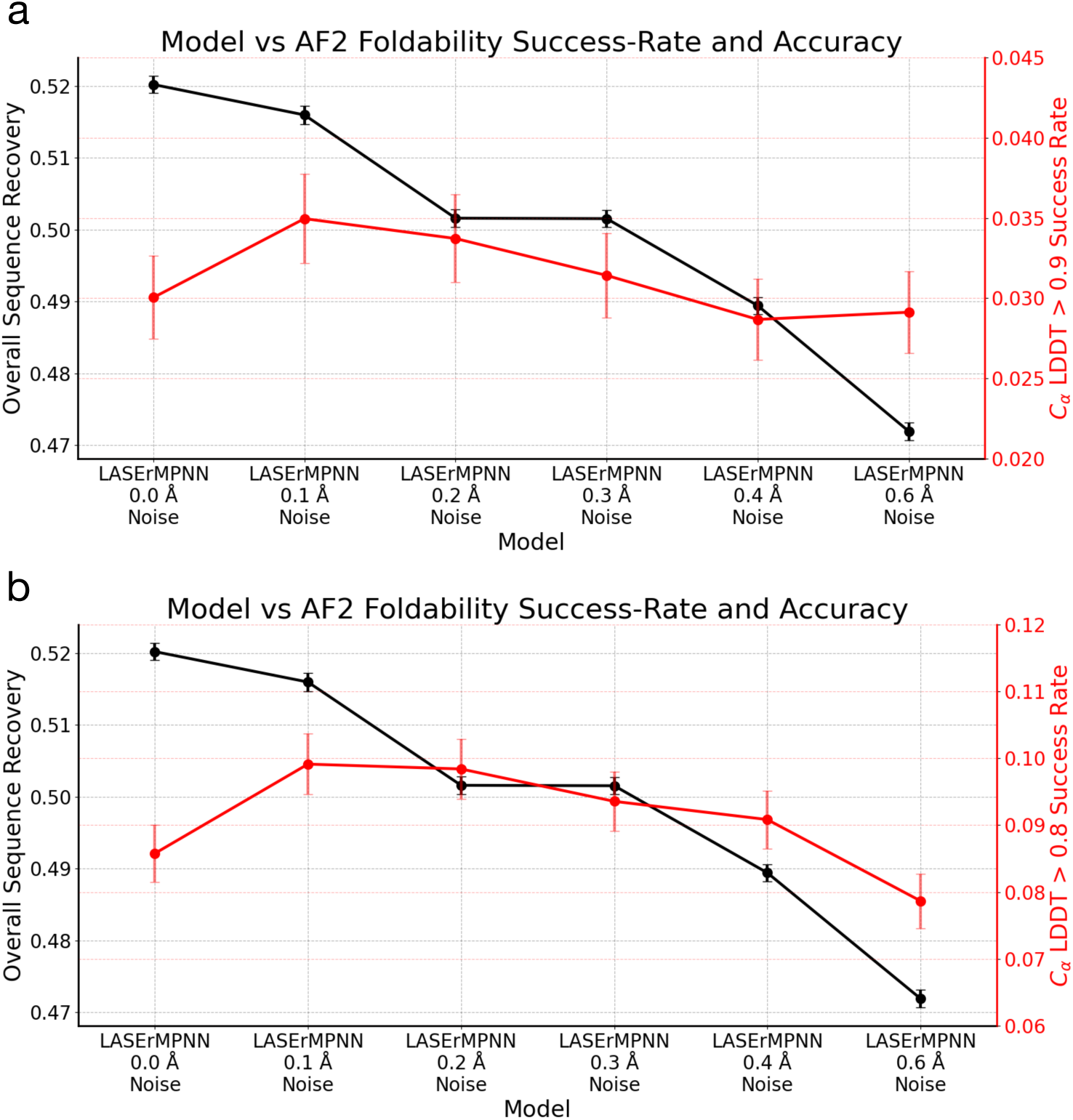
Using AF2 foldability to identify optimal amount of training noise. **a,** Similar to the approach taken with ProteinMPNN, we monitor, as a function of training noise level, the fraction of sequences designed by LASErMPNN that have C⍺ local distance difference test (LDDT) over a certain threshold. This process identified 0.1 Å as an optimum noise level on residue frames during training that maximizes overall sequence recovery and the fraction of a monomeric validation protein set (N=882) predicted by AF2 with C⍺ LDDT > 0.9. **b,** We repeat this analysis with a success rate threshold of 0.8 which also identifies 0.1 Å as an optimum maximizing sequence recovery and foldability. The presented values and uncertainties are means and standard deviations computed by bootstrapping (N=250,000) the respective metric uniformly over the 5 sampled sequences for each validation protein.

**Fig. E2.**
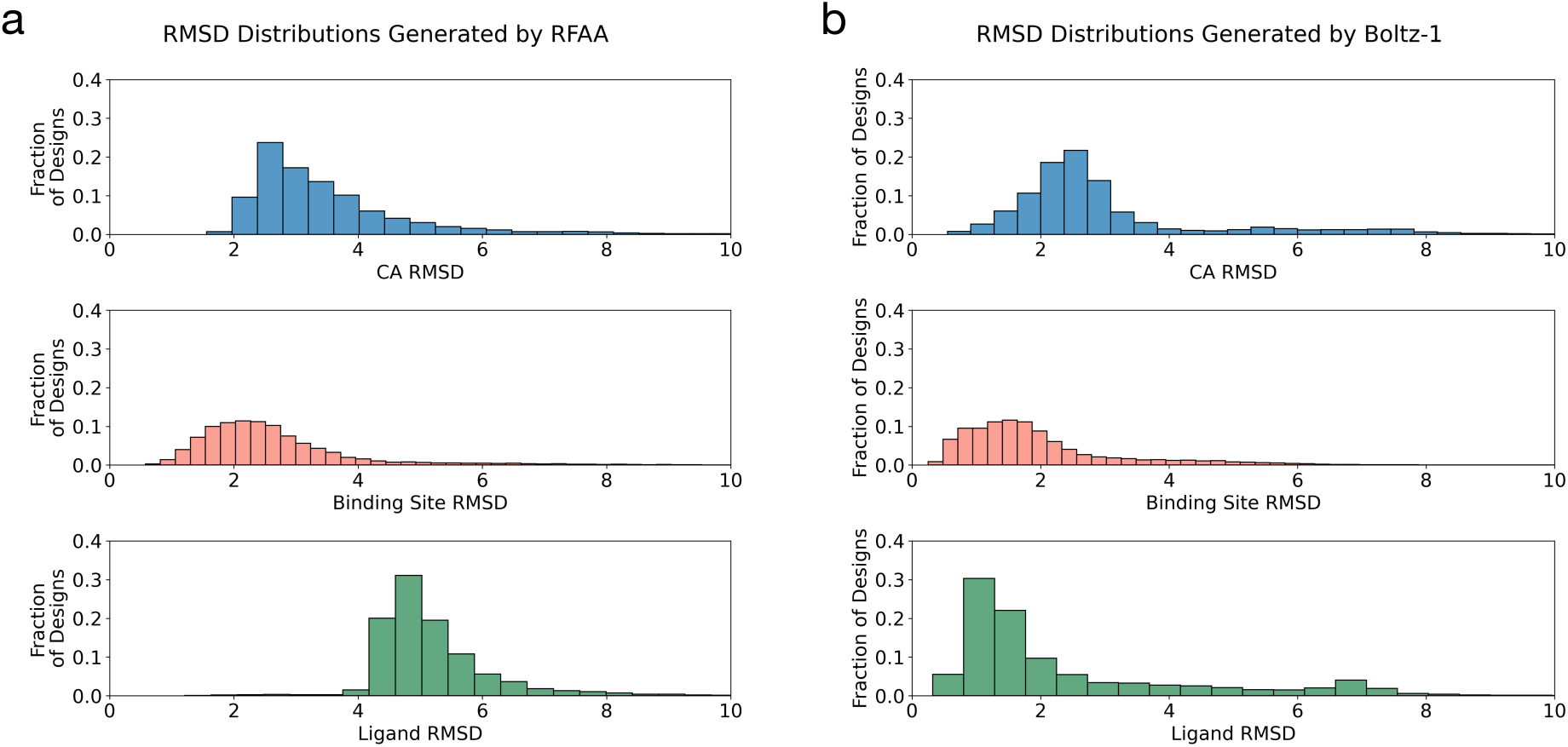
Distributions of various RMSD metrics for LASErMPNN-designed sequences of streptavidin-biotin (pdb 4jnj), using predicted protein–ligand co-structures from RFAA or Boltz-1. The same designed sequences were folded using RFAA (**a**) and Boltz-1 (**b**). CA RMSD refers to the RMSD of all protein backbone C⍺ atoms after superposition onto the input 4jnj structure. Binding Site RMSD refers to the RMSD of protein backbone C⍺ atoms of residues with heavy atoms within 5.0 Å of biotin in the input crystal structure. Ligand RMSD refers to RMSD taken over the ligand heavy atoms after superposition on binding-site residues.

**Fig. E3.**
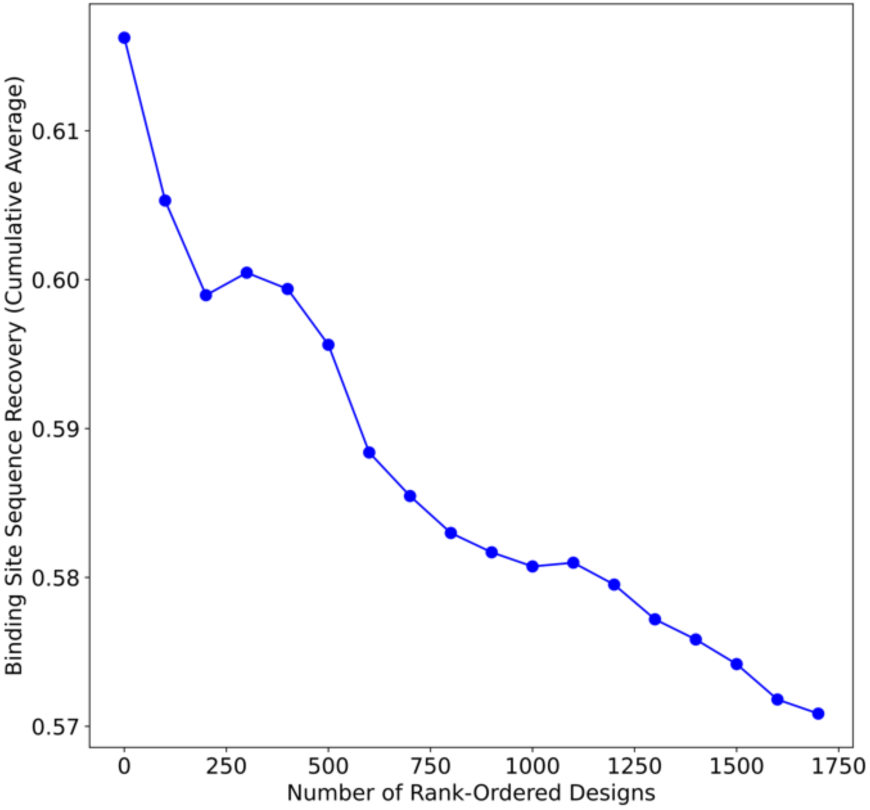
Binding-site sequence recovery is inversely correlated with the number of buried, non-H-bonded, polar atoms. Data shown is an analysis from 10,000 designed sequences of monomeric streptavidin and biotin complex (pdb 4jnj). Structures were predicted by Boltz-1 for each of the sequences designed by LASErMPNN, which were then energy-minimized (backbone and sidechains) using Rosetta (Ref15 energy function). We then computed the number of buried, non-H-bonded (“unsatisfied”) polar atoms (BUNs) of the ligand and first-shell interacting residues. We sorted the designs by a formula that is 2x first-shell BUNs + 1x ligand BUNs. Then we compute a cumulative average of binding-site sequence recovery after inclusion of the next-best (by BUNs) 100 designs (circles). The plot shows that average binding-site sequence recovery is highest for the top-ranked designs by BUNs and decreases as number of BUNs increases.

**Fig. E4.**
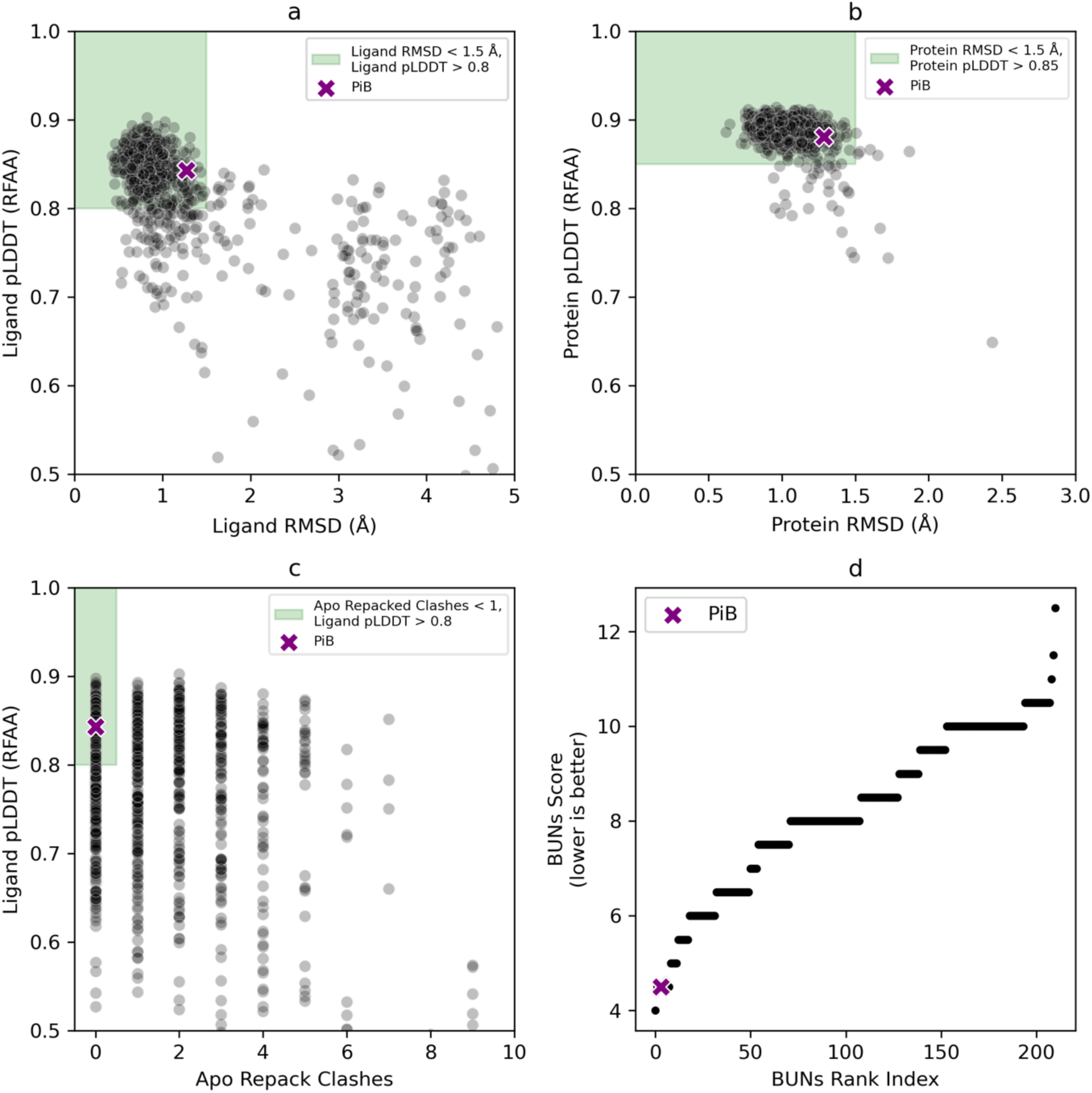
Filtering and ranking protocol of LASErMPNN-designed sequences ranks PiB favorably. We used LASErMPNN to design 1000 sequences for the rucaparib-bound PiB helical bundle (pdb 8tn6). The sequence of PiB (< 5 nM dissociation constant with rucaparib) was added to the set of LASErMPNN-design sequences and is denoted with a purple “x” in each plot. Sequences were filtered by **a,** ligand heavy-atom RMSD < 1.5 Å and ligand pLDDT > 0.8, **b,** protein C⍺ RMSD < 1.5 Å and protein C⍺ pLDDT > 0.85. **c,** We performed a calculation that approximates the protein’s preorganization for binding, by computing sidechain clashes with the ligand after binding-site sidechains were repacked in the absence of the ligand. We selected designs with no clashes. **d,** We ranked remaining designs by BUNs Score (2x ligand BUNs + 1x protein BUNs). The PiB sequence ranks 4^th^ out of the total 1001 sequences (using a subsequent ranking by ligand pLDDT to break degeneracy of designs with equal BUNs Score).

**Fig. E5.**
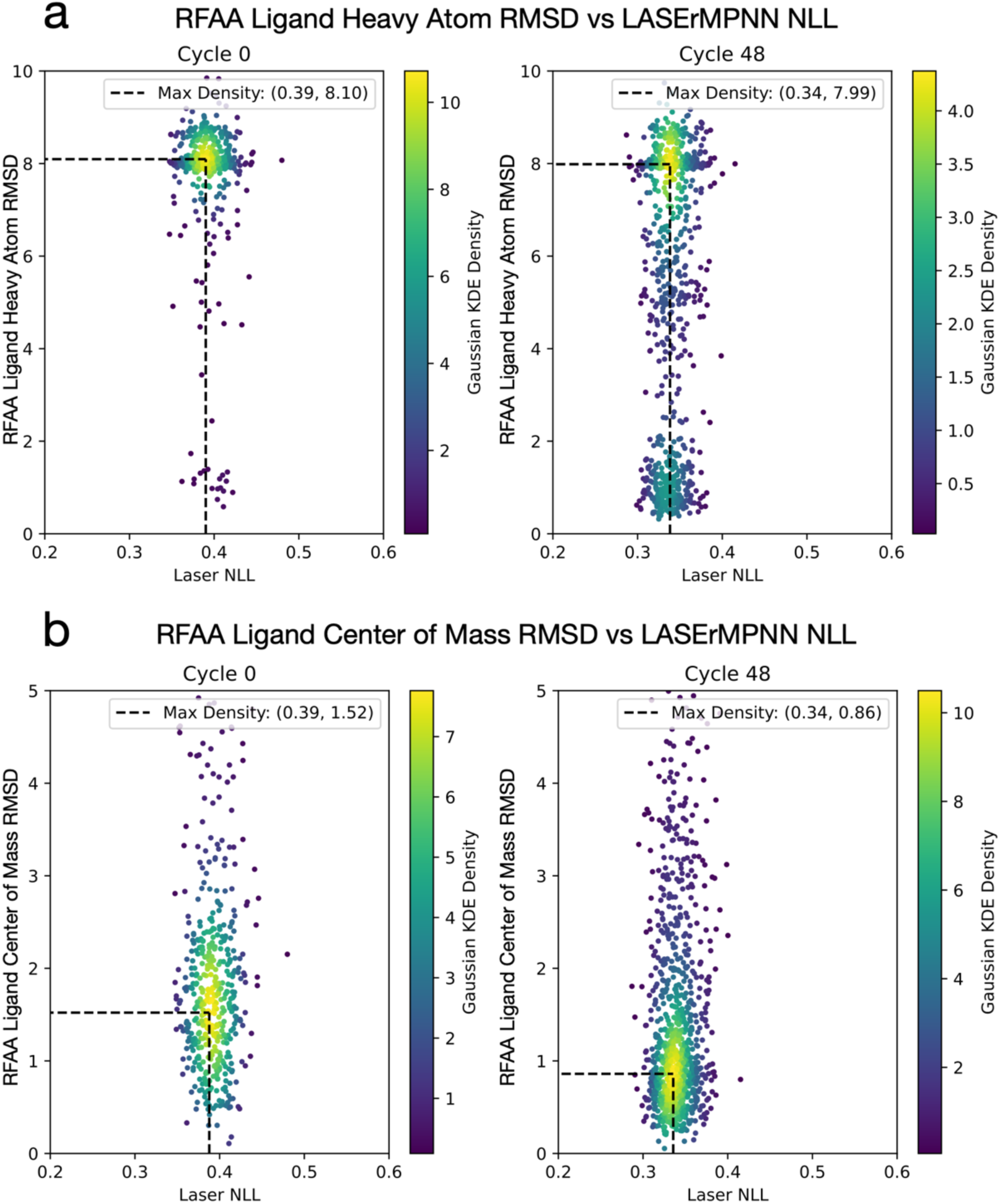
NISE improves the self-consistency of the predicted ligand and the negative log likehood (NLL) of designed sequences. We ran the NISE protocol on the CR4 pose with exatecan (all sequence information was removed). We plot ligand heavy-atom and center-of-mass RMSD vs NLL of the designed sequences, computed for the first and a later iteration of NISE. NISE improves both the ligand RMSD and NLL, shown by the distribution shift to lower values for each variable at the later iteration (right side). **a,** Ligand heavy atom RMSD (Å) between input to LASErMPNN and RFAA-predicted structure vs NLL of LASErMPNN designed sequences for first NISE iteration and the 48^th^ NISE iteration. **b,** Ligand center of mass RMSD between input to LASErMPNN and RFAA predicted structure vs NLL of LASErMPNN designed sequences for first NISE iteration and 48^th^ NISE iteration. Throughout the trajectory, RFAA is generally able to place exatecan’s center of mass within 2.0 Å of the input but struggles to predict the correct orientation of exatecan (high heavy-atom RMSD). However, the NISE protocol is able to increase the number of RFAA-predicted structures with the correct ligand orientation (low ligand heavy-atom RMSD).

**Fig. E6.**
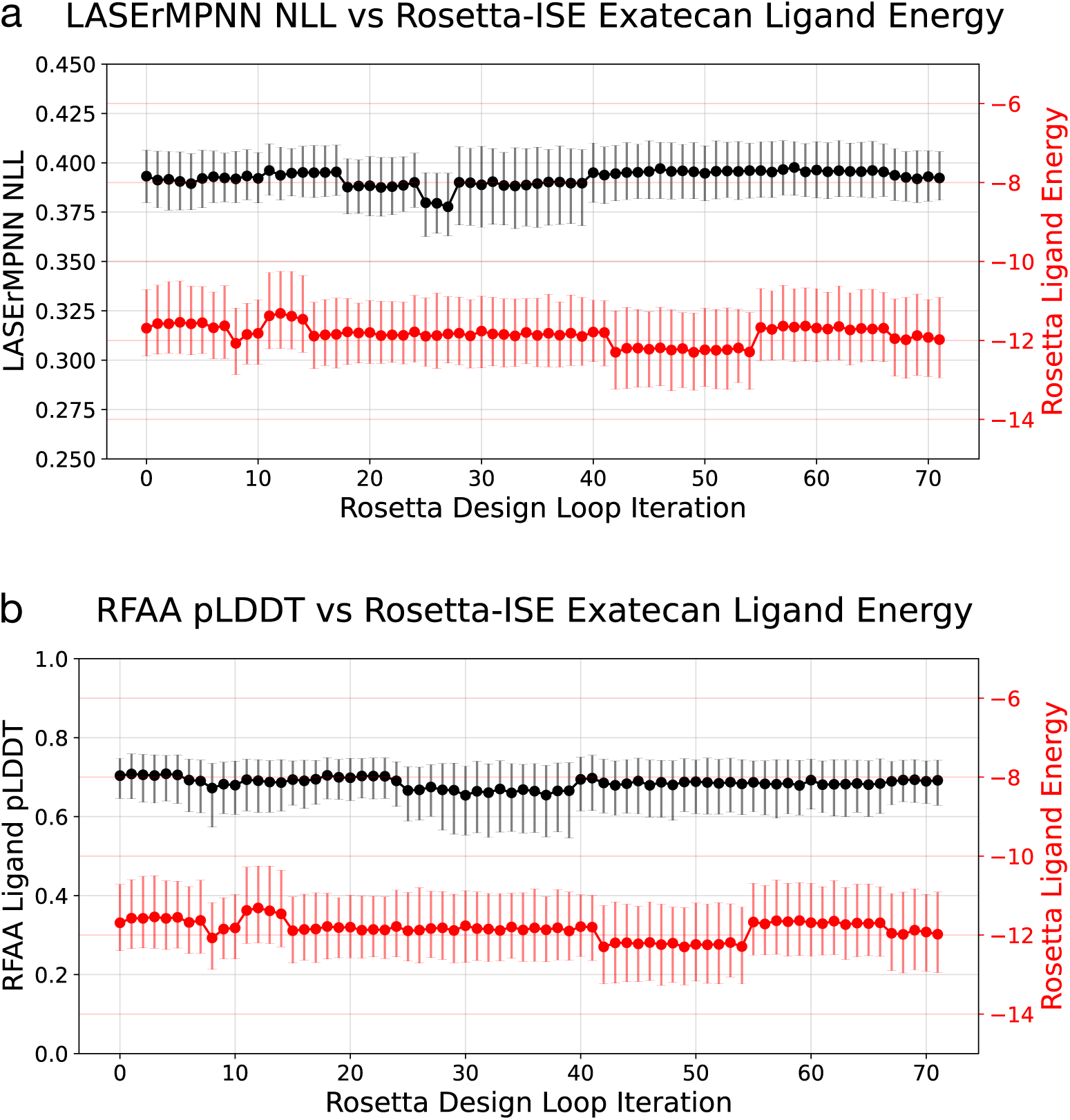
Energy-based selection-expansion is unable to optimize sequence NLL or ligand pLDDT. We ran an iterative selection-expansion (ISE) protocol using Rosetta instead of RFAA to sample new backbones from the LASErMPNN-designed sequences. We selected the top 3 Rosetta-minimized backbones by the Rosetta energy of the ligand. **a,** Median NLL of sequences and median Rosetta ligand energy as a function of ISE design cycle number. **b,** After we ran the entire trajectory (which did not use RFAA), we used RFAA to predict the structures of the sequences designed during each round. We plot the median ligand pLDDT from RFAA for these sequences and the median Rosetta ligand energy as a function of ISE design cycle number. Error bars represent first and third quartiles. N = 1500 sequences per loop iteration.

**Fig. E7.**
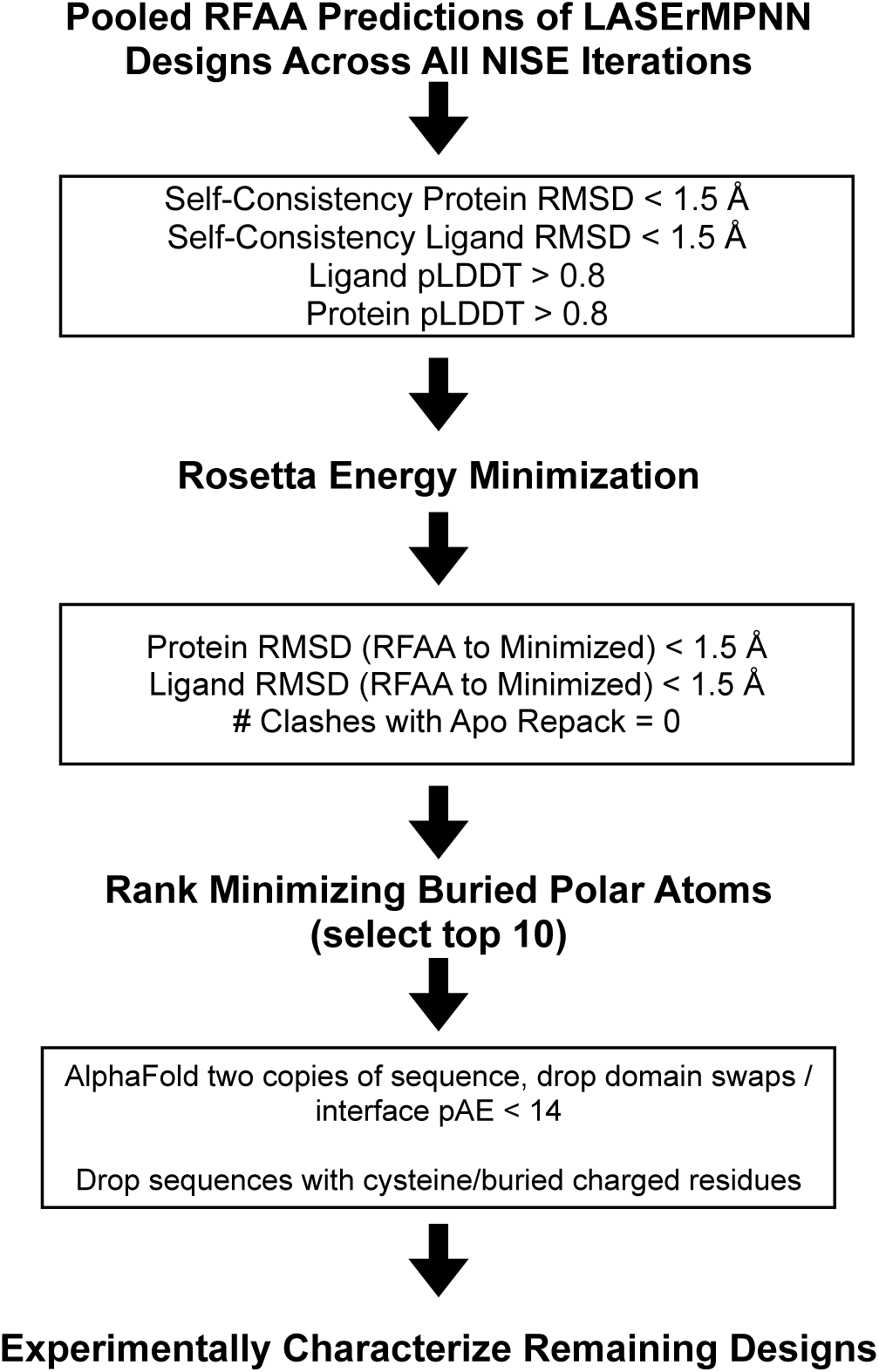
NISE filtering and ranking protocol for selecting exatecan-binding proteins for experimental characterization. We pooled the designed sequences across all NISE iterations and used the RFAA-predicted co-structures to filter out designs with low self-consistency or low confidence. The remaining sequences were energy minimized using Rosetta, and additional RMSD filters were applied that compared the structures before and after minimization. For remaining structures, we checked for preorganization by performing energy minimization of the protein in the absence of the ligand. We then re-introduce the ligand and check if any relaxed sidechains in the binding site now clash with the ligand. Sequences with no clashes were then ranked by the number of buried, non-H-bonded polar atoms of the ligand. The top 10 ranked designs were finally filtered to remove sequences with buried charged or cysteine residues and for domain-swapping or aggregation propensity using AlphaFold.

**Fig. E8.**
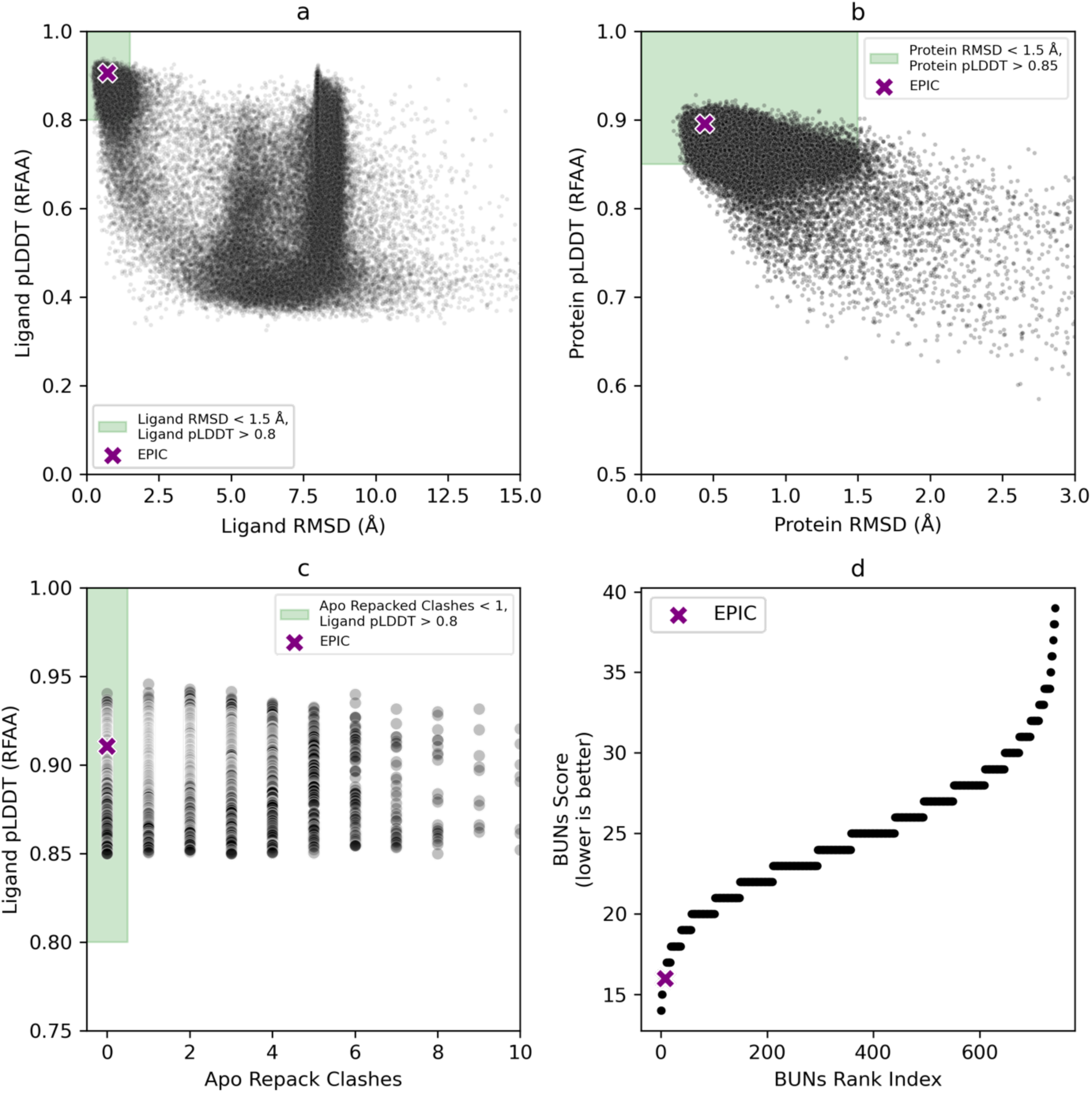
NISE filtering and ranking protocol for selection of exatecan binders. **a**, **b**, LASErMPNN-designed sequences were filtered by ligand heavy-atom and protein C⍺ self-consistency RMSD < 1.5 Å, ligand pLDDT > 0.8, and protein pLDDT > 0.85. **c**, We repacked the unbound proteins and then computed clashes with the re-introduced ligand, selecting designs with no clashes (green). **d**, We ranked remaining designs by the number of buried non-H-bonded polar atoms (BUNs) in the ligand, which places EPIC (purple “x”) as 8^th^ out of the total 72,501 sequences (using ligand pLDDT to break degeneracy of designed with equal BUNs).

**Fig. E9.**
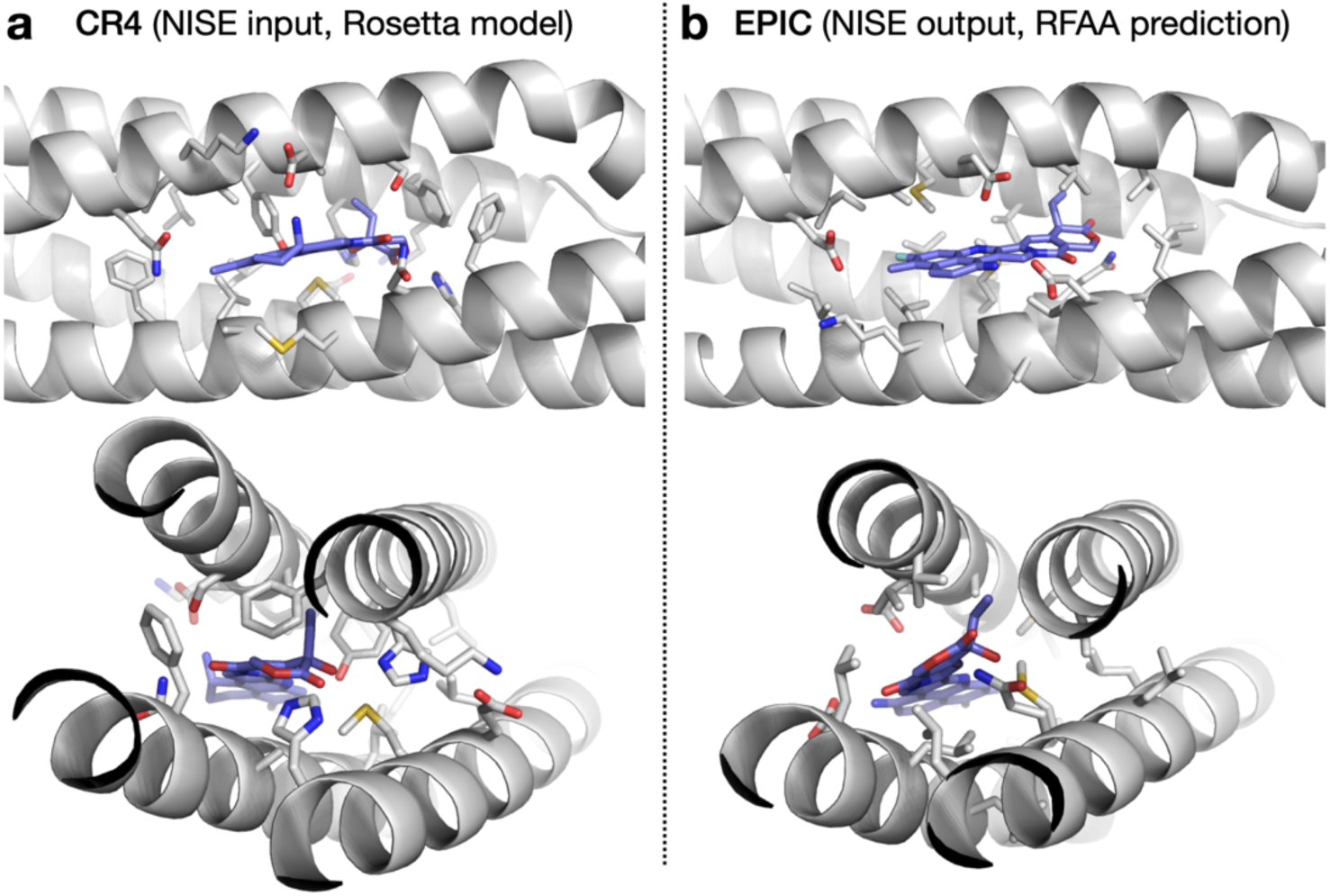
Comparison of traditional design model (COMBS and Rosetta) with output from NISE (EPIC model). **a**, Rosetta model of an exatecan-binding protein, CR4, that was designed with a traditional approach using COMBS and Rosetta. This model, after its sequence was removed, was input (backbone and ligand coordinates only) to the NISE protocol that lead to EPIC. Exatecan is shown in blue. Sidechains near the binding site are shown as stick representation. CR4 is designed to engage with exatecan using two polar His residues and Tyr, Asn, Ser, Asp residues. Aromatic Phe residues also surround the ligand. **b**, RFAA-predicted exatecan-protein co-structure of EPIC, which was designed via NISE (iterative selection-expansion using LASErMPNN and RFAA). LASErMPNN designed a binding site of EPIC that is less polar than CR4 but still engages the ligand with H-bonds from Gln51 and Asp132. Packing around the ligand is more extensive than in CR4 and uses smaller aliphatic sidechains such as Val, Ile, and Leu. The structures were aligned via their C⍺ coordinates (1.4 Å RMSD), and the views shown are from the same vantage point after alignment.

**Fig. E10.**
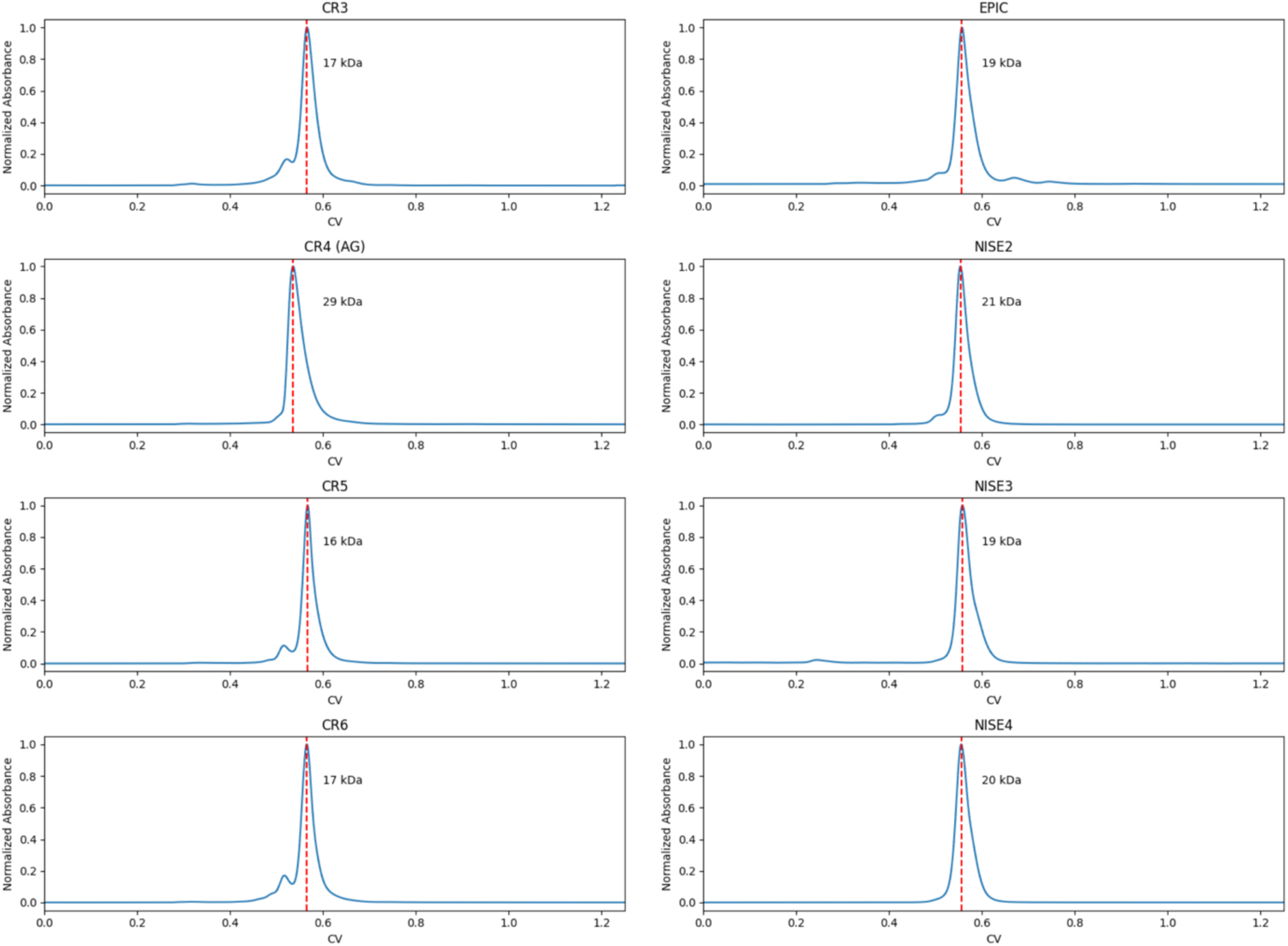
Size exclusion chromatograms of various designs. Purified COMBS/Rosetta-(left) and NISE-generated (right) protein constructs were injected on a Biorad NGC system at a concentration of > 100 µM and run on a Biorad 650 ENrich chromatography column in PBS pH 7.4 buffer. The proteins run slightly larger than the expected molecular weight (∼17 kDa), but the results are consistent with monomers.

**Fig. E11.**
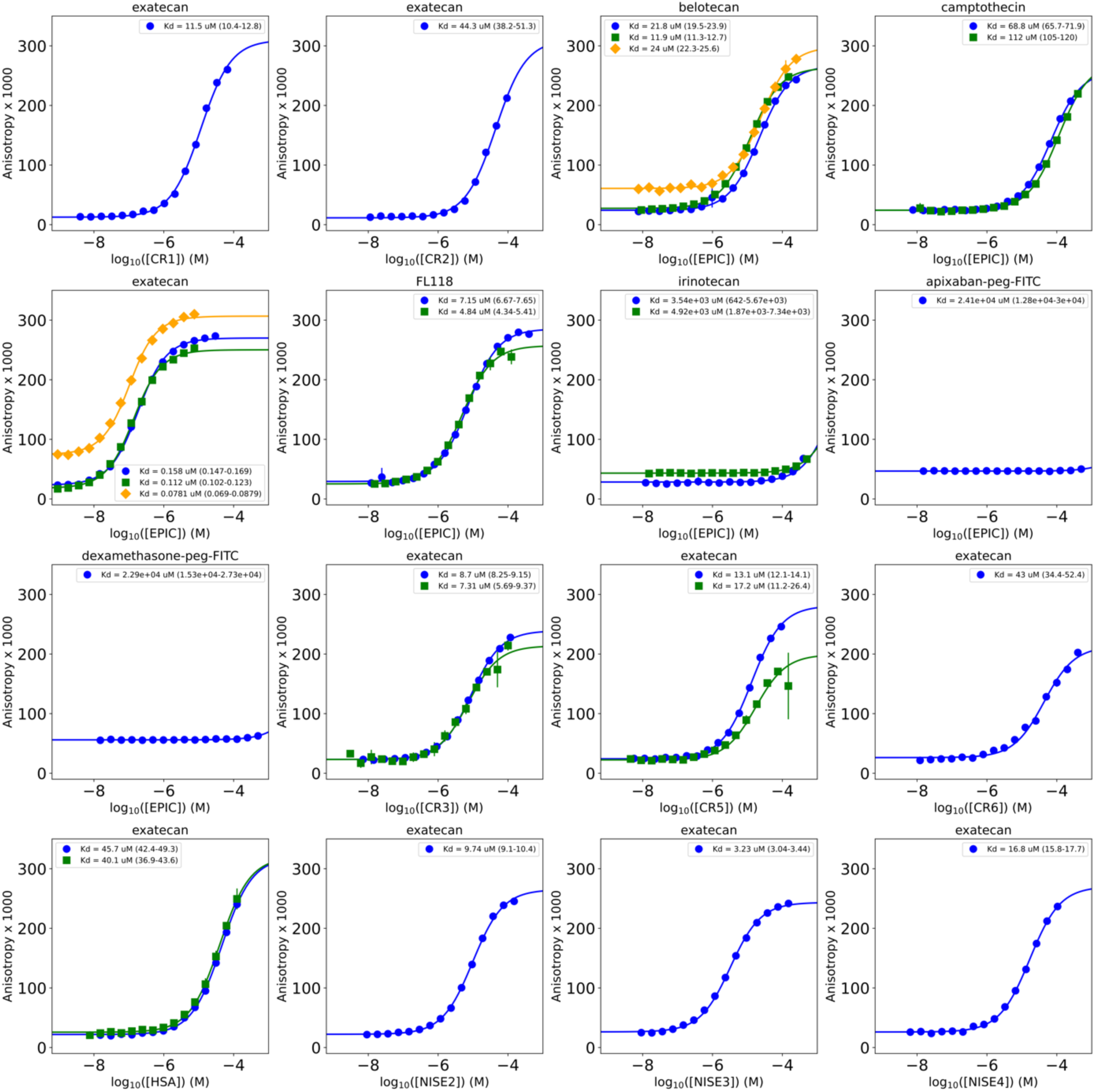
Fluorescence anisotropy data and fits to a single-site binding model. Each plot shows an individual protein (labeled on x-axis) and an individual ligand (plot title). Multiple curves are biological replicates. The figure legends show each *K*_d_ (units of µM) fit from a single-site model. The *K*_d_ shown is the mean of 1000 fits performed with bootstrapping of the optimal fit residuals. The parentheses show the lower and upper bounds of 95% confidence intervals derived from the bootstrapped fits. Error bars on the points denote standard deviations from technical replicates. Each experiment was performed with 3 technical replicates. Any systematic baseline shifts in anisotropy values between experiments are a result of slightly different gain settings on the plate reader. See Table S3 to match protein names of designed proteins with sequences. HSA is human serum albumin. See Table S2 to match ligand names to chemical structures. Buffer conditions: PBS, pH 7.4, 0.1% w/v PEG-8000.

**Fig. E12.**
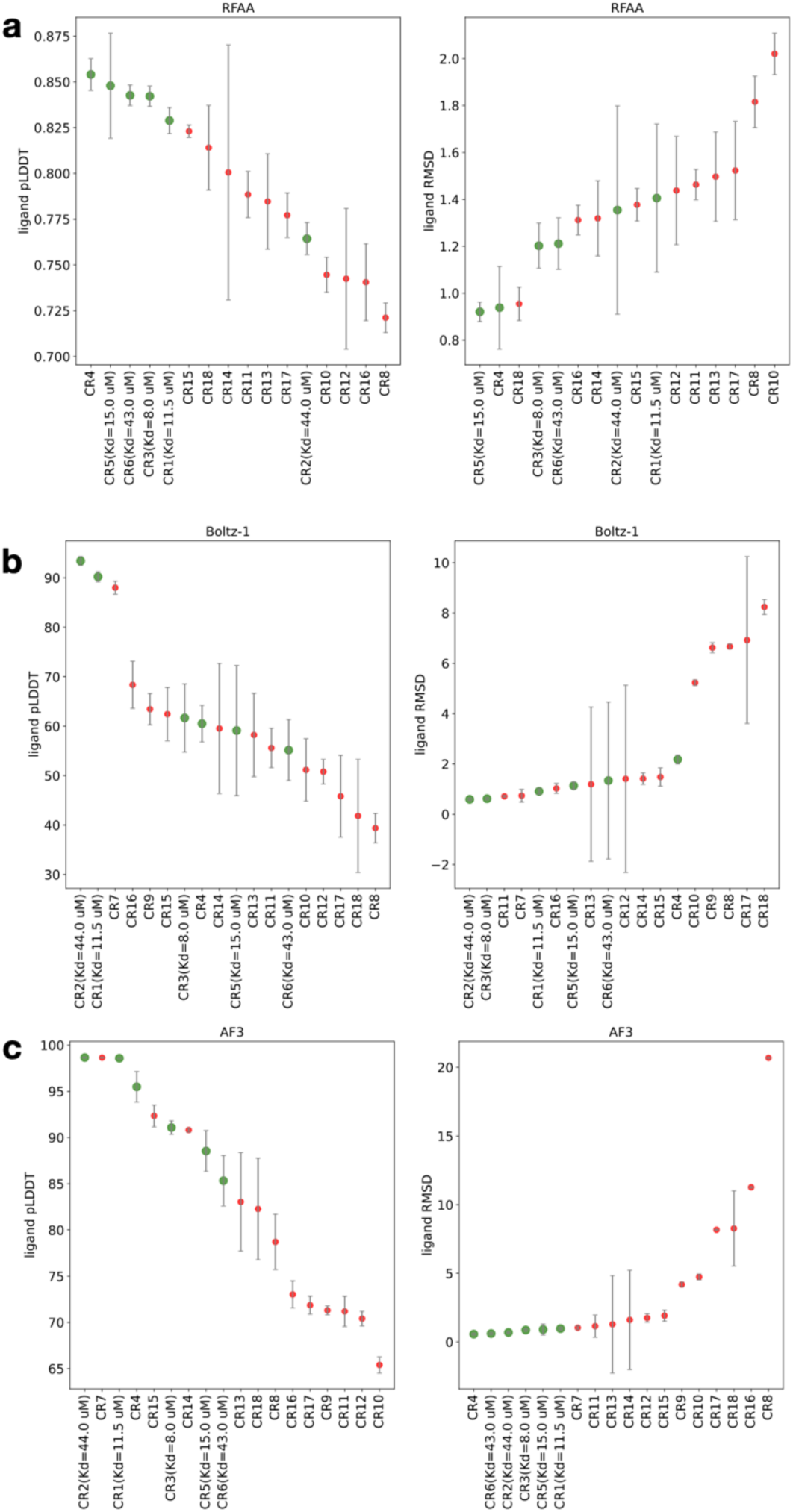
Ligand confidence and self-consistency are correlated with experimental outcomes of binding. Designs are sorted by mean ligand pLDDT (left) and ligand heavy-atom RMSD (right, RMSD with respect to the COMBS/Rosetta model after aligning on binding-site residues by C⍺ atoms) using output from **a**, RoseTTAFold-All Atom (RFAA), **b**, Boltz-1, and **c**, AlphaFold3 (AF3). We used sequences computed from COMBS and Rosetta that we experimentally tested for binding to exatecan. The designs are labeled on the x-axis (along with any measured dissociation constants). Markers for designs that bound exatecan are colored green and non-binders are colored red. Error bars denote the standard deviation of results computed across 5 different seeds. RFAA shows a clear correlation between ligand pLDDT and binding and a slightly weaker correlation for ligand RMSD. Boltz-1 shows the weakest correlations but both variables show some enrichment for binders. AF3 results correlate strongly with binding, with ligand RMSD particularly successful at ranking binders over non-binders.

**Fig. E13.**
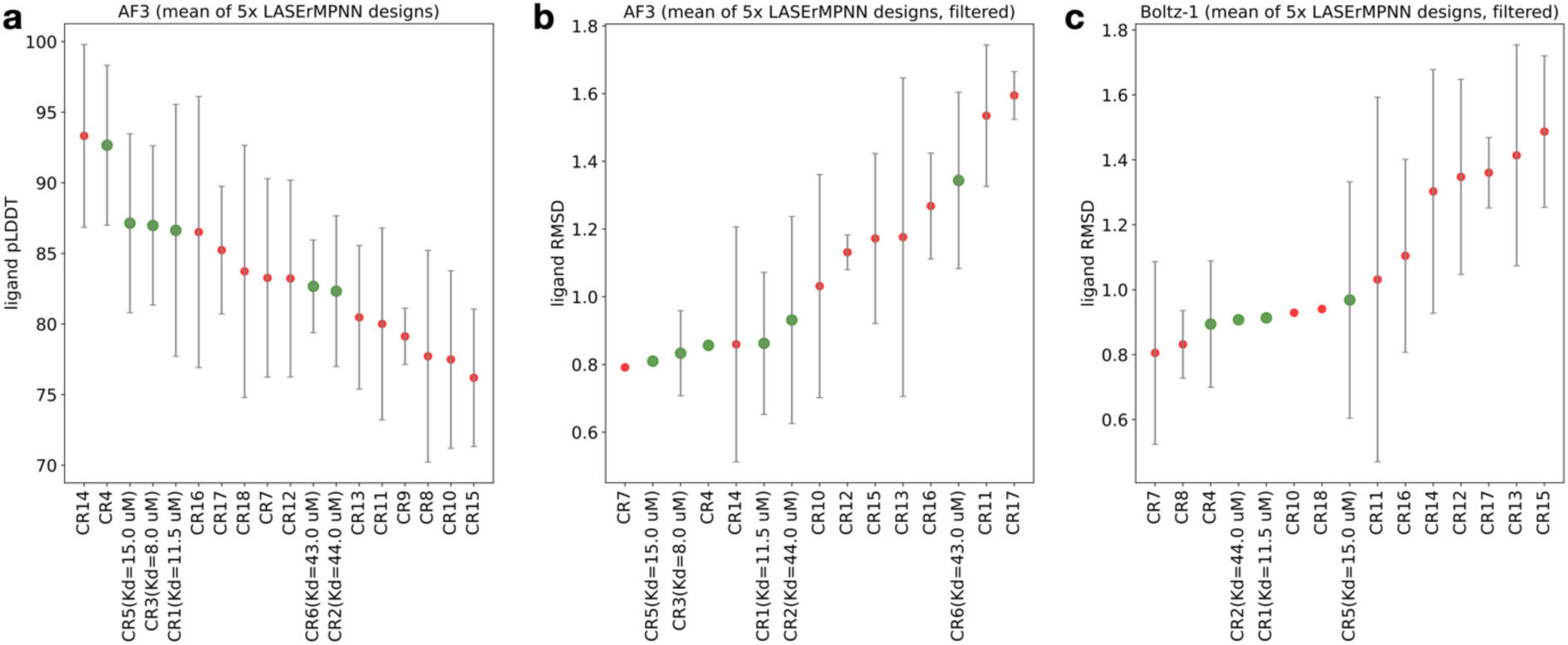
LASErMPNN and structure prediction can be used to rank poses that will lead to binders. Designs from LASErMPNN are sorted by mean ligand pLDDT (left) and ligand heavy-atom RMSD (right, RMSD with respect to the COMBS/Rosetta model after aligning on binding-site residues by C⍺ atoms) using output from **a**, RoseTTAFold-All Atom (RFAA), **b**, Boltz-1, and **c**, AlphaFold3 (AF3). These results are means and standard deviations from 5 LASErMPNN designs using the COMBS/Rosetta backbones and ligand positions (poses) as input. We discarded the COMBS/Rosetta sequences and used LASErMPNN to design 5 sequences each, which were then folded using the three protein-ligand co-structure prediction models. The designs are labeled on the x-axis (along with any measured dissociation constants). Markers for designs that bound exatecan are colored green and non-binders are colored red. **a**, The mean of ligand pLDDT across structures predicted by AF3 for 5 designed LASErMPNN sequences correlates with binding. **b**, **c**, Mean ligand RMSD computed from AF3 and Boltz-1 also enriches for binders, after removing any predicted structures with low self-consistency (C⍺ RMSD or ligand RMSD > 2 Å). These metrics can be used to prioritize poses for input into the more computationally intensive NISE algorithm for optimization.

**Fig. E14.**
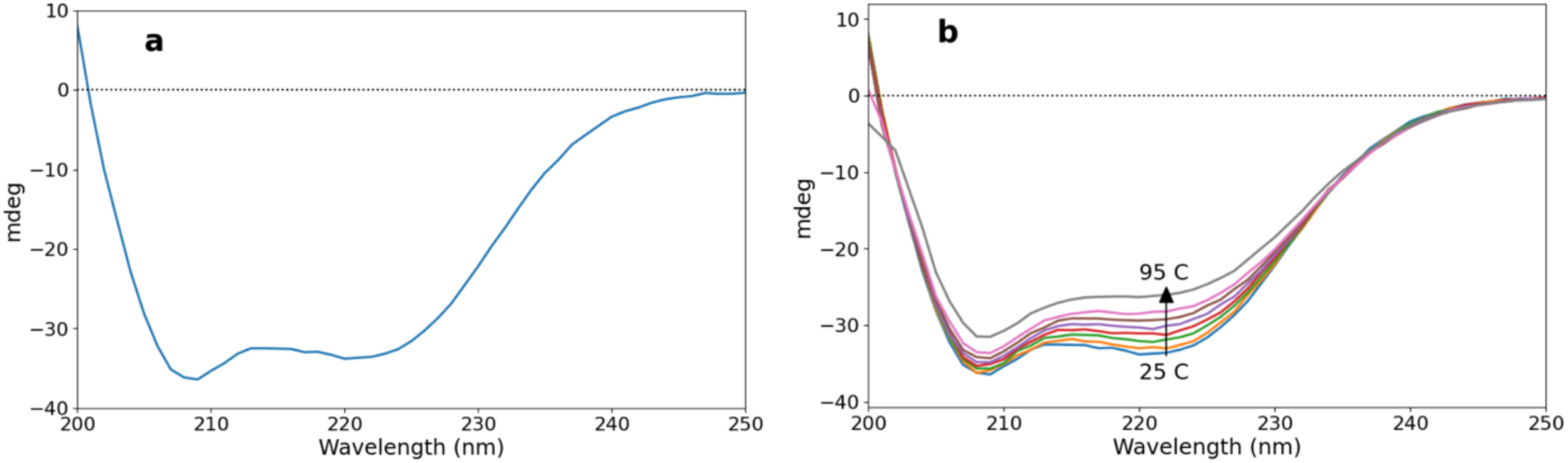
Determination of secondary structure content and thermal stability of EPIC by circular dichroism (CD) spectroscopy. CD spectra of ligand-free EPIC collected at **a,** 25 °C or **b,** varying temperatures ranging from 25 to 95 °C. No melting transition was observed. Conditions: PBS buffer, pH 7.4, protein concentration of 0.1 mg/mL (5.93 µM).

**Fig. E15.**
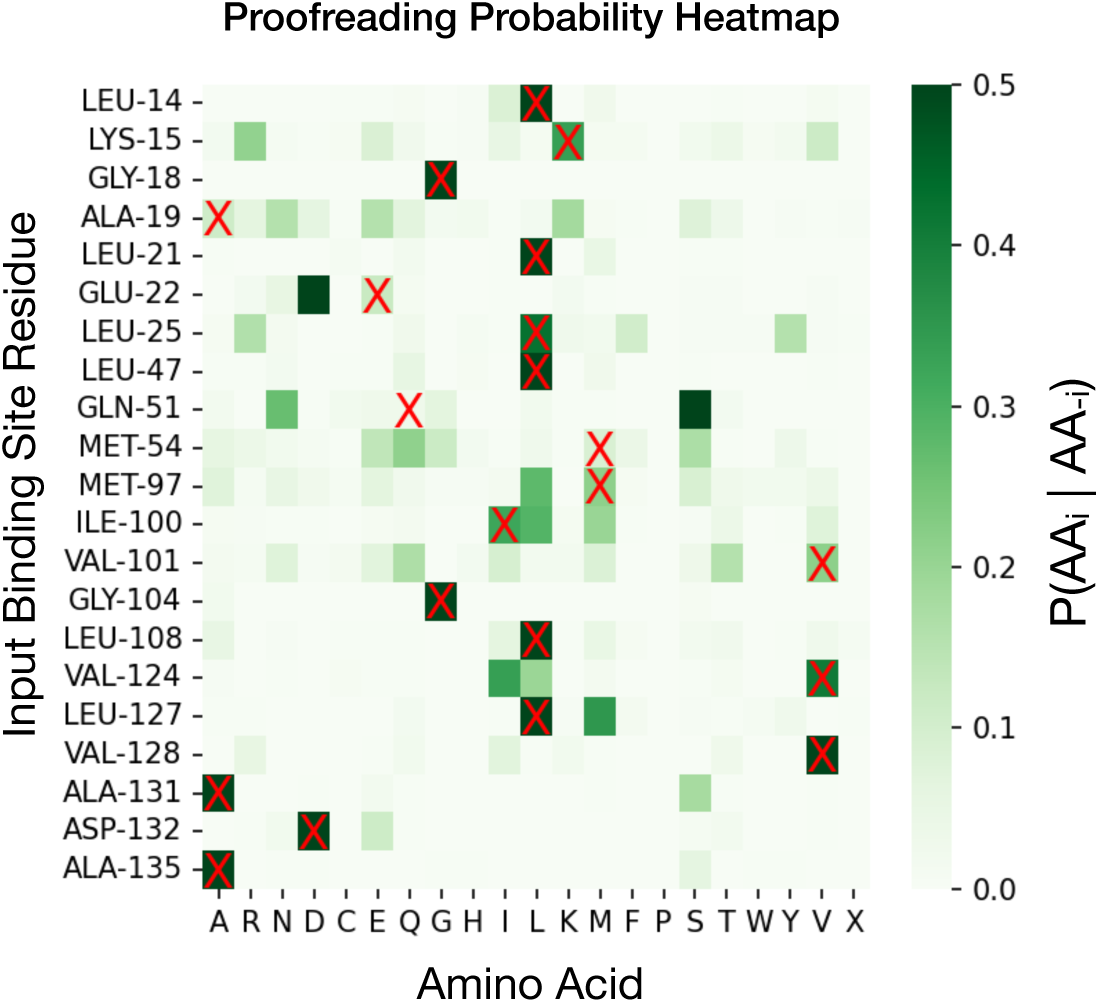
Heatmap of probabilities after proofreading of binding-site residues using the Amber-minimized RFAA prediction of exatecan-bound EPIC as the input structure to LASErMPNN. The residues in the original EPIC sequence are denoted with a red “X”, and all residues with heavy atoms within 5.0 Å of ligand heavy atoms are shown. Proofreading identifies GLN-51 to N mutation and MET-97 to L mutation, as these mutants have higher probabilities than the originally designed residue. Color bar denotes probability of the amino acid at that position conditioned on the structure and identities of all other residues in the EPIC sequence, P(AA_i_ | AA_-i_).

**Fig. E16.**
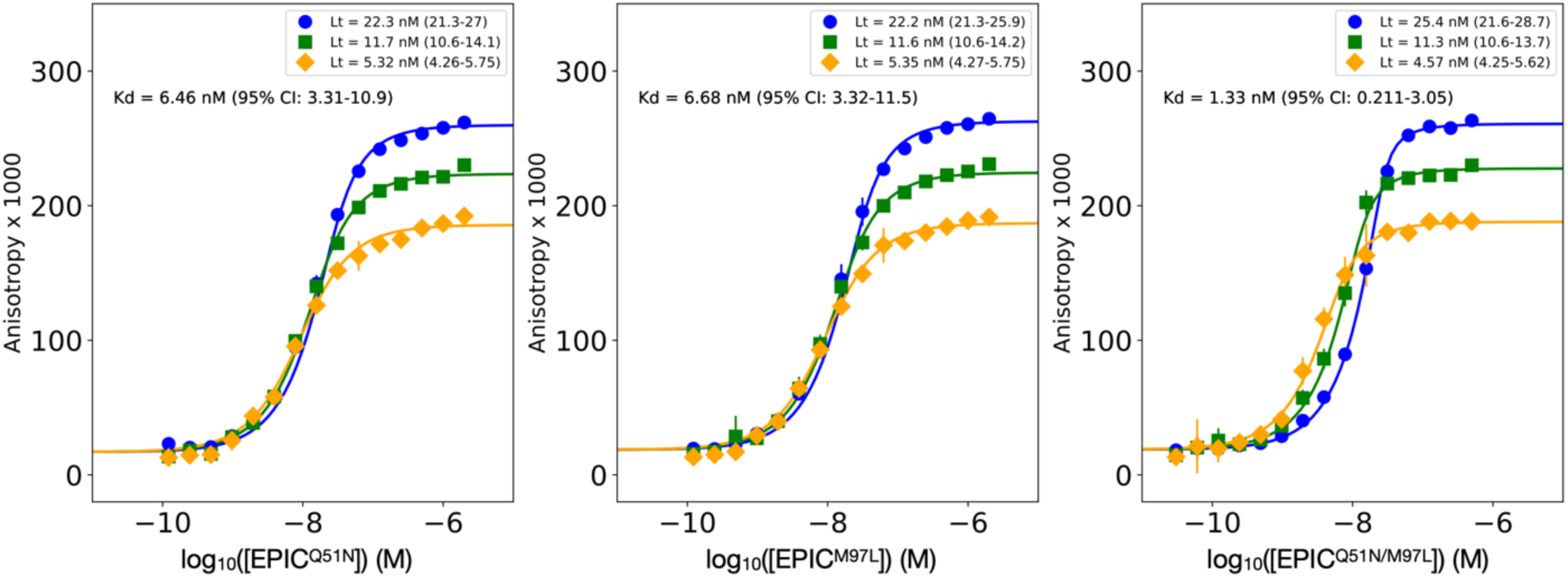
Global binding fits of EPIC mutants to exatecan. A single-site binding model with shared dissociation constant (*K*_d_) was fit globally across three fluorescence anisotropy experiments with varying concentrations of exatecan ligand for each mutant protein (labeled on x-axis). Parameters fit locally to each trace were anisotropy maximum and total ligand concentration (Lt). The anisotropy baseline was fit globally per mutant. The per-mutant globally fit *K*_d_ is shown in the plot. *K*_d_ (units of nM) is the mean from 1000 bootstrapped fits, shown with 95% confidence intervals in parentheses. The fits of total ligand concentration are shown in the figure legend with 95% confidence intervals in parentheses (derived from 1000 bootstrapped fits). Lt was allowed to vary by 15% of the known ligand concentration to account for experimental errors (shown in insets). Note that the anisotropy maxima differ across ligand concentrations because the anisotropy data are not baseline corrected, which should not affect the fit *K*_d_ values. Buffer conditions: PBS, pH 7.4, 0.1% w/v PEG-8000.

**Fig. E17.**
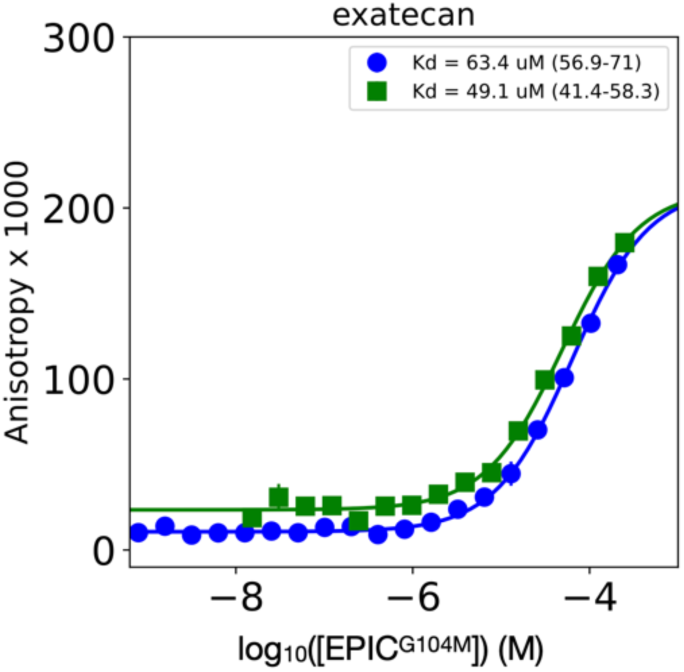
Fluorescence anisotropy data and fit to a single-site binding model for EPIC^G104M^ with exatecan. The *K*_d_ (units of µM) shown is the mean of 1000 fits performed with bootstrapping of the optimal fit residuals. The parentheses show the lower and upper bounds of 95% confidence intervals derived from the bootstrapped fits. The experiment was performed with 3 technical replicates. Buffer conditions: PBS, pH 7.4, 0.1% w/v PEG-8000.

**Fig. E18.**
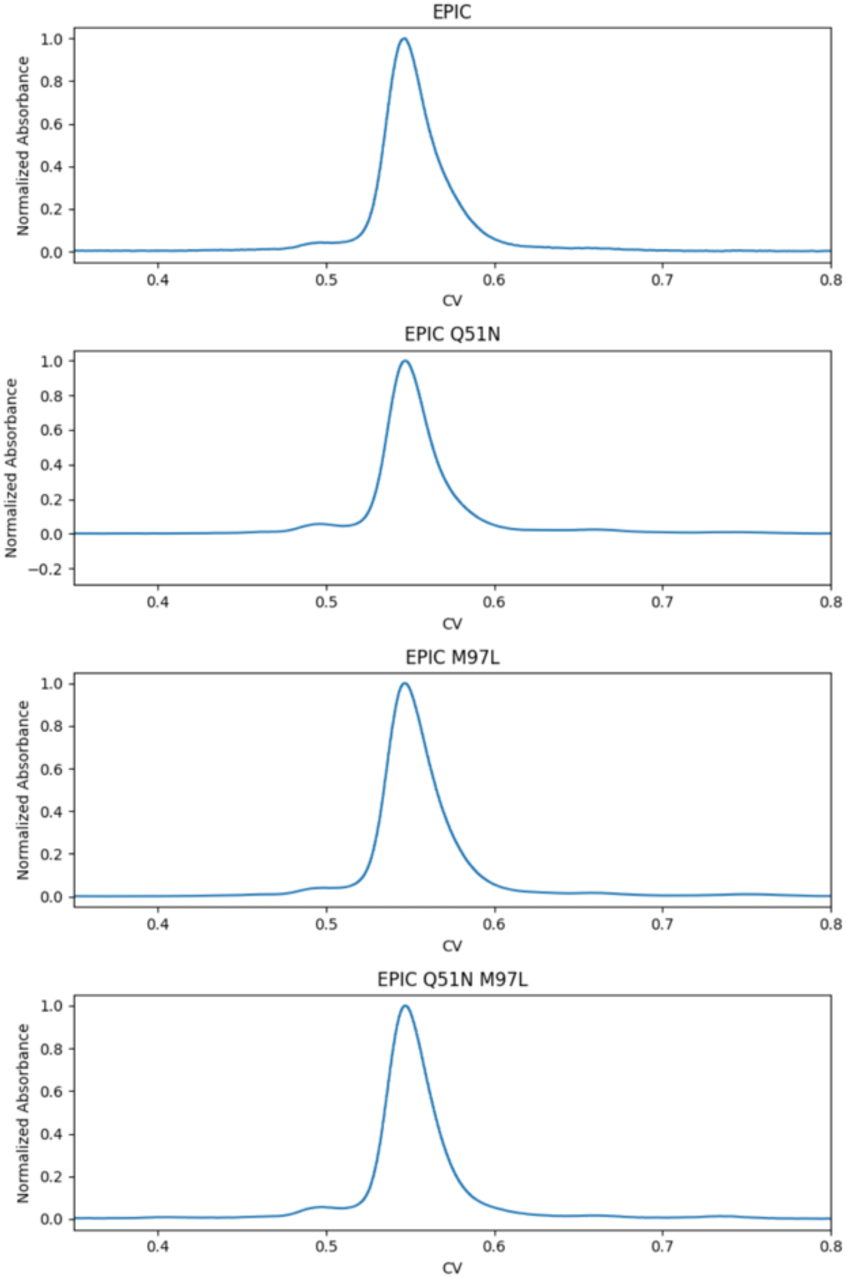
Size exclusion chromatograms of EPIC and its mutants show that each is monomeric. Ligand-free protein samples were injected at a concentration of > 100 µM and run on a Biorad 650 ENrich chromatography column in PBS buffer (pH 7.4). The proteins run slightly larger than the expected molecular weight (∼17 kDa), but the results are consistent with monomers.

**Fig. E19.**
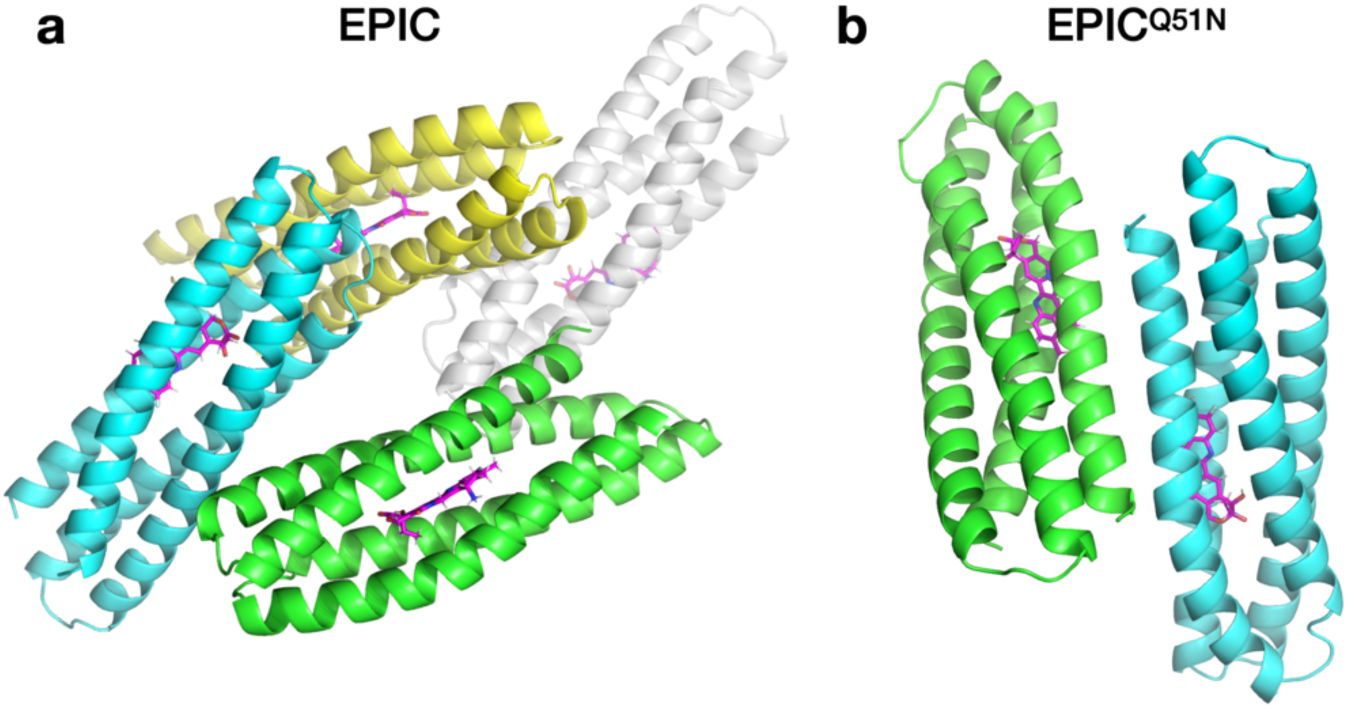
Asymmetric units of x-ray crystals of exatecan-bound EPIC and EPIC^Q51N^. **a**, EPIC was solved with four chains in the asymmetric unit, and **b**, EPIC^Q51N^ with two chains. The crystals were formed under different crystallization conditions (see supplementary methods), and the proteins crystalized in different space groups. Exatecan is colored magenta and is bound in each chain with 100% occupancy.

**Fig. E20.**
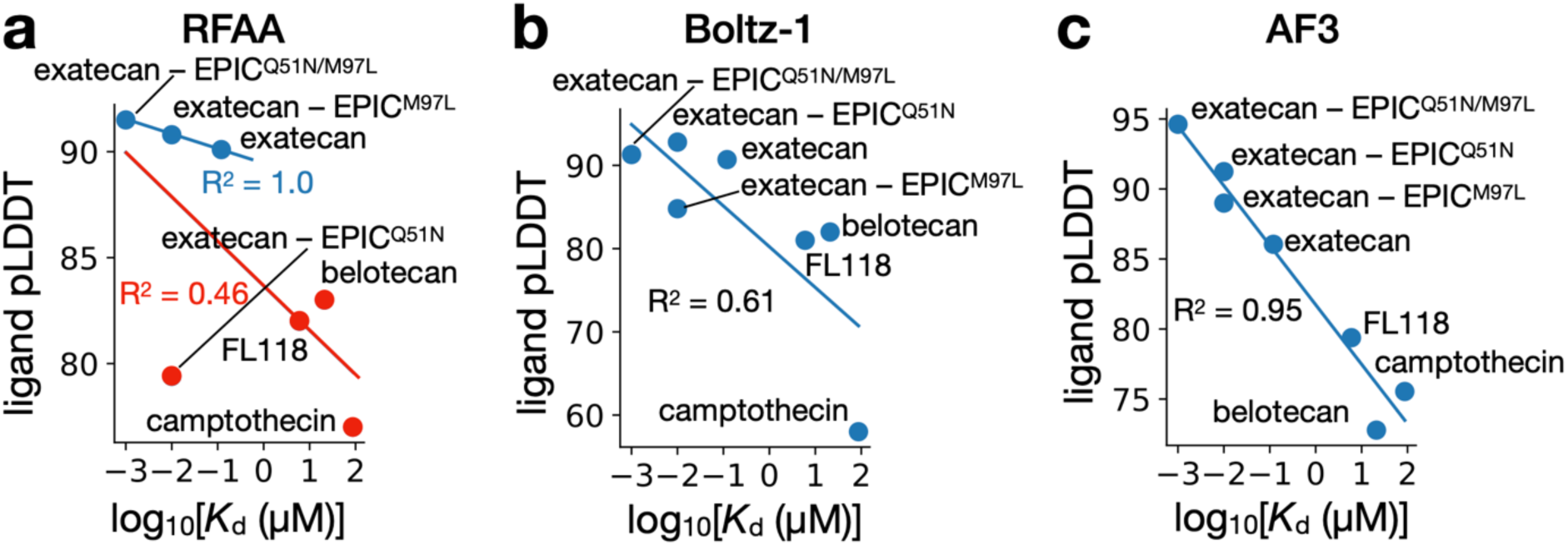
Correlation between predicted ligand confidence and binding affinity. Protein–ligand co-structure prediction for various camptothecin-class compounds with EPIC and its mutants, using **a**, RFAA, **b**, Boltz-1, and **c**, AF3. RFAA was run with 100 recycles. Points labeled with a ligand name and without a protein name are co-folded with EPIC protein sequence. For RFAA, points in red were predicted to bind in a different binding mode than exatecan in the crystal structures of EPIC and EPIC^Q51N^. For Boltz-1 and AF3, all ligands were predicted to bind in the same orientation as exatecan (consistent with crystal structures of exatecan-bound EPIC and EPIC^Q51N^). If only including the three points predicted to bind in the observed crystallographic binding mode (blue points), the correlation between affinity and ligand pLDDT is strong (R^2^ = 1) but statistically underpowered. The correlation drops dramatically if including the red points (R^2^ = 0.46). Boltz-1 shows a stronger correlation and AF3 shows the strongest correlation with affinity. Data for non-binders (not shown): EPIC and dexamethasone, ligand pLDDT (70.4, 35.9, 72.7 for RFAA, Boltz-1, and AF3, respectively); EPIC and apixaban, ligand pLDDT (83.4, 52.5, 74.8 for RFAA, Boltz-1, and AF3, respectively).

**Fig. E21.**
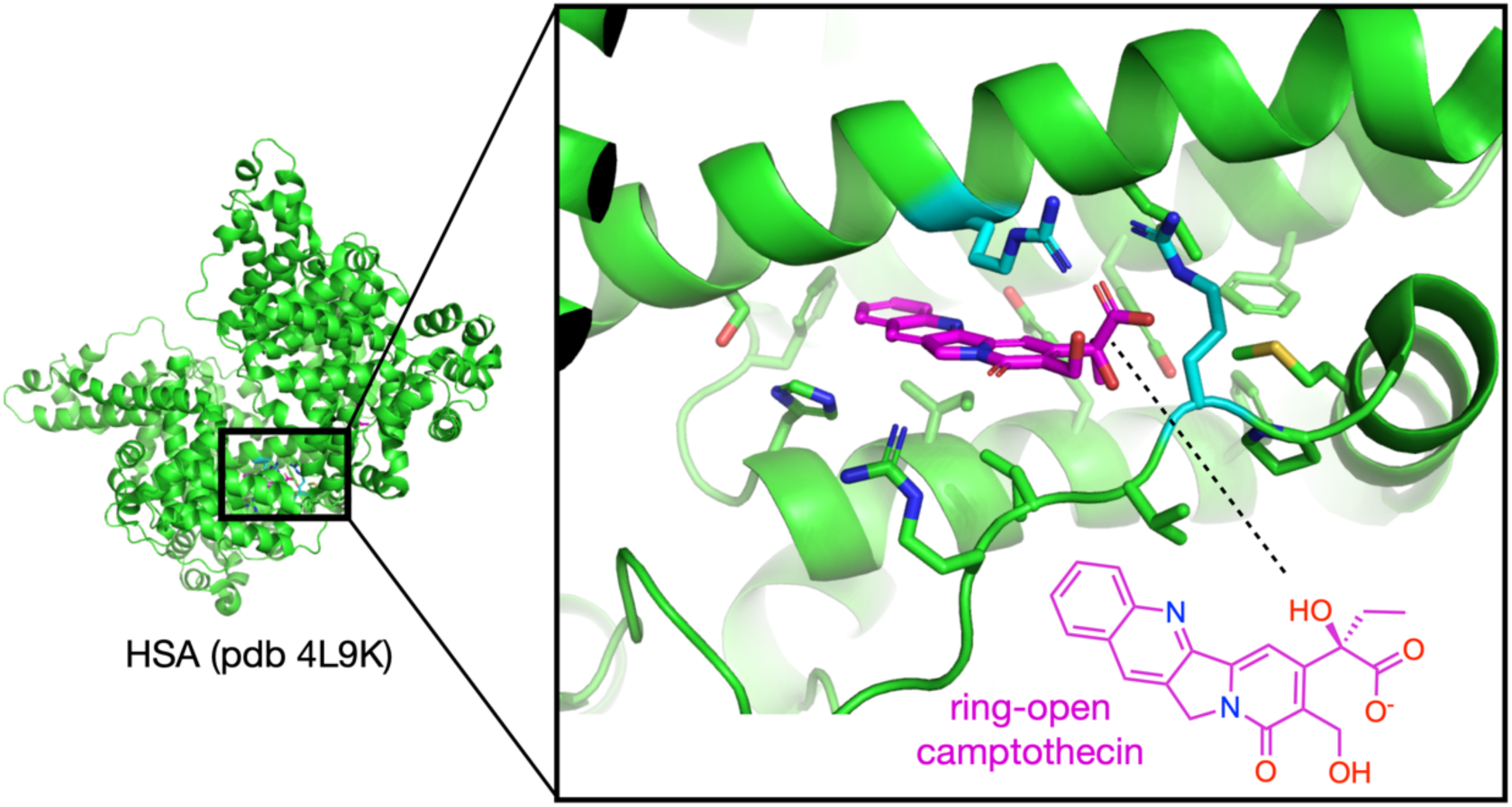
Camptothecin-bound structure of human serum albumin. Camptothecin (magenta) is bound to human serum albumin (HSA) in its ring-open carboxylate form, stabilized by nearby positively charged Arg residues (cyan). PDB accession code: 4L9K.

**Fig. E22.**
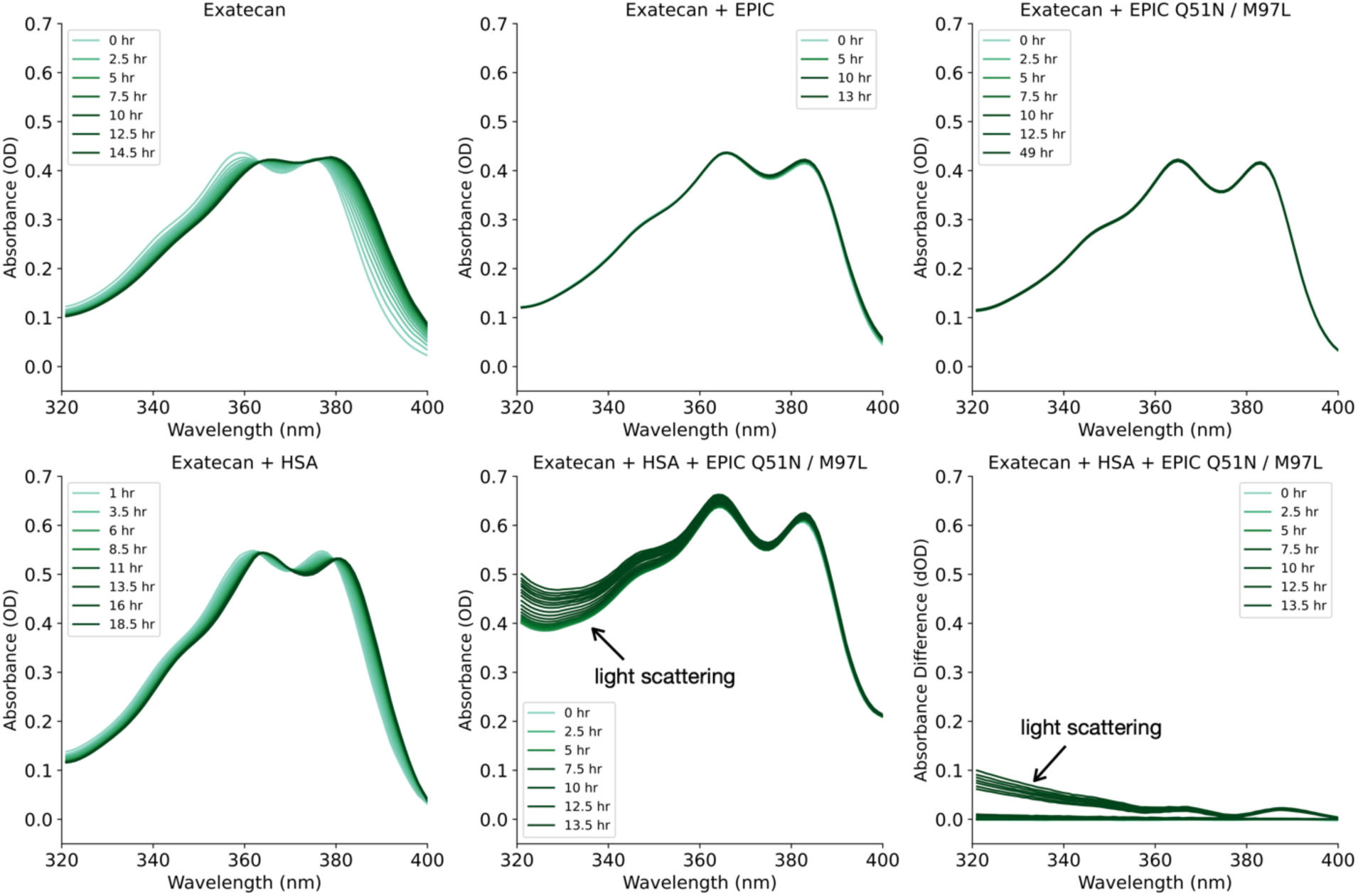
Time-resolved absorbance spectra of exatecan alone or with various proteins. Experiments were initiated from a ring-closed form of exatecan from a DMSO stock and spectrally monitored for ring opening. Experiments were performed at room temperature (20°C) in phosphate-buffered saline (pH 7.4) with 20 µM exatecan. Spectra for each 30 min time interval are shown, but only some spectra with various time delays are labeled in the figure legend to avoid cluttering. Designed proteins (EPIC and EPIC^Q51N/M97L^) were included at equimolar concentrations (20 µM), whereas human serum albumin (HSA) was included at physiological concentration (500 µM), where most exatecan is expected to be bound. The high concentration of HSA resulted in some light scattering, which increased as a function of time. We predominantly removed the scattering component of the measured spectra through baseline correction (fitting and extrapolating a polynomial to wavelengths 450-850 nm). Parts of the spectra that show residual contributions from scattering (after correction) are labeled with an arrow in the lower right plots. The spectral dynamics observed in the HSA and EPIC^Q51N/M97L^ mixture are consistent with a 100% scattering component. The lower right plot shows the absorbance difference of the data to the left (the time = 0 hr spectrum was subtracted for each subsequent spectrum).

**Fig. E23.**
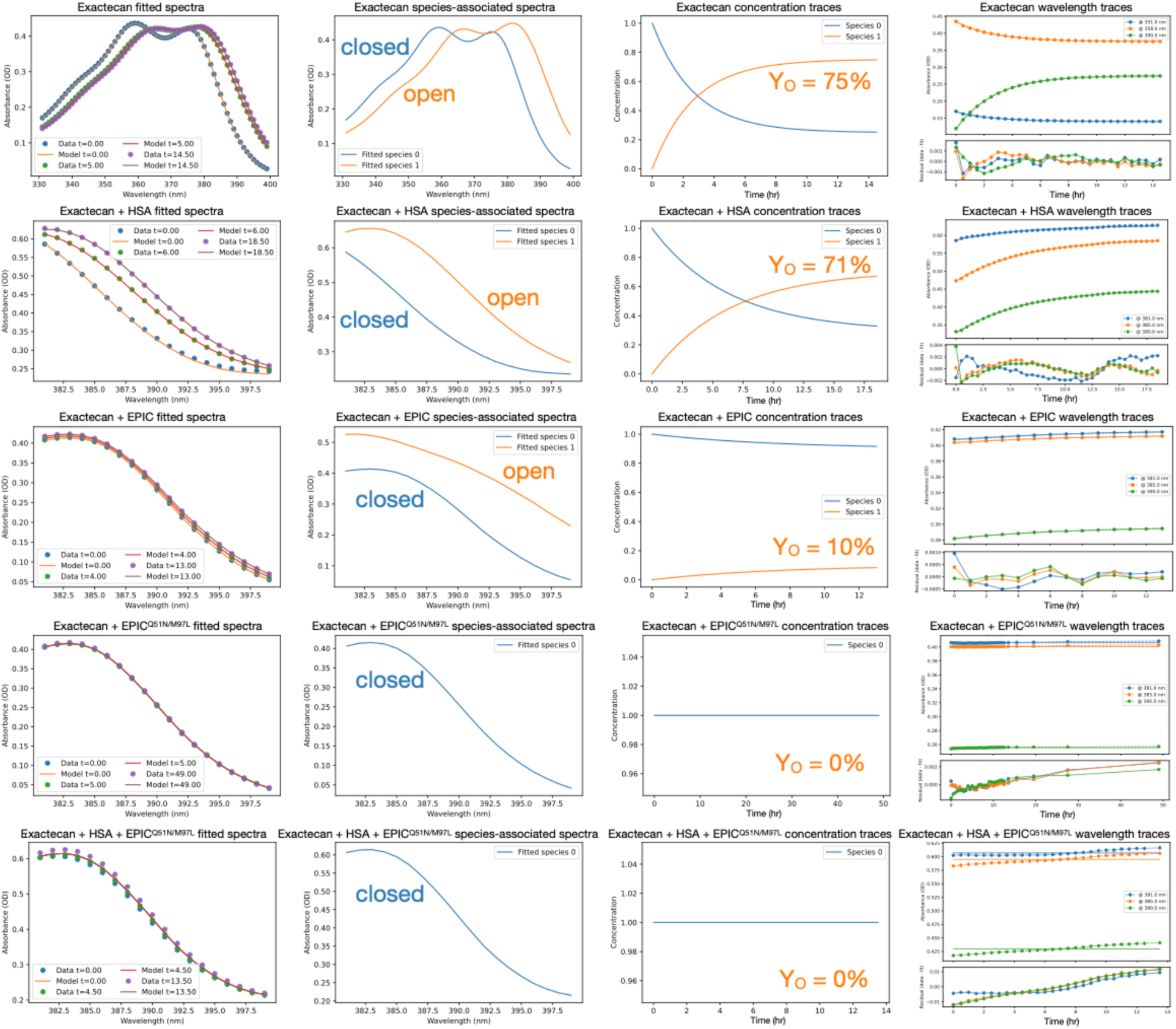
Fits of time-resolved hydrolysis experiments. Spectral ranges of exatecan absorbance spectra were fit to a one- or two-state kinetic model. The one-state model was a constant non-evolving species. The two-state model (0 or 1 in inset) was a reversible model with forward and reverse rate constants, with initial condition that all population starts in state 0. State 0 (blue) corresponds to the exatecan lactone-ring-closed state. State 1 (orange) corresponds to exatecan lactone-ring-open state. From left to right is shown i) a few spectra (circles) with overlayed fits (lines) at various time delays (units of hr), ii) the species-associated spectra resulting from the fit of the kinetic model, iii) the concentration profiles of species 0 and 1 from the fits, and iv) a few time traces at select wavelengths (markers) overlayed with the fits (lines), with fit residuals shown below (for a much narrower range of y-axis. Note the residuals are all very small relative to the magnitude of the fit data.). Experiments of exatecan that include EPIC^Q51N/M97L^ are well-approximated by a single-state (constant) model of ring-closed exatecan with no time evolution. Equilibrium yields of ring-open exatecan (Y_O_), computed from a thermodynamic model using the fit rate constants, are shown in the plots of concentration profiles. Fits were performed on data from Fig. E22. See Fig. E22 for experimental details.

**Fig. E24.**
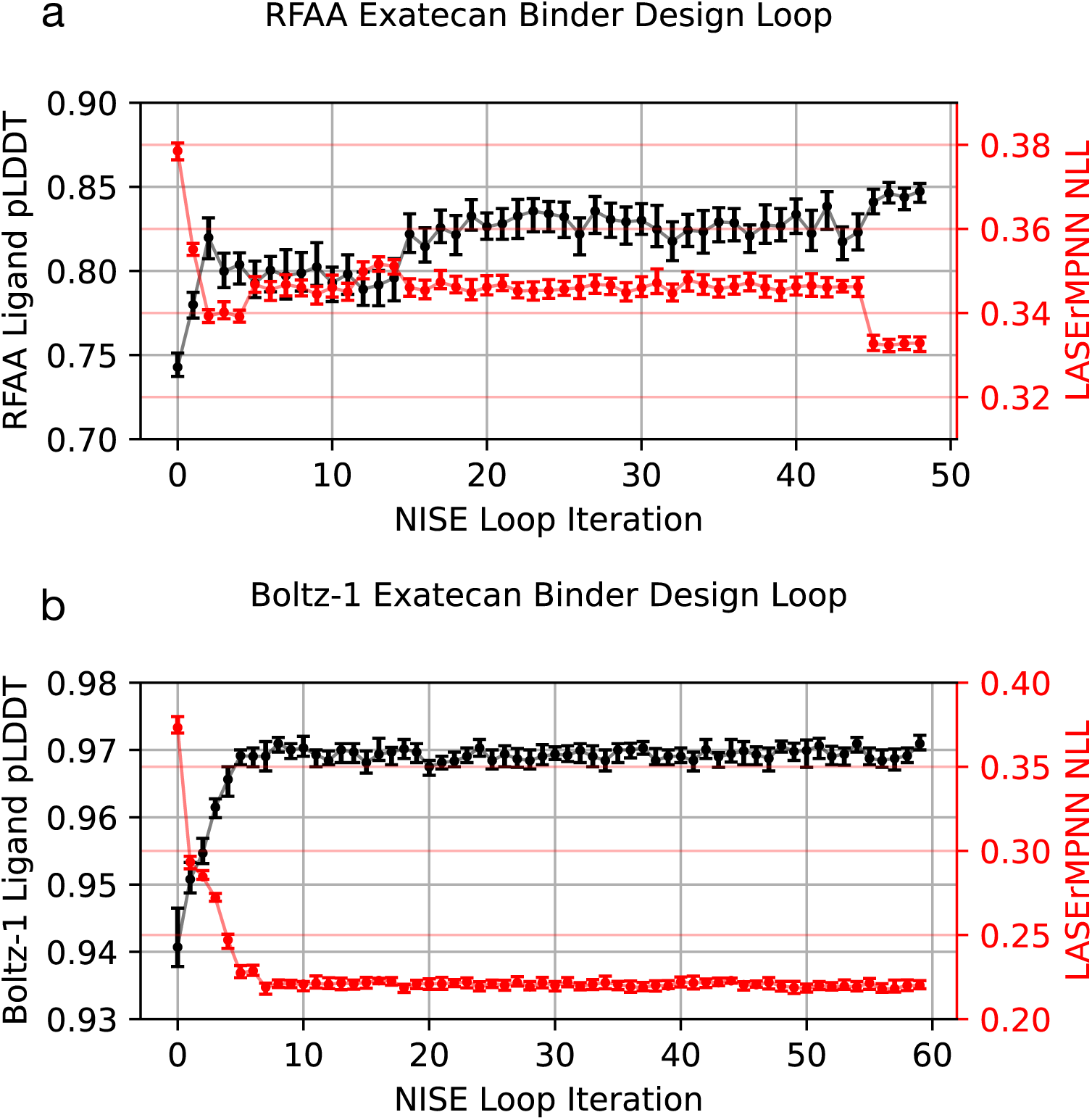
NISE jointly optimizes sequence and structure confidence using either RFAA or Boltz-1. **a**, Ligand pLDDT vs LASErMPNN Negative Log-Likelihood (NLL) for a NISE design loop mimicking that which generated EPIC using ligand pLDDT from RFAA as a proxy for structure confidence. **b,** Ligand pLDDT vs LASErMPNN NLL for a NISE design loop mimicking that which generated EPIC using Boltz-1 in place of RFAA. Boltz-1 appears to optimize joint confidence more efficiently than RFAA but both models jointly optimize the sequence and structure confidence. Data points show the value of the third quartile for sampled pLDDT and first quartile for NLL, with error bars representing 95% confidence intervals for these values computed by bootstrapping (N=100,000) the sampled data.

**Fig. E25.**
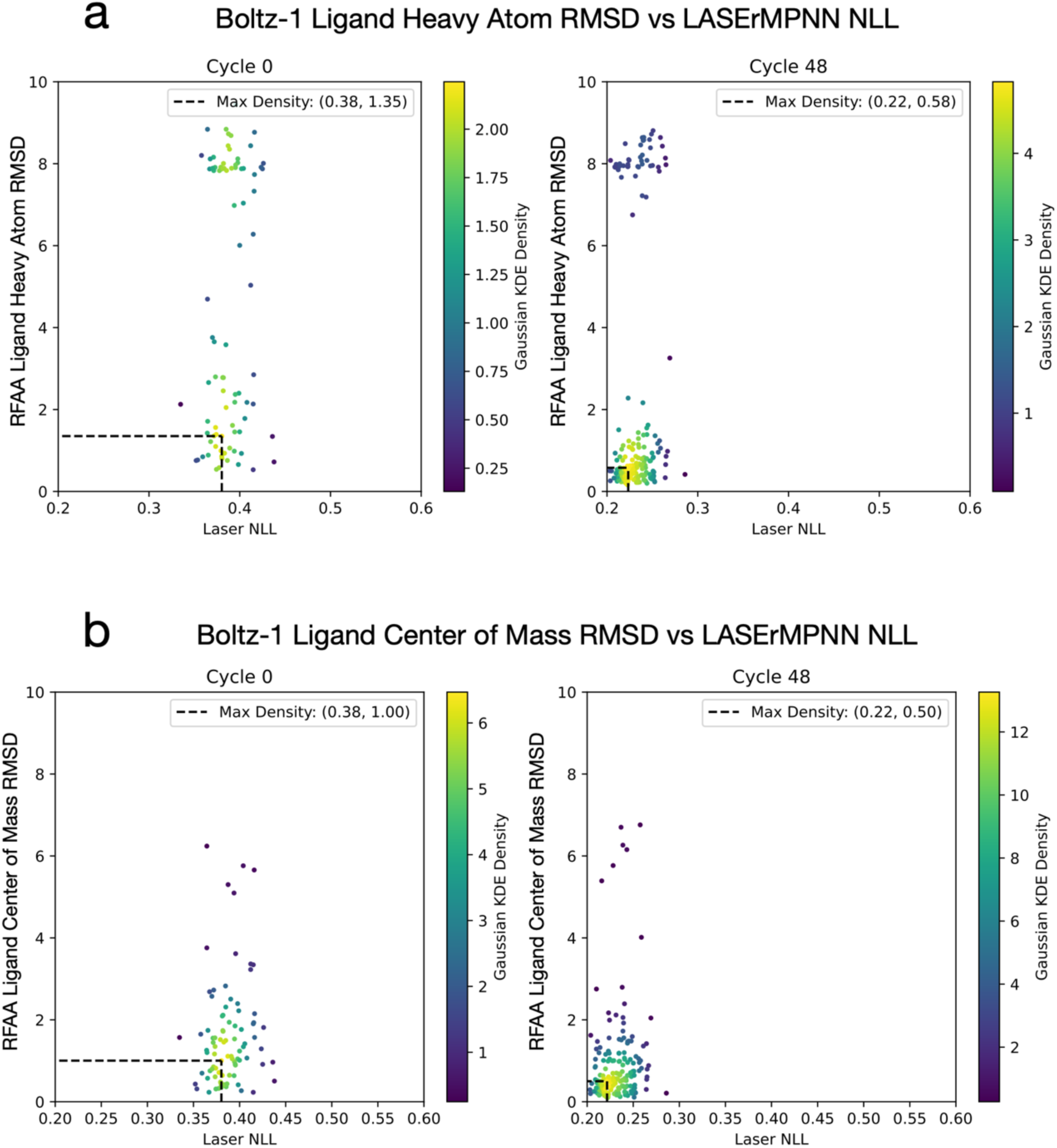
Boltz-1 outperforms RFAA at predicting exatecan bound in the designed orientation. **a**, Ligand heavy atom RMSD between input to LASErMPNN and Boltz-1 predicted structure vs LASErMPNN negative log-likelihood (NLL) for first NISE iteration (input backbone-ligand pair only) and the 48^th^ NISE iteration. **b**, Ligand center-of-mass RMSD between input to LASErMPNN and Boltz-1 predicted structure vs LASErMPNN negative log-likelihood for first NISE iteration and 48^th^ NISE iteration. Throughout the NISE design trajectory, Boltz-1 is generally able to place the ligand’s center of mass within 2.0 Å of the binding site. Compared to RFAA (see Fig. E5), Boltz-1 shows better success at predicting the designed binding mode.

**Fig. S1.**
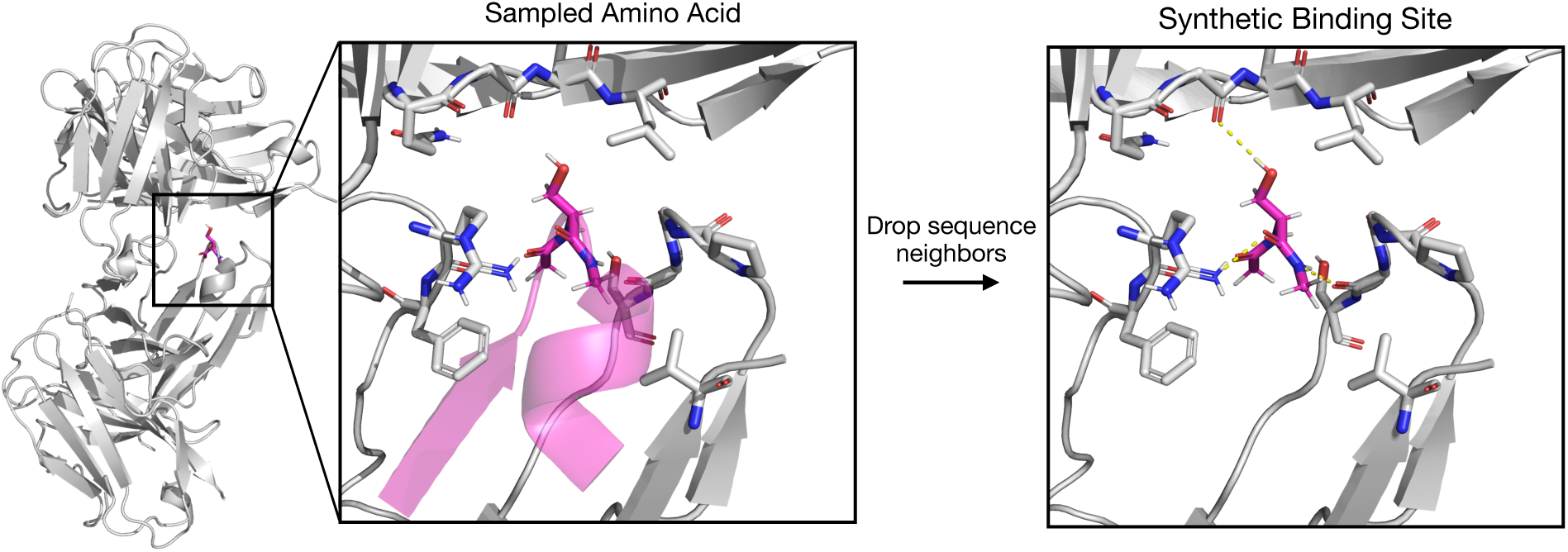
Amino acids are used as “pseudoligands” to augment ligand training data for LASErMPNN. An example pseudoligand is shown, using a serine residue. A synthetic binding site is generated around a serine pseudoligand (magenta sticks) sampled within the protein complex with PDB Code 7w7q. Methyl caps are applied to ensure the pseudoligand is physically plausible and in distribution of the pre-trained ligand encoder. Sequence-local residues, which are dropped from the protein graph to form the synthetic binding site, are depicted as a transparent magenta cartoon in the first panel. Sidechains within 5.0 Å of the synthetic ligand forming a synthetic binding site are shown as grey sticks.

**Fig. S2.**
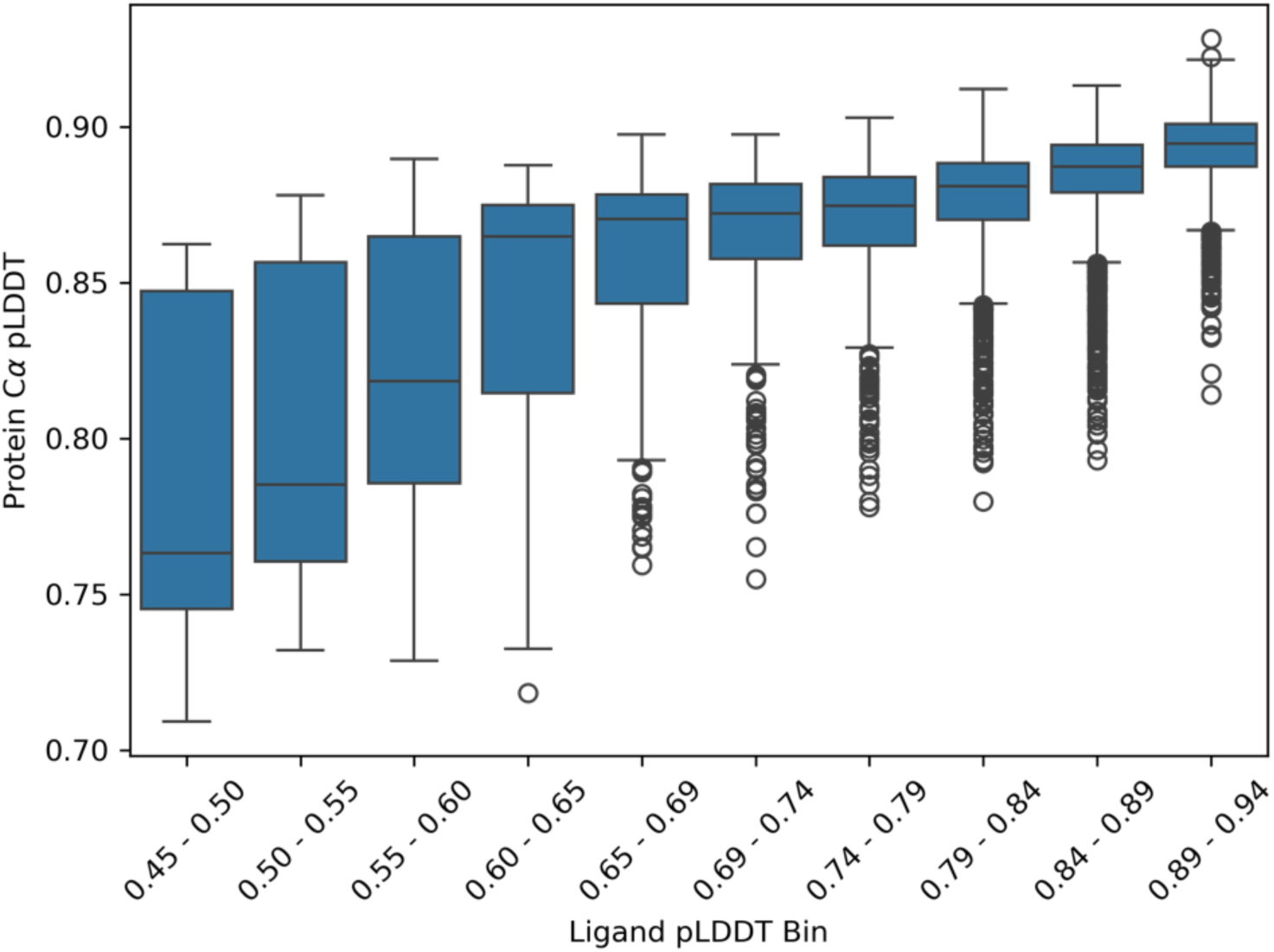
Ligand pLDDT and protein C⍺ pLDDT are positively correlated. We plot pooled ligand and protein pLDDT values generated by RoseTTAFold-All Atom (RFAA) over 48 iterations of NISE using the CR4 exatecan-bound co-structure as input. The box plots show protein C⍺ pLDDT statistics of designs with ligand pLDDT falling in the labeled bin range. The data show that by maximizing ligand pLDDT, NISE also implicitly maximizes protein C⍺ pLDDT.

**Fig. S3.**
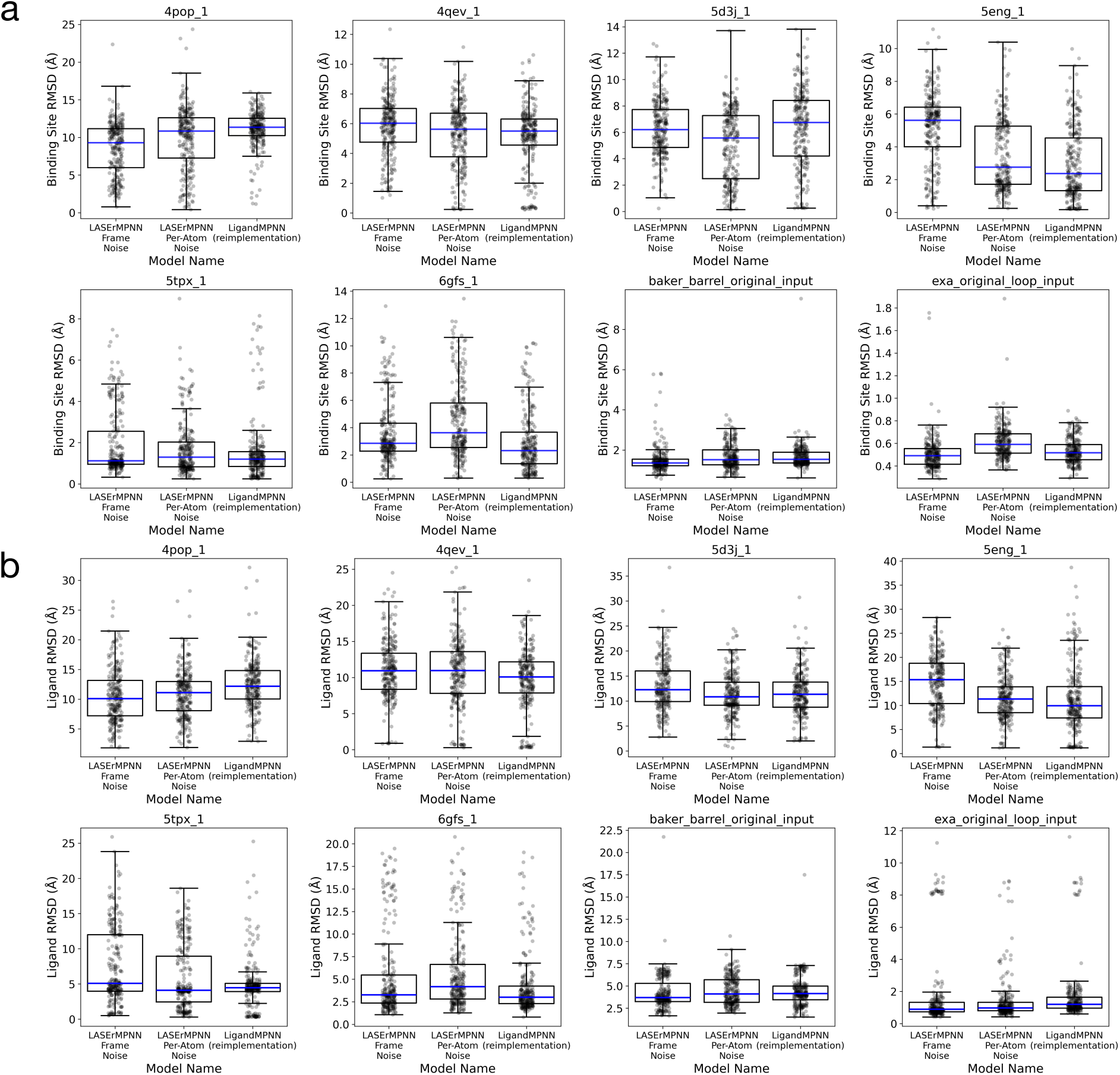
LASErMPNN and LigandMPNN models perform similarly by self-consistency RMSDs on a small test set. **a**, AlphaFold3 predicted binding site residue C⍺ RMSD distributions for LASErMPNN models trained with different protein backbone noising strategies and a reimplementation of the LigandMPNN architecture trained on the same dataset. **b**, AlphaFold3 predicted ligand RMSD distributions for LASErMPNN models trained with different protein backbone noising strategies and a reimplementation of the LigandMPNN architecture trained on the same dataset. Median values are highlighted in blue.

**Fig. S4.**
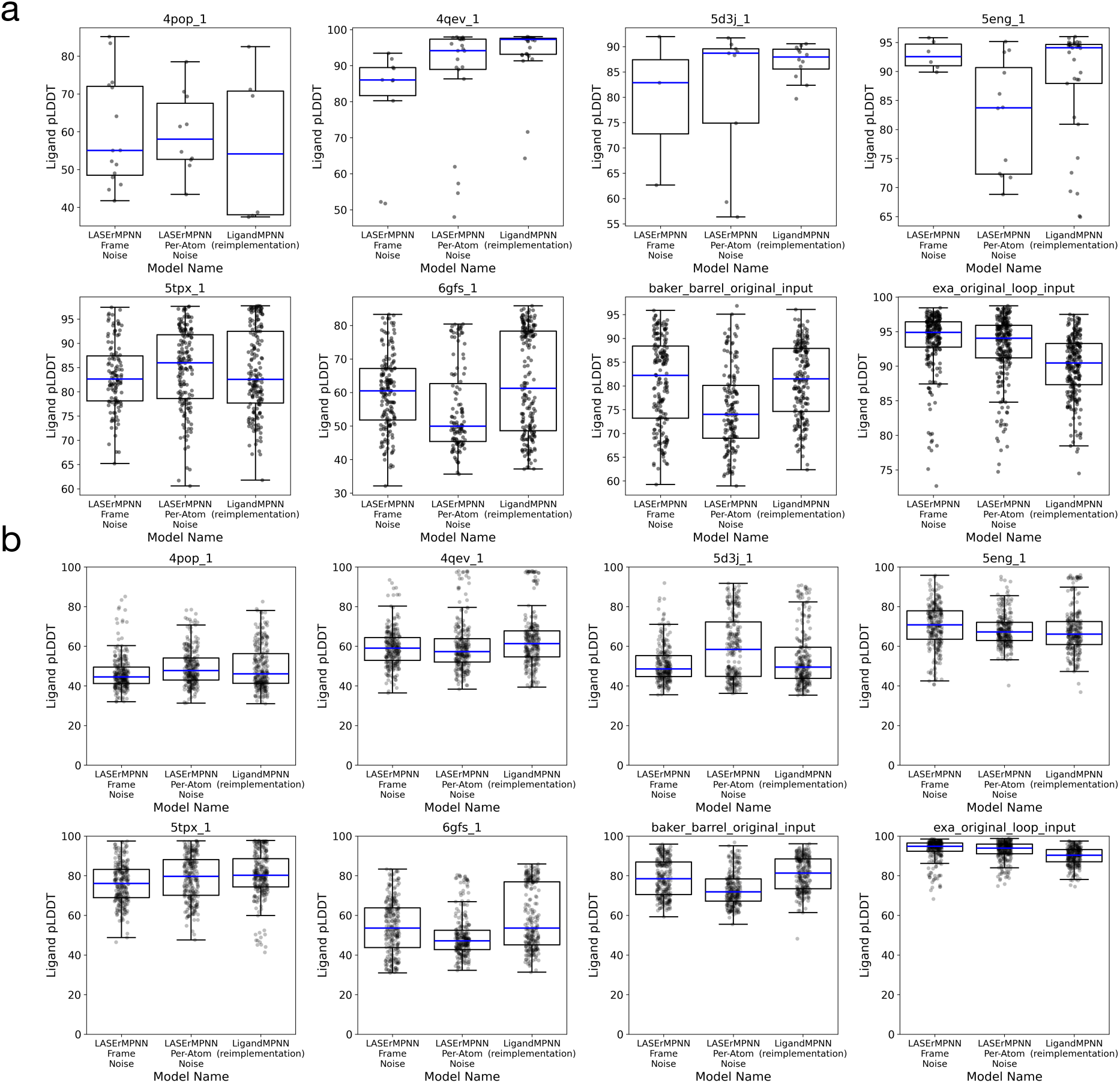
LASErMPNN and LigandMPNN models generate sequences with similar AlphaFold3 ligand pLDDT distributions on a small test set. **a**, AlphaFold3 predicted ligand pLDDT distributions only over self-consistent predictions (ligand RMSD < 5.0 Å, binding site C⍺ RMSD < 5.0 Å). **b**, AlphaFold3 predicted ligand pLDDT distributions for LASErMPNN models trained with different protein backbone noising strategies and a reimplementation of the LigandMPNN architecture trained on the same dataset. Median values are highlighted in blue.

**Fig. S5.**
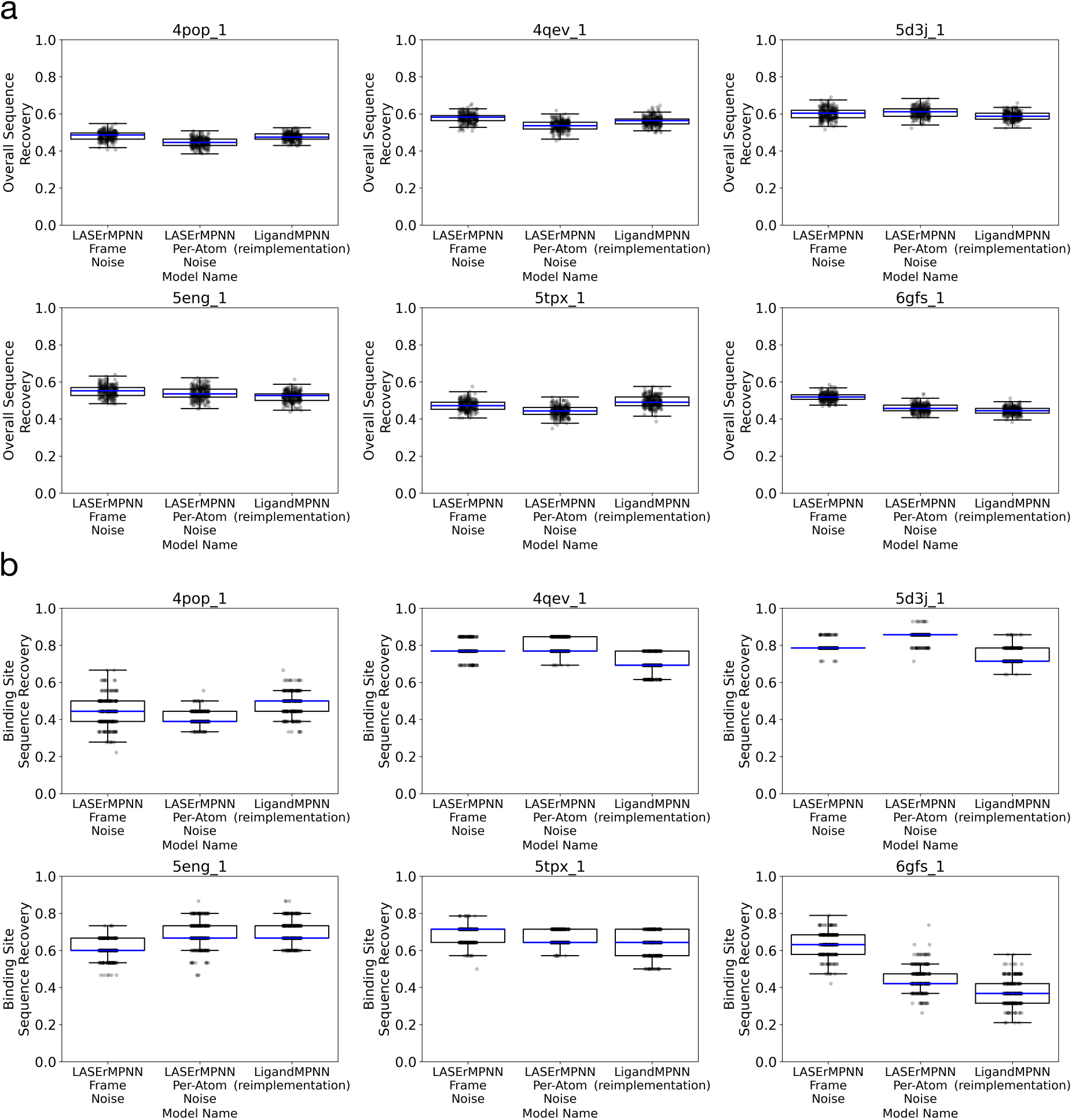
LASErMPNN and LigandMPNN models generate sequences with similar sequence recoveries over a small test set. **a**, Overall sequence recovery distributions for single chain test set of ligand binding proteins. **b**, Distributions of binding-site sequence recovery of single-chain test set of ligand-binding proteins. Median values are highlighted in blue.

**Fig. S6.**
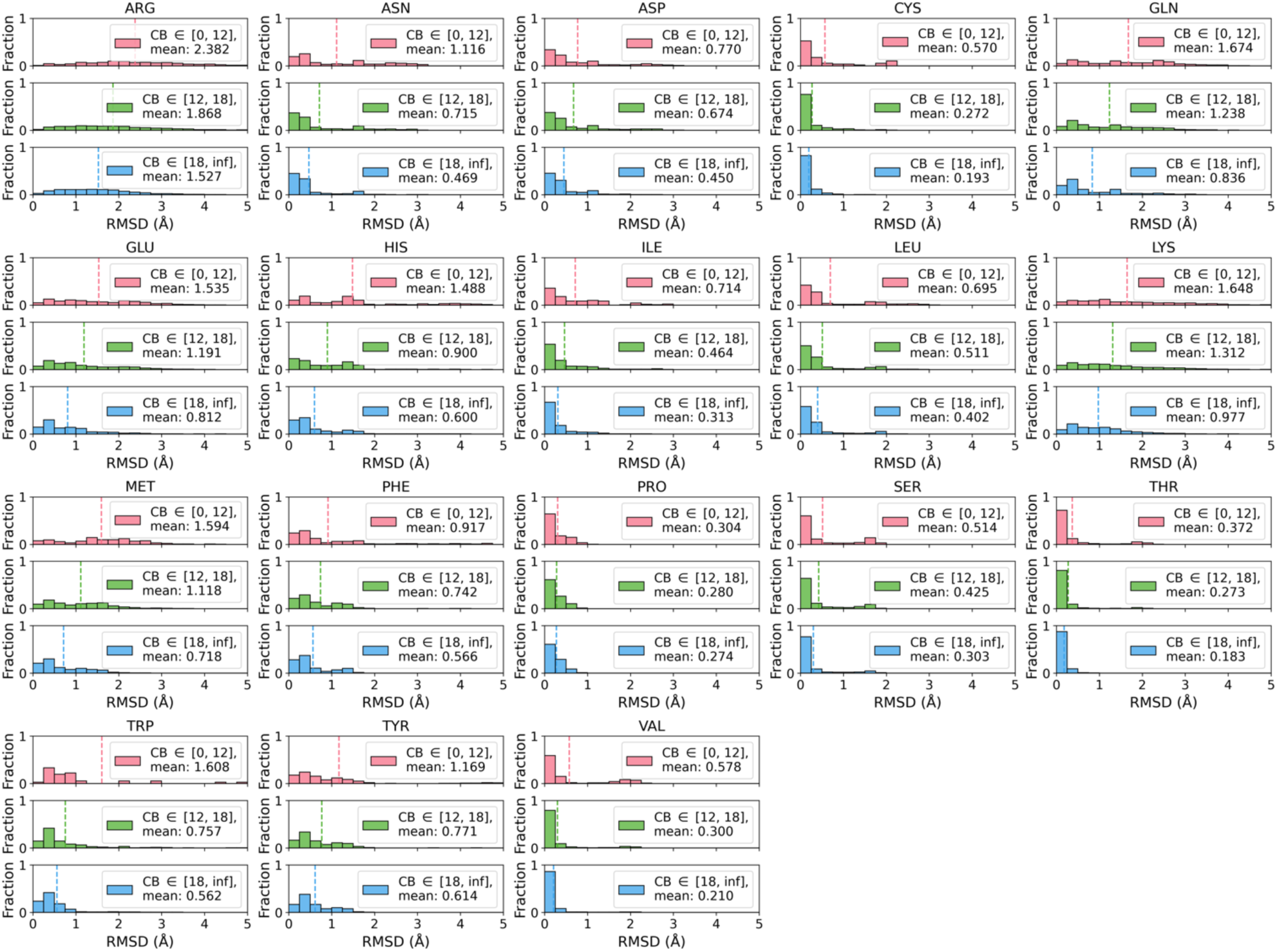
LASErMPNN generates sidechain conformations with low RMSD to crystallographic conformations. LASErMPNN repacked sidechain RMSD distributions as a function of neighboring Cβ atoms (used as a proxy for residue burial) computed over the test set of the LigandMPNN data split (n=317). For all residues, sidechain RMSD tends to decrease as the sidechain becomes more buried.

**Fig. S7.**
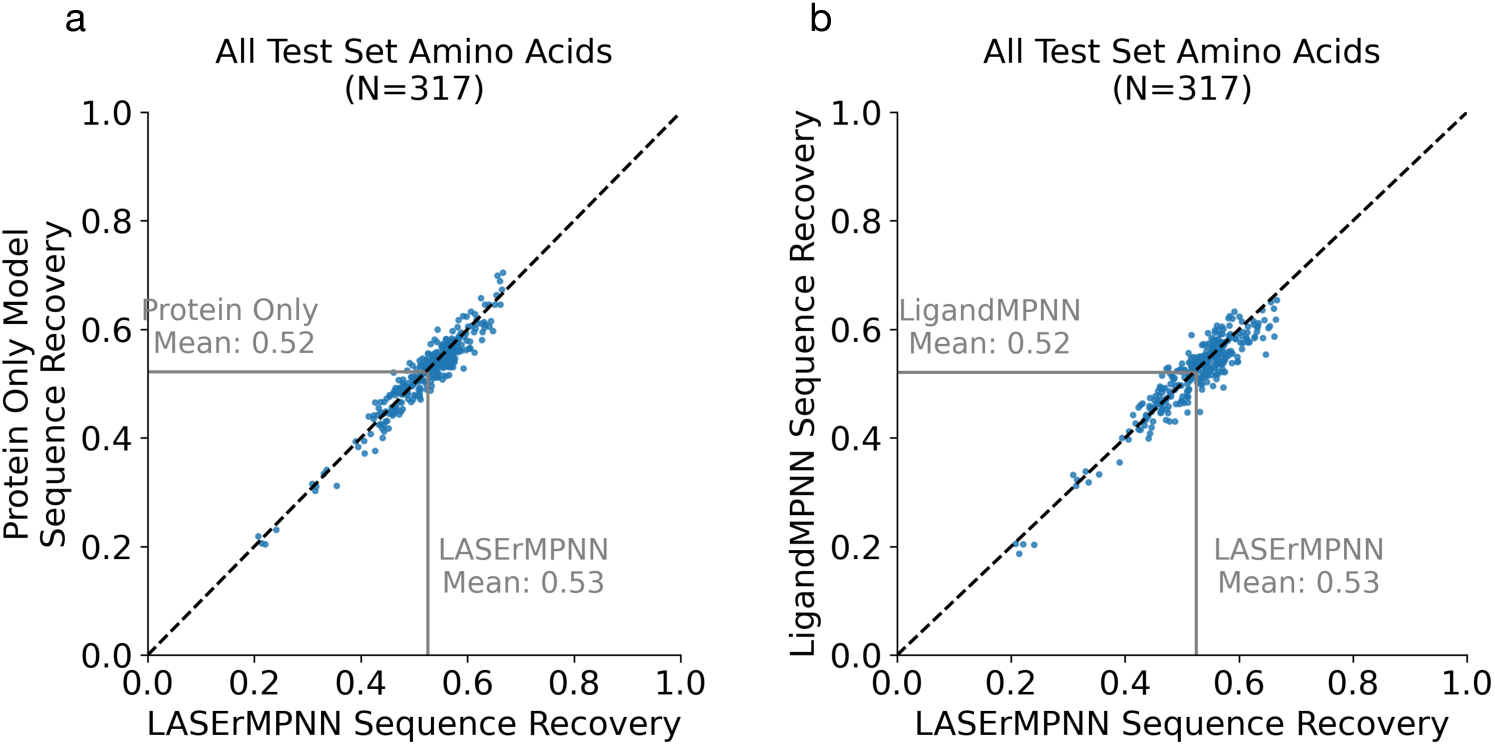
Overall residue sequence recovery is similar between models. The “protein only model” refers to a LASErMPNN model trained without ligand information and LigandMPNN refers to a reimplementation of the LigandMPNN architecture. All models were trained on the same dataset, and each data point represents a single biological assembly from the reconstructed LigandMPNN test set.

**Fig. S8.**
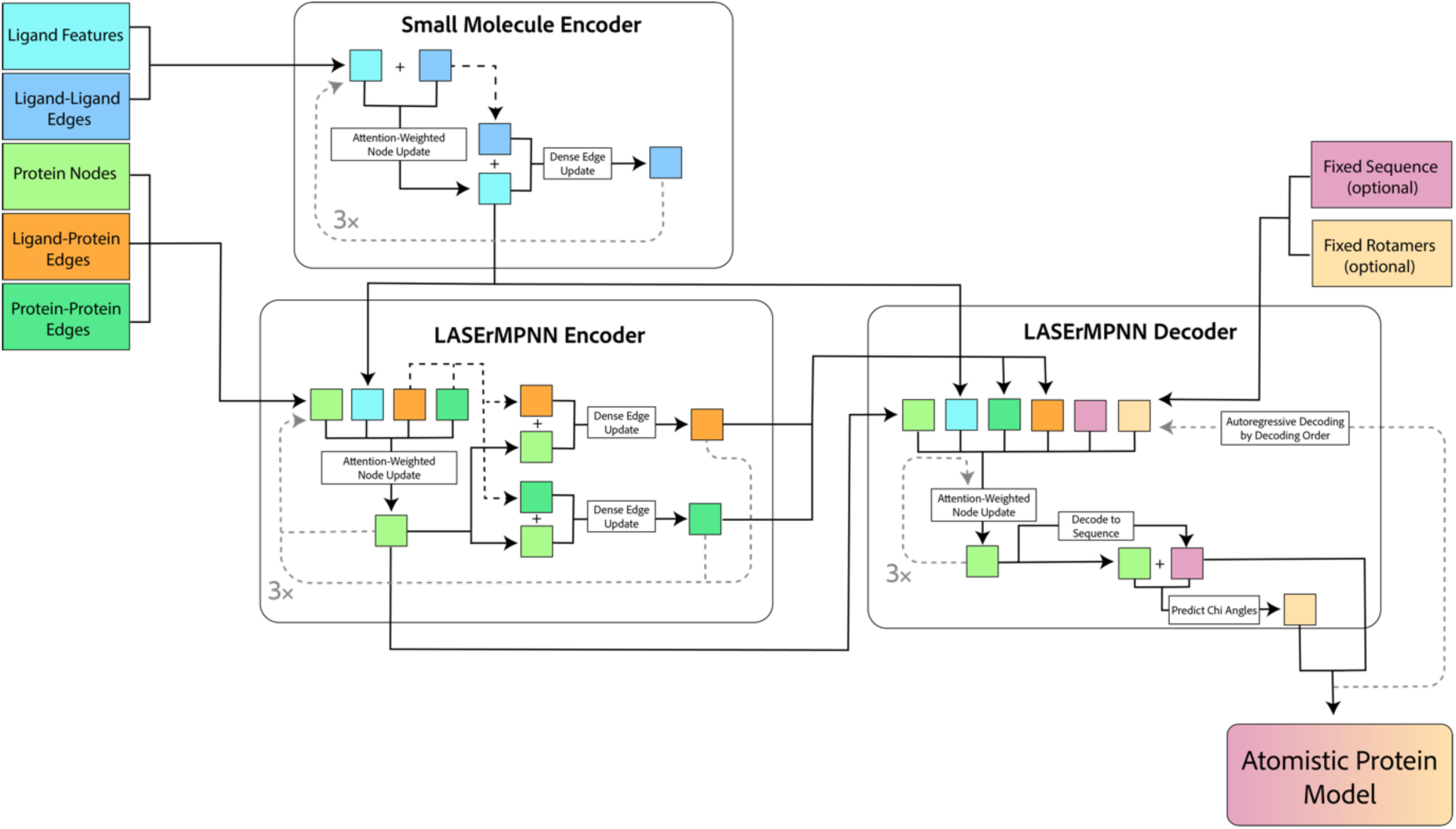
Block diagram of information flow through the LASErMPNN architecture. The LASErMPNN small-molecule encoder is a distinct pre-trainable module used to generate per-atom embeddings of each ligand atom from the input ligand cartesian coordinates and atomic numbers. These embeddings are frozen and used as input to the LASErMPNN Encoder and Decoder modules, which use a heterograph message passing formulation to build up embeddings for protein nodes, which are initialized to zeros from edge distance information. These embeddings are decoded to both a choice of amino acid identity and sidechain dihedral angles.

**Fig. S9.**
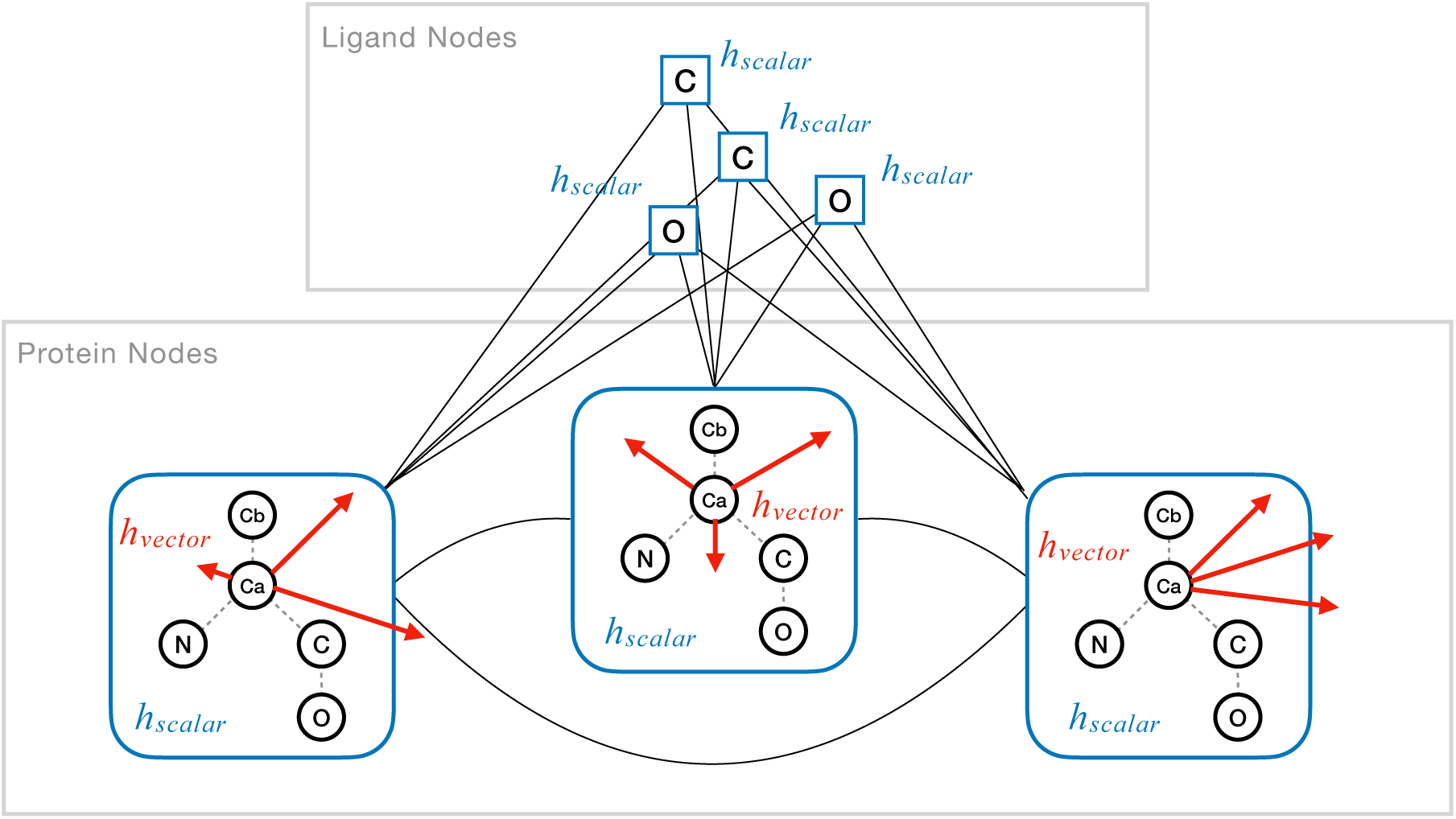
Protein nodes store equivariant vector representations. Geometric vector perceptron (GVP) layers are used to learn both scalar and vector representations to encode residue orientation on all protein nodes in the LASErMPNN architecture. The vector orientations are initialized using the normalized displacement vectors pointing from the C⍺ carbon to the N, Cβ, C, and O atoms within each frame and are updated by each encoder and decoder layer. Ligand nodes store only scalar representations. Edges denoted as black lines store a scalar edge embedding *e_ij_* which are initialized with RBF encoded distances between connected nodes.

**Fig. S10.**
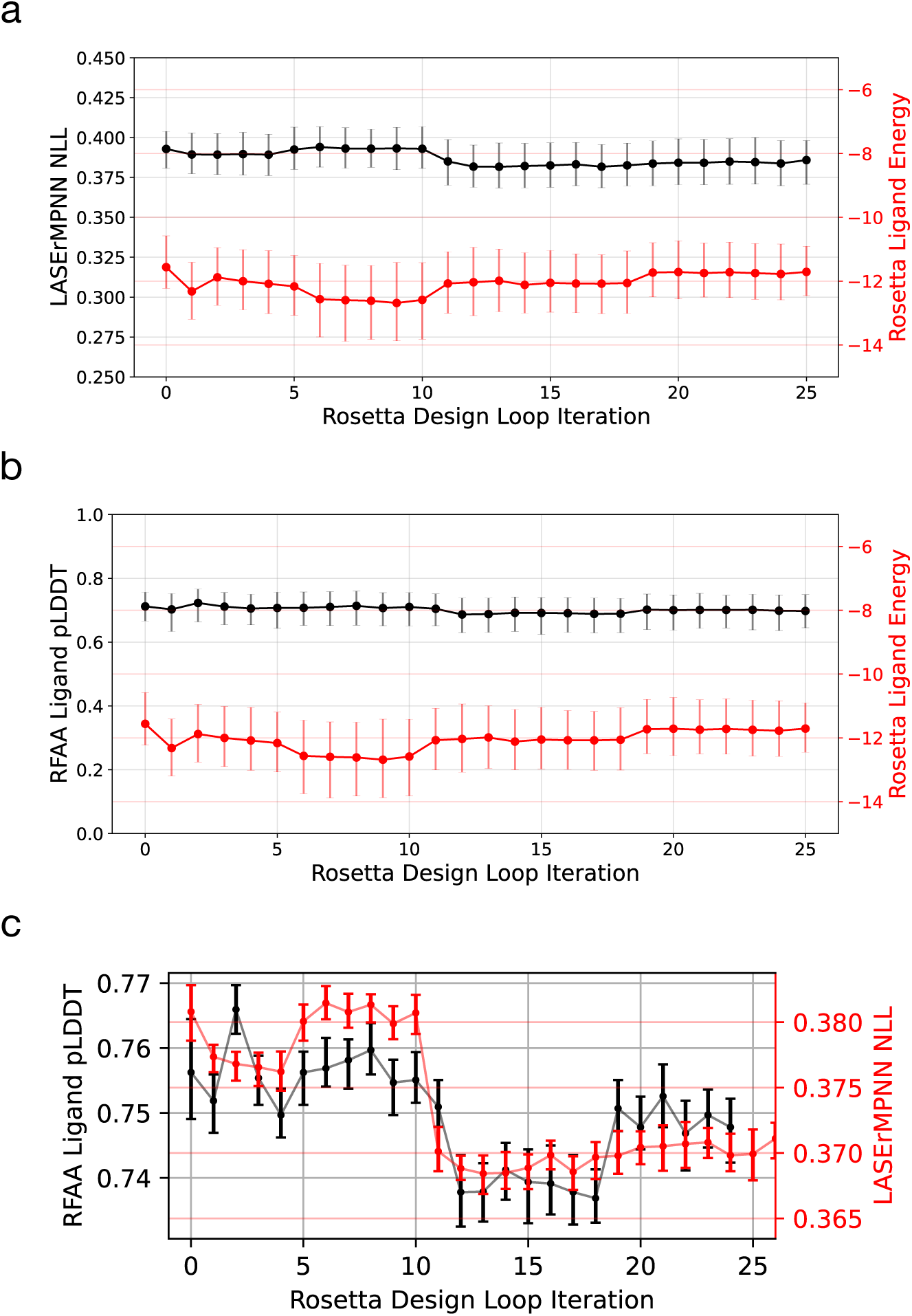
An energy-based iterative selection–expansion protocol using Rosetta does not optimize neural network confidence. We performed iterative selection–expansion starting from the same input backbone and exatecan location as in Fig. 3c. We swapped out RFAA for energy minimization (of protein and ligand coordinates) with Rosetta (Ref15 energy function). **a, b,** Median values of LASErMPNN negative log likelihood (NLL) of designed sequences, ligand pLDDT from RFAA, and Rosetta ligand energy with error bars displaying the first and third quartiles for each iteration. **c,** Data points show the value of the third quartile for sampled pLDDT and first quartile for NLL, with error bars representing 95% confidence intervals for these values computed by bootstrapping (N=100,000) the sampled data for each iteration.

**Fig. S11.**
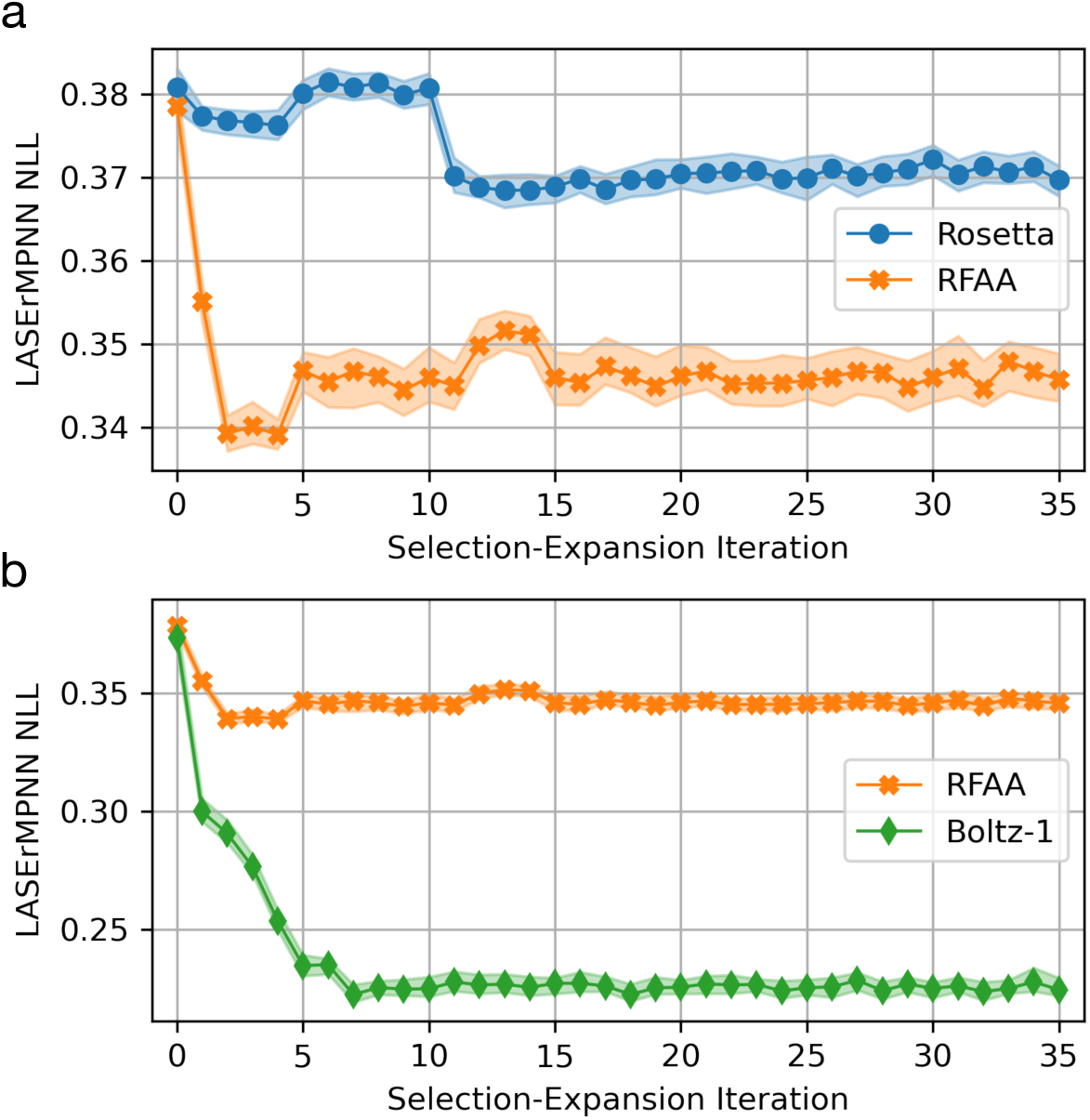
Neural networks for protein–ligand co-structure prediction are more efficient than Rosetta at optimizing LASErMPNN confidence. **a,** Comparison of LASErMPNN’s negative log likelihood (NLL) of designed sequences of a selection–expansion algorithm, which selects the top three poses per iteration by minimum Rosetta ligand energy (flexible protein backbone and ligand trajectory) *vs* a selection-expansion algorithm which maximizes ligand pLDDT from RFAA **b,** Comparison of LASErMPNN’s NLL of designed sequences of a selection–expansion algorithm, which selects the top three poses per iteration by maximum ligand pLDDT from RFAA *vs* maximum ligand pLDDT from Boltz-1. All datapoints show the value of the first quartile of LASErMPNN NLL for each selection–expansion trajectory, and shaded regions show 99% confidence intervals on the value of the first quartile computed by bootstrapping (N=100,000) the sampled data for each iteration.

**Table E1.** LASErMPNN Ablation Studies*.

| Table E1. LASerMPNN Ablation Studies* |  |  |  |  |
| --- | --- | --- | --- | --- |
|  | Sequence Recovery (Overall %) | Sequence Recovery (Binding Site %) | BLOSUM62 Score (Overall Incorrect) | BLOSUM62 Score (Binding Site Incorrect) |
| LASerMPNN (baseline) | 51.7 ± 0.2 | 60.3 ± 0.3 | -0.108 ± 0.006 | 0.036 ± 0.016 |
| LASerMPNN (add edge angle features) | **52.5 ± 0.2** | **61.2 ± 0.3** | -0.117 ± 0.006 | **<u>0.143 ± 0.016</u>** |
| Ablate MSA loss | 52.3 ± 0.2 | 61.1 ± 0.3 | **<u>-0.097 ± 0.006</u>** | 0.098 ± 0.017 |
| Ablate MSA loss & chi angle decoding | 51.3 ± 0.2 | 56.8 ± 0.3 | -0.129 ± 0.007 | -0.113 ± 0.018 |
| Unfreeze pretrained ligand encoder | 52.1 ± 0.2 | 55.0 ± 0.3 | -0.137 ± 0.006 | -0.085 ± 0.016 |
| Ablate ligand encoder pretraining | 51.6 ± 0.2 | 53.2 ± 0.3 | -0.131 ± 0.006 | -0.116 ± 0.015 |
| Ablate vector representations | 51.3 ± 0.2 | 57.6 ± 0.3 | -0.115 ± 0.006 | -0.03 ± 0.015 |
| LigandMPNN (retrained) | 52.1 ± 0.2 | 56.5 ± 0.3 | -0.125 ± 0.006 | -0.099 ± 0.017 |
| LigandMPNN (author weights) | <u>53.1 ± 0.2</u> | <u>62.5 ± 0.3</u> | -0.147 ± 0.006 | 0.036 ± 0.016 |
| * The best values achieved over our reimplementation of the LigandMPNN test set are highlighted in bold, and the best values across all models are underlined. All values are means and standard deviations computed by bootstrapping (N=100,000) the value of the mean sequence recovery or BLOSUM62 score uniformly over the 5 sequences generated for each of the 317 members of the test set. Data for each ablation was generated by a single model. Variance due to randomness in training initializations and optimization is not quantified (due to compute expense). |  |  |  |  |

**Table S1.** NISE generated exatecan binder designs.

| Table S1. NISE generated exatecan binder designs. |  |  |  |  |  |  |  |  |
| --- | --- | --- | --- | --- | --- | --- | --- | --- |
| Design | Ligand pLDDT | Protein pLDDT | Ligand RMSD | Protein RMSD | Rosetta PackStat | # Ligand BUNs | BUNs Score | K<sub>d</sub> |
| NISE 2 | 0.908 | 0.902 | 0.272 | 0.280 | 0.641 | 2.0 | 23.0 | 9.7 μM |
| NISE 3 | 0.929 | 0.903 | 0.626 | 0.177 | 0.593 | 3.0 | 17.0 | 3.2 μM |
| NISE 4 | 0.859 | 0.875 | 0.672 | 0.959 | 0.596 | 3.0 | 28.0 | 16.8 μM |
| EPIC | 0.911 | 0.896 | 0.440 | 0.726 | 0.649 | 4.0 | 16.0 | 0.12 μM |

**Table S2.**
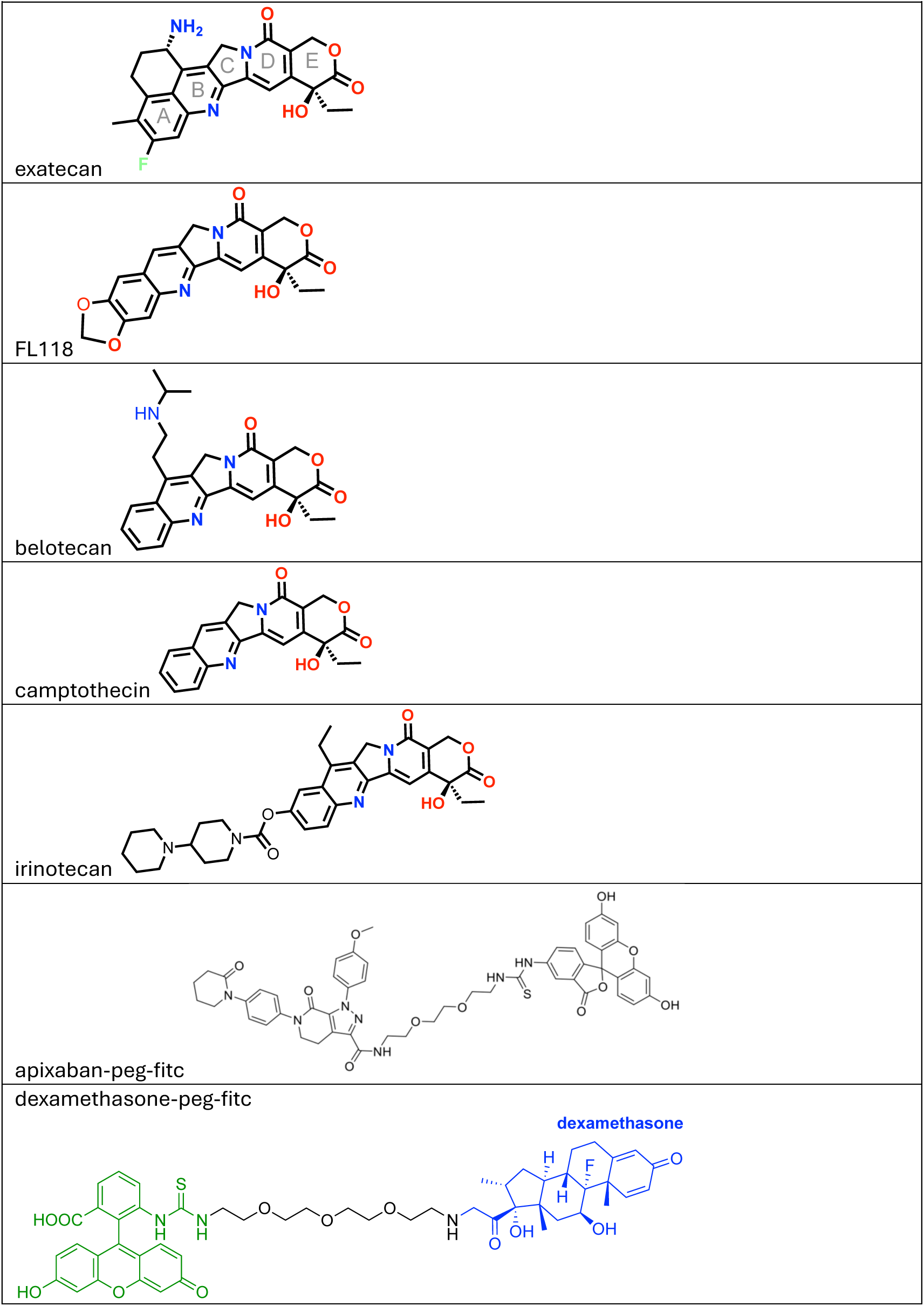
Chemical structures of ligands used in binding experiments.

| Table S2. Chemical structures of ligands used in binding experiments |
| --- |
| exatecan |
| FL118 |
| belotecan |
| camptothecin |
| irinotecan |
| apixaban-peg-fitc |
| dexamethasone-peg-fitc |

**Table S3.**
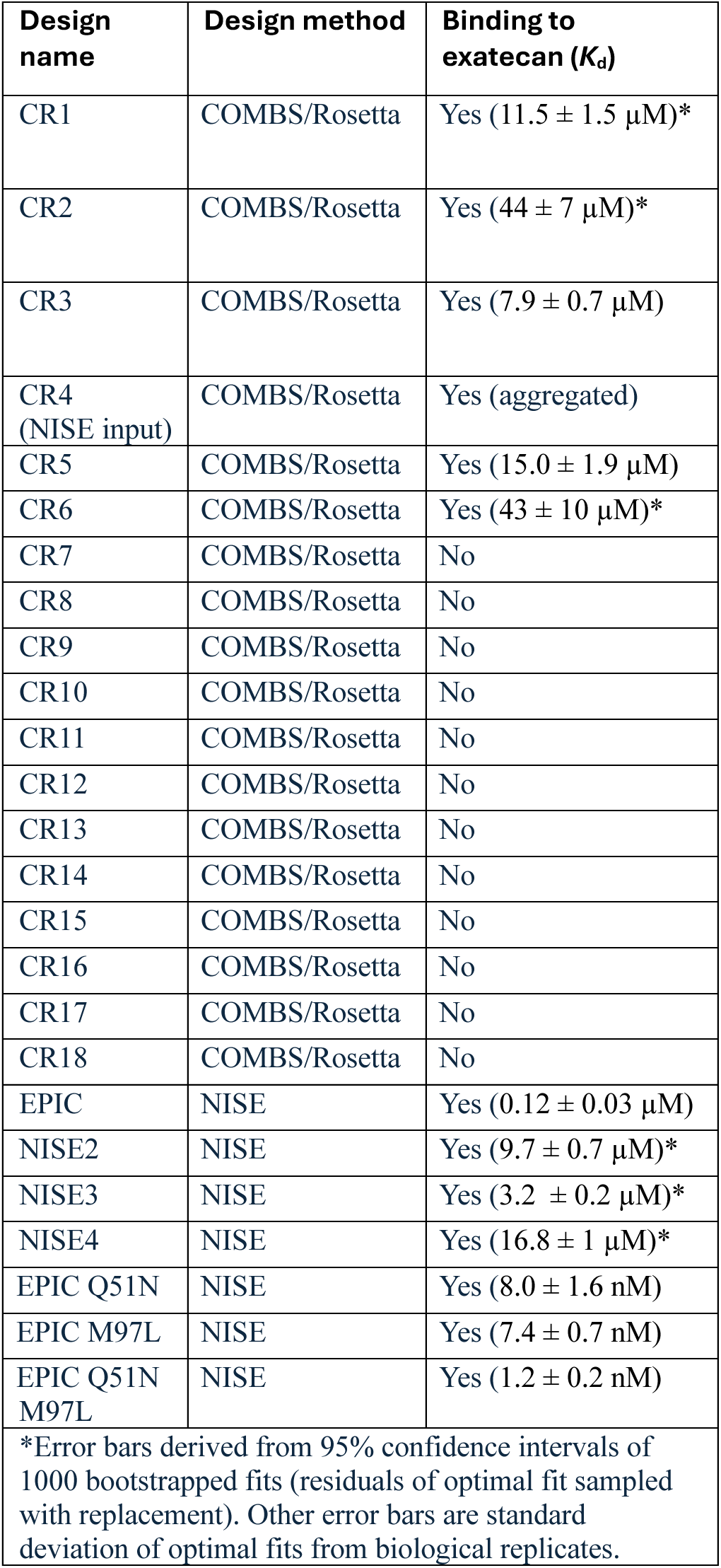
Characterized protein sequences.

| **Table S3. Characterized protein sequences** |  |  |
| --- | --- | --- |
| **Design name** | **Design method** | **Binding to exatecan ($K_d$)** |
| CR1 | COMBS/Rosetta | Yes ($11.5 \pm 1.5 \mu\text{M}$)* |
| CR2 | COMBS/Rosetta | Yes ($44 \pm 7 \mu\text{M}$)* |
| CR3 | COMBS/Rosetta | Yes ($7.9 \pm 0.7 \mu\text{M}$) |
| CR4 (NISE input) | COMBS/Rosetta | Yes (aggregated) |
| CR5 | COMBS/Rosetta | Yes ($15.0 \pm 1.9 \mu\text{M}$) |
| CR6 | COMBS/Rosetta | Yes ($43 \pm 10 \mu\text{M}$)* |
| CR7 | COMBS/Rosetta | No |
| CR8 | COMBS/Rosetta | No |
| CR9 | COMBS/Rosetta | No |
| CR10 | COMBS/Rosetta | No |
| CR11 | COMBS/Rosetta | No |
| CR12 | COMBS/Rosetta | No |
| CR13 | COMBS/Rosetta | No |
| CR14 | COMBS/Rosetta | No |
| CR15 | COMBS/Rosetta | No |
| CR16 | COMBS/Rosetta | No |
| CR17 | COMBS/Rosetta | No |
| CR18 | COMBS/Rosetta | No |
| EPIC | NISE | Yes ($0.12 \pm 0.03 \mu\text{M}$) |
| NISE2 | NISE | Yes ($9.7 \pm 0.7 \mu\text{M}$)* |
| NISE3 | NISE | Yes ($3.2 \pm 0.2 \mu\text{M}$)* |
| NISE4 | NISE | Yes ($16.8 \pm 1 \mu\text{M}$)* |
| EPIC Q51N | NISE | Yes ($8.0 \pm 1.6 \text{nM}$) |
| EPIC M97L | NISE | Yes ($7.4 \pm 0.7 \text{nM}$) |
| EPIC Q51N M97L | NISE | Yes ($1.2 \pm 0.2 \text{nM}$) |
| *Error bars derived from 95% confidence intervals of 1000 bootstrapped fits (residuals of optimal fit sampled with replacement). Other error bars are standard deviation of optimal fits from biological replicates. |  |  |

**Table S4.** Data collection and refinement statistics for EPIC.

| **Table S4. Data collection and refinement statistics for EPIC** |  |
| --- | --- |
| Wavelength | 0.9201 |
| Resolution range | 31.22 - 1.96 (2.03 - 1.96) |
| Space group | P 21 21 21 |
| Unit cell | 91.027 93.683 94.328 90 90 90 |
| Total reflections | 363998 (39395) |
| Unique reflections | 55943 (5743) |
| Multiplicity | 6.5 (6.9) |
| Completeness (%) | 97.34 (99.86) |
| Mean I/sigma(I) | 14.02 (1.75) |
| Wilson B-factor | 30.28 |
| R-merge | 0.09092 (1.142) |
| R-meas | 0.09896 (1.236) |
| R-pim | 0.03853 (0.469) |
| CC1/2 | 0.999 (0.527) |
| CC* | 1 (0.831) |
| Reflections used in refinement | 56994 (5742) |
| Reflections used for R-free | 2756 (309) |
| R-work | 0.2496 (0.3360) |
| R-free | 0.2853 (0.3697) |
| CC(work) | 0.938 (0.660) |
| CC(free) | 0.894 (0.509) |
| Number of non-hydrogen atoms | 5057 |
| macromolecules | 4657 |
| ligands | 324 |
| solvent | 214 |
| Protein residues | 592 |
| RMS(bonds) | 0.007 |
| RMS(angles) | 1.31 |
| Ramachandran favored (%) | 99.49 |
| Ramachandran allowed (%) | 0.51 |
| Ramachandran outliers (%) | 0.00 |
| Rotamer outliers (%) | 4.38 |
| Clashscore | 7.54 |
| Average B-factor | 40.40 |
| macromolecules | 40.92 |
| ligands | 32.40 |
| solvent | 35.91 |
| Number of TLS groups | 1 |
| Statistics for the highest-resolution shell are shown in parentheses. |  |

**Table S5.** Data collection and refinement statistics for EPIC^Q51N^.

| **Table S5. Data collection and refinement statistics for EPIC<sup>Q51N</sup>** |  |
| --- | --- |
| Wavelength | 0.9201 |
| Resolution range | 44.45 - 2.214 (2.294 - 2.214) |
| Space group | I 21 21 21 |
| Unit cell | 55.182 66.003 177.789 90 90 90 |
| Total reflections | 76617 (7650) |
| Unique reflections | 16539 (1204) |
| Multiplicity | 4.6 (4.7) |
| Completeness (%) | 97.13 (74.14) |
| Mean I/sigma(I) | 6.83 (0.94) |
| Wilson B-factor | 33.85 |
| R-merge | 0.1782 (2.683) |
| R-meas | 0.2015 (3.023) |
| R-pim | 0.09203 (1.372) |
| CC1/2 | 0.997 (0.443) |
| CC* | 0.999 (0.783) |
| Reflections used in refinement | 16154 (1204) |
| Reflections used for R-free | 801 (61) |
| R-work | 0.2746 (0.3707) |
| R-free | 0.3245 (0.3808) |
| CC(work) | 0.881 (0.459) |
| CC(free) | 0.940 (0.484) |
| Number of non-hydrogen atoms | 2414 |
| macromolecules | 2305 |
| ligands | 110 |
| solvent | 45 |
| Protein residues | 296 |
| RMS(bonds) | 0.002 |
| RMS(angles) | 0.63 |
| Ramachandran favored (%) | 98.97 |
| Ramachandran allowed (%) | 0.68 |
| Ramachandran outliers (%) | 0.34 |
| Rotamer outliers (%) | 1.27 |
| Clashscore | 2.70 |
| Average B-factor | 48.75 |
| macromolecules | 49.09 |
| ligands | 38.28 |
| solvent | 46.21 |
| Number of TLS groups | 1 |
| Statistics for the highest-resolution shell are shown in parentheses. |  |

## References

1. An, L., Said, M., Tran, L., Majumder, S., Goreshnik, I., Lee, G. R., Juergens, D., Dauparas, J., Anishchenko, I., Coventry, B., Bera, A. K., Kang, A., Levine, P. M., Alvarez, V., Pillai, A., Norn, C., Feldman, D., Zorine, D., Hicks, D. R., Li, X., Sanchez, M. G., Vafeados, D. K., Salveson, P. J., Vorobieva, A. A. & Baker, D. Binding and sensing diverse small molecules using shape-complementary pseudocycles. Science 385, 276–282 (2024).

2. Lee, G. R., Pellock, S. J., Norn, C., Tischer, D., Dauparas, J., Anishchenko, I., Mercer, J. A. M., Kang, A., Bera, A., Nguyen, H., Goreshnik, I., Vafeados, D., Roullier, N., Han, H. L., Coventry, B., Haddox, H. K., Liu, D. R., Yeh, A. H.-W. & Baker, D. Small-molecule binding and sensing with a designed protein family. bioRxiv 2023.11.01.565201 (2023) doi:10.1101/2023.11.01.565201.

3. Kortemme, T. De novo protein design-From new structures to programmable functions. Cell 187, 526–544 (2024).

4. Dou, J., Vorobieva, A. A., Sheffler, W., Doyle, L. A., Park, H., Bick, M. J., Mao, B., Foight, G. W., Lee, M. Y., Gagnon, L. A., Carter, L., Sankaran, B., Ovchinnikov, S., Marcos, E., Huang, P.-S., Vaughan, J. C., Stoddard, B. L. & Baker, D. De novo design of a fluorescence-activating β-barrel. Nature 561, 485–491 (2018).

5. Lu, L., Gou, X., Tan, S. K., Mann, S. I., Yang, H., Zhong, X., Gazgalis, D., Valdiviezo, J., Jo, H., Wu, Y., Diolaiti, M. E., Ashworth, A., Polizzi, N. F. & DeGrado, W. F. De novo design of drug-binding proteins with predictable binding energy and specificity. Science 384, 106–112 (2024).

6. Polizzi, N. F. & DeGrado, W. F. A defined structural unit enables de novo design of small-molecule-binding proteins. Science 369, 1227–1233 (2020).

7. Chen, Y., Bhattacharya, S., Bergmann, L., Correy, G. J., Tan, S., Hou, K., Biel, J., Lu, L., Bakanas, I., Polizzi, N. F., Fraser, J. S. & DeGrado, W. F. Emergence of specific binding and catalysis from a designed generalist binding protein. bioRxiv 2025.01.30.635804 (2025) doi:10.1101/2025.01.30.635804.

8. Krishna, R., Wang, J., Ahern, W., Sturmfels, P., Venkatesh, P., Kalvet, I., Lee, G. R., Morey-Burrows, F. S., Anishchenko, I., Humphreys, I. R., McHugh, R., Vafeados, D., Li, X., Sutherland, G. A., Hitchcock, A., Hunter, C. N., Kang, A., Brackenbrough, E., Bera, A. K., Baek, M., DiMaio, F. & Baker, D. Generalized biomolecular modeling and design with RoseTTAFold All-Atom. Science 384, eadl2528 (2024).

9. Minami, H., Fujii, H., Igarashi, T., Itoh, K., Tamanoi, K., Oguma, T. & Sasaki, Y. Phase I and pharmacological study of a new camptothecin derivative, exatecan mesylate (DX-8951f), infused over 30 minutes every three weeks. Clin. Cancer Res. 7, 3056–3064 (2001).

10. Marchand, A., Buckley, S., Schneuing, A., Pacesa, M., Elia, M., Gainza, P., Elizarova, E., Neeser, R. M., Lee, P.-W., Reymond, L., Miao, Y., Scheller, L., Georgeon, S., Schmidt, J., Schwaller, P., Maerkl, S. J., Bronstein, M. & Correia, B. E. Targeting protein–ligand neosurfaces with a generalizable deep learning tool. Nature 639, 522–531 (2025).

11. Noske, J., Kynast, J. P., Lemm, D., Schmidt, S. & Höcker, B. PocketOptimizer 2.0: A modular framework for computer-aided ligand-binding design. Protein Sci. 32, e4516 (2023).

12. Glasgow, A. A., Huang, Y.-M., Mandell, D. J., Thompson, M., Ritterson, R., Loshbaugh, A. L., Pellegrino, J., Krivacic, C., Pache, R. A., Barlow, K. A., Ollikainen, N., Jeon, D., Kelly, M. J. S., Fraser, J. S. & Kortemme, T. Computational design of a modular protein sense-response system. Science 366, 1024–1028 (2019).

13. Leonard, A. C., Friedman, A. J., Chayer, R., Petersen, B. M., Woojuh, J., Xing, Z., Cutler, S. R., Kaar, J. L., Shirts, M. R. & Whitehead, T. A. Rationalizing diverse binding mechanisms to the same protein fold: Insights for ligand recognition and biosensor design. ACS Chem. Biol. 19, 1757–1772 (2024).

14. Zhu, J., Liang, M., Sun, K., Wei, Y., Guo, R., Zhang, L., Shi, J., Ma, D., Hu, Q., Huang, G. & Lu, P. De novo design of transmembrane fluorescence-activating proteins. Nature 1–9 (2025).

15. Watson, J. L., Juergens, D., Bennett, N. R., Trippe, B. L., Yim, J., Eisenach, H. E., Ahern, W., Borst, A. J., Ragotte, R. J., Milles, L. F., Wicky, B. I. M., Hanikel, N., Pellock, S. J., Courbet, A., Sheffler, W., Wang, J., Venkatesh, P., Sappington, I., Torres, S. V., Lauko, A., De Bortoli, V., Mathieu, E., Ovchinnikov, S., Barzilay, R., Jaakkola, T. S., DiMaio, F., Baek, M. & Baker, D. De novo design of protein structure and function with RFdiffusion. Nature 620, 1089–1100 (2023).

16. Torres, S. V., Leung, P. J. Y., Venkatesh, P., Lutz, I. D., Hink, F., Huynh, H.-H., Becker, J., Yeh, A. H.-W., Juergens, D., Bennett, N. R., Hoofnagle, A. N., Huang, E., MacCoss, M. J., Expòsit, M., Lee, G. R., Bera, A. K., Kang, A., De La Cruz, J., Levine, P. M., Li, X., Lamb, M., Gerben, S. R., Murray, A., Heine, P., Korkmaz, E. N., Nivala, J., Stewart, L., Watson, J. L., Rogers, J. M. & Baker, D. De novo design of high-affinity binders of bioactive helical peptides. Nature 1–3 (2023).

17. Vázquez Torres, S., Benard Valle, M., Mackessy, S. P., Menzies, S. K., Casewell, N. R., Ahmadi, S., Burlet, N. J., Muratspahić, E., Sappington, I., Overath, M. D., Rivera-de-Torre, E., Ledergerber, J., Laustsen, A. H., Boddum, K., Bera, A. K., Kang, A., Brackenbrough, E., Cardoso, I. A., Crittenden, E. P., Edge, R. J., Decarreau, J., Ragotte, R. J., Pillai, A. S., Abedi, M., Han, H. L., Gerben, S. R., Murray, A., Skotheim, R., Stuart, L., Stewart, L., Fryer, T. J. A., Jenkins, T. P. & Baker, D. De novo designed proteins neutralize lethal snake venom toxins. Nature 1–7 (2025).

18. Bennett, N. R., Watson, J. L., Ragotte, R. J., Borst, A. J., See, D. L., Weidle, C., Biswas, R., Yu, Y., Shrock, E. L., Ault, R., Leung, P. J. Y., Huang, B., Goreshnik, I., Tam, J., Carr, K. D., Singer, B., Criswell, C., Wicky, B. I. M., Vafeados, D., Sanchez, M. G., Kim, H. M., Vázquez Torres, S., Chan, S., Sun, S. M., Spear, T., Sun, Y., O’Reilly, K., Maris, J. M., Sgourakis, N. G., Melnyk, R. A., Liu, C. C. & Baker, D. Atomically accurate de novo design of antibodies with RFdiffusion. bioRxiv 2024.03.14.585103 (2025) doi:10.1101/2024.03.14.585103.

19. Pacesa, M., Nickel, L., Schellhaas, C., Schmidt, J., Pyatova, E., Kissling, L., Barendse, P., Choudhury, J., Kapoor, S., Alcaraz-Serna, A., Cho, Y., Ghamary, K. H., Vinué, L., Yachnin, B. J., Wollacott, A. M., Buckley, S., Westphal, A. H., Lindhoud, S., Georgeon, S., Goverde, C. A., Hatzopoulos, G. N., Gönczy, P., Muller, Y. D., Schwank, G., Swarts, D. C., Vecchio, A. J., Schneider, B. L., Ovchinnikov, S. & Correia, B. E. BindCraft: one-shot design of functional protein binders. bioRxiv 2024.09.30.615802 (2024) doi:10.1101/2024.09.30.615802.

20. Eguchi, R. R., Choe, C. A. & Huang, P.-S. Ig-VAE: Generative modeling of protein structure by direct 3D coordinate generation. PLoS Comput. Biol. 18, e1010271 (2022).

21. Bhat, S., Palepu, K., Hong, L., Mao, J., Ye, T., Iyer, R., Zhao, L., Chen, T., Vincoff, S., Watson, R., Wang, T. Z., Srijay, D., Kavirayuni, V. S., Kholina, K., Goel, S., Vure, P., Deshpande, A. J., Soderling, S. H., DeLisa, M. P. & Chatterjee, P. De novo design of peptide binders to conformationally diverse targets with contrastive language modeling. Sci. Adv. 11, eadr8638 (2025).

22. Shuai, R. W., Ruffolo, J. A. & Gray, J. J. IgLM: Infilling language modeling for antibody sequence design. Cell Syst. 14, 979–989.e4 (2023).

23. Zambaldi, V., La, D., Chu, A. E., Patani, H., Danson, A. E., Kwan, T. O. C., Frerix, T., Schneider, R. G., Saxton, D., Thillaisundaram, A., Wu, Z., Moraes, I., Lange, O., Papa, E., Stanton, G., Martin, V., Singh, S., Wong, L. H., Bates, R., Kohl, S. A., Abramson, J., Senior, A. W., Alguel, Y., Wu, M. Y., Aspalter, I. M., Bentley, K., Bauer, D. L. V., Cherepanov, P., Hassabis, D., Kohli, P., Fergus, R. & Wang, J. De novo design of high-affinity protein binders with AlphaProteo. bioRxiv (2024).

24. Ingraham, J. B., Baranov, M., Costello, Z., Barber, K. W., Wang, W., Ismail, A., Frappier, V., Lord, D. M., Ng-Thow-Hing, C., Van Vlack, E. R., Tie, S., Xue, V., Cowles, S. C., Leung, A., Rodrigues, J. V., Morales-Perez, C. L., Ayoub, A. M., Green, R., Puentes, K., Oplinger, F., Panwar, N. V., Obermeyer, F., Root, A. R., Beam, A. L., Poelwijk, F. J. & Grigoryan, G. Illuminating protein space with a programmable generative model. Nature 623, 1070–1078 (2023).

25. Dauparas, J., Lee, G. R., Pecoraro, R., An, L., Anishchenko, I., Glasscock, C. & Baker, D. Atomic context-conditioned protein sequence design using LigandMPNN. Nat. Methods 22, 717–723 (2025).

26. Tinberg, C. E., Khare, S. D., Dou, J., Doyle, L., Nelson, J. W., Schena, A., Jankowski, W., Kalodimos, C. G., Johnsson, K., Stoddard, B. L. & Baker, D. Computational design of ligand-binding proteins with high affinity and selectivity. Nature 501, 212–216 (2013).

27. Polizzi, N. F., Wu, Y., Lemmin, T., Maxwell, A. M., Zhang, S.-Q., Rawson, J., Beratan, D. N., Therien, M. J. & DeGrado, W. F. De novo design of a hyperstable non-natural protein-ligand complex with sub-Å accuracy. Nat. Chem. 9, 1157–1164 (2017).

28. McCann, J. J., Pike, D. H., Brown, M. C., Crouse, D. T., Nanda, V. & Koder, R. L. Computational design of a sensitive, selective phase-changing sensor protein for the VX nerve agent. Science Advances 8, eabh3421 (2022).

29. Kuhlman, B., Dantas, G., Ireton, G. C., Varani, G., Stoddard, B. L. & Baker, D. Design of a novel globular protein fold with atomic-level accuracy. Science 302, 1364–1368 (2003).

30. Dauparas, J., Anishchenko, I., Bennett, N., Bai, H., Ragotte, R. J., Milles, L. F., Wicky, B. I. M., Courbet, A., de Haas, R. J., Bethel, N., Leung, P. J. Y., Huddy, T. F., Pellock, S., Tischer, D., Chan, F., Koepnick, B., Nguyen, H., Kang, A., Sankaran, B., Bera, A. K., King, N. P. & Baker, D. Robust deep learning-based protein sequence design using ProteinMPNN. Science 378, 49–56 (2022).

31. Jumper, J., Evans, R., Pritzel, A., Green, T., Figurnov, M., Ronneberger, O., Tunyasuvunakool, K., Bates, R., Žídek, A., Potapenko, A., Bridgland, A., Meyer, C., Kohl, S. A. A., Ballard, A. J., Cowie, A., Romera-Paredes, B., Nikolov, S., Jain, R., Adler, J., Back, T., Petersen, S., Reiman, D., Clancy, E., Zielinski, M., Steinegger, M., Pacholska, M., Berghammer, T., Bodenstein, S., Silver, D., Vinyals, O., Senior, A. W., Kavukcuoglu, K., Kohli, P. & Hassabis, D. Highly accurate protein structure prediction with AlphaFold. Nature 596, 583–589 (2021).

32. Goudy, O. J., Nallathambi, A., Kinjo, T., Randolph, N. Z. & Kuhlman, B. In silico evolution of autoinhibitory domains for a PD-L1 antagonist using deep learning models. Proc. Natl. Acad. Sci. U. S. A. 120, e2307371120 (2023).

33. Wohlwend, J., Corso, G., Passaro, S., Reveiz, M., Leidal, K., Swiderski, W., Portnoi, T., Chinn, I., Silterra, J., Jaakkola, T. & Barzilay, R. Boltz-1 democratizing biomolecular interaction modeling. bioRxiv 2024.11.19.624167 (2024) doi:10.1101/2024.11.19.624167.

34. Abramson, J., Adler, J., Dunger, J., Evans, R., Green, T., Pritzel, A., Ronneberger, O., Willmore, L., Ballard, A. J., Bambrick, J., Bodenstein, S. W., Evans, D. A., Hung, C.-C., O’Neill, M., Reiman, D., Tunyasuvunakool, K., Wu, Z., Žemgulytė, A., Arvaniti, E., Beattie, C., Bertolli, O., Bridgland, A., Cherepanov, A., Congreve, M., Cowen-Rivers, A. I., Cowie, A., Figurnov, M., Fuchs, F. B., Gladman, H., Jain, R., Khan, Y. A., Low, C. M. R., Perlin, K., Potapenko, A., Savy, P., Singh, S., Stecula, A., Thillaisundaram, A., Tong, C., Yakneen, S., Zhong, E. D., Zielinski, M., Žídek, A., Bapst, V., Kohli, P., Jaderberg, M., Hassabis, D. & Jumper, J. M. Accurate structure prediction of biomolecular interactions with AlphaFold 3. Nature 630, 493–500 (2024).

35. Besag, J. On the Statistical Analysis of Dirty Pictures. Journal of the Royal Statistical Society. Series B (Methodological) 48, 259–302 (1986).

36. Eastman, P., Pritchard, B. P., Chodera, J. D. & Markland, T. E. Nutmeg and SPICE: Models and data for biomolecular machine learning. J. Chem. Theory Comput. 20, 8583–8593 (2024).

37. Eastman, P., Behara, P. K., Dotson, D. L., Galvelis, R., Herr, J. E., Horton, J. T., Mao, Y., Chodera, J. D., Pritchard, B. P., Wang, Y., De Fabritiis, G. & Markland, T. E. SPICE, A dataset of drug-like molecules and peptides for training machine learning potentials. Sci. Data 10, 11 (2023).

38. Fu, Z., Li, S., Han, S., Shi, C. & Zhang, Y. Antibody drug conjugate: the “biological missile” for targeted cancer therapy. Signal Transduct. Target. Ther. 7, 93 (2022).

39. Li, W., Veale, K. H., Qiu, Q., Sinkevicius, K. W., Maloney, E. K., Costoplus, J. A., Lau, J., Evans, H. L., Setiady, Y., Ab, O., Abbott, S. M., Lee, J., Wisitpitthaya, S., Skaletskaya, A., Wang, L., Keating, T. A., Chari, R. V. J. & Widdison, W. C. Synthesis and evaluation of camptothecin antibody-drug conjugates. ACS Med. Chem. Lett. 10, 1386–1392 (2019).

40. Grigoryan, G. & DeGrado, W. F. Probing designability via a generalized model of helical bundle geometry. J. Mol. Biol. 405, 1079–1100 (2011).

41. Zhou, J. & Grigoryan, G. Rapid search for tertiary fragments reveals protein sequence-structure relationships. Protein Sci. 24, 508–524 (2015).

42. Leman, J. K. et al. Macromolecular modeling and design in Rosetta: recent methods and frameworks. Nat. Methods 17, 665–680 (2020).

43. Bryant, P. & Elofsson, A. Peptide binder design with inverse folding and protein structure prediction. Commun. Chem. 6, 229 (2023).

44. Wei, J., Wang, X., Schuurmans, D., Bosma, M., Ichter, B., Xia, F., Chi, E., Le, Q. & Zhou, D. Chain-of-thought prompting elicits reasoning in large language models. arXiv [cs.CL] (2022).

45. Hayes, T., Rao, R., Akin, H., Sofroniew, N. J., Oktay, D., Lin, Z., Verkuil, R., Tran, V. Q., Deaton, J., Wiggert, M., Badkundri, R., Shafkat, I., Gong, J., Derry, A., Molina, R. S., Thomas, N., Khan, Y., Mishra, C., Kim, C., Bartie, L. J., Nemeth, M., Hsu, P. D., Sercu, T., Candido, S. & Rives, A. Simulating 500 million years of evolution with a language model. bioRxiv 2024.07.01.600583 (2024) doi:10.1101/2024.07.01.600583.

46. Eastman, P., Galvelis, R., Peláez, R. P., Abreu, C. R. A., Farr, S. E., Gallicchio, E., Gorenko, A., Henry, M. M., Hu, F., Huang, J., Krämer, A., Michel, J., Mitchell, J. A., Pande, V. S., Rodrigues, J. P., Rodriguez-Guerra, J., Simmonett, A. C., Singh, S., Swails, J., Turner, P., Wang, Y., Zhang, I., Chodera, J. D., De Fabritiis, G. & Markland, T. E. OpenMM 8: Molecular dynamics simulation with machine learning potentials. J. Phys. Chem. B 128, 109–116 (2024).

47. Cho, Y., Pacesa, M., Zhang, Z., Correia, B. & Ovchinnikov, S. BoltzDesign1: Inverting all-atom structure prediction model for generalized biomolecular binder design. bioRxiv 2025.04.06.647261 (2025) doi:10.1101/2025.04.06.647261.

48. Landrum, G., Tosco, P., Kelley, B., Rodriguez, R., Cosgrove, D., Vianello, R., Gedeck, P., Jones, G., NadineSchneider, Kawashima, E., Nealschneider, D., Dalke, A., Swain, M., Cole, B., Turk, S., Savelev, A., tadhurst-cdd, Vaucher, A., Wójcikowski, M., Take, I., Walker, R., Scalfani, V. F., Faara, H., Ujihara, K., Probst, D., Lehtivarjo, J., Godin, G., Pahl, A. & Monat, J. rdkit/rdkit: 2025_03_1 (Q1 2025) Release. Preprint at 10.5281/zenodo.15115844.

49. Brody, S., Alon, U. & Yahav, E. How Attentive are Graph Attention Networks? arXiv [cs.LG] (2021).

50. Veličković, P., Cucurull, G., Casanova, A., Romero, A., Liò, P. & Bengio, Y. Graph Attention Networks. arXiv [stat.ML] (2017).

51. Jing, B., Eismann, S., Suriana, P., Townshend, R. J. L. & Dror, R. Learning from protein structure with geometric vector perceptrons. arXiv [q-bio.BM] (2020).

52. Ahdritz, G., Bouatta, N., Kadyan, S., Jarosch, L., Berenberg, D., Fisk, I., Watkins, A. M., Ra, S., Bonneau, R. & AlQuraishi, M. OpenProteinSet: Training data for structural biology at scale. arXiv [q-bio.BM] (2023).

53. Corso, G., Deng, A., Fry, B., Polizzi, N., Barzilay, R. & Jaakkola, T. Deep confident steps to new pockets: Strategies for docking generalization. arXiv [q-bio.BM] (2024).

54. Word, J. M., Lovell, S. C., Richardson, J. S. & Richardson, D. C. Asparagine and glutamine: using hydrogen atom contacts in the choice of side-chain amide orientation. J. Mol. Biol. 285, 1735–1747 (1999).

55. Steinegger, M. & Söding, J. MMseqs2 enables sensitive protein sequence searching for the analysis of massive data sets. Nat. Biotechnol. 35, 1026–1028 (2017).

56. Fry, B. & Polizzi, N. LASErMPNN Training Dataset (REDUCE Protonated PDB). (2025) doi:10.5281/ZENODO.15035128.

57. Fleishman, S. J., Leaver-Fay, A., Corn, J. E., Strauch, E.-M., Khare, S. D., Koga, N., Ashworth, J., Murphy, P., Richter, F., Lemmon, G., Meiler, J. & Baker, D. RosettaScripts: a scripting language interface to the Rosetta macromolecular modeling suite. PLoS One 6, e20161 (2011).

58. Polizzi, N. Designs and analysis code for Neural Iterative Selection-Expansion paper. (Zenodo, 2025). doi:10.5281/zenodo.15257832.

59. Vonrhein, C., Flensburg, C., Keller, P., Sharff, A., Smart, O., Paciorek, W., Womack, T. & Bricogne, G. Data processing and analysis with the autoPROC toolbox. Acta Crystallogr. D Biol. Crystallogr. 67, 293–302 (2011).

60. Adams, P. D., Afonine, P. V., Bunkóczi, G., Chen, V. B., Echols, N., Headd, J. J., Hung, L.-W., Jain, S., Kapral, G. J., Grosse Kunstleve, R. W., McCoy, A. J., Moriarty, N. W., Oeffner, R. D., Read, R. J., Richardson, D. C., Richardson, J. S., Terwilliger, T. C. & Zwart, P. H. The Phenix software for automated determination of macromolecular structures. Methods 55, 94–106 (2011).

61. Casañal, A., Lohkamp, B. & Emsley, P. Current developments in Coot for macromolecular model building of Electron Cryo-microscopy and Crystallographic Data. Protein Sci. 29, 1069–1078 (2020).

